# The Influence of Ecosystem and Phylogeny on Tropical Tree Crown Size and Shape

**DOI:** 10.1101/789255

**Authors:** Alexander Shenkin, Lisa Patrick Bentley, Imma Oliveras, Norma Salinas, Stephen Adu-Bredu, Ben Hur Marimon, Beatriz Marimon, Theresa Peprah, Efrain Lopez Choque, Lucio Trujillo Rodriguez, Edith Rosario Clemente Arenas, Christian Adonteng, John Seidu, Fabio Barbosa Passos, Simone Matias Reis, Benjamin Blonder, Miles Silman, Brian J. Enquist, Gregory P. Asner, Yadvinder Malhi

**Author notes:** Correspondence: Alexander Shenkin.

## Abstract

The sizes and shapes of tree crowns are of fundamental importance in ecology, yet understanding the forces that determine them remains elusive. A cardinal question facing ecologists is the degree to which general and non-specific versus ecological and context-dependent processes are responsible for shaping tree crowns. Here, we test this question for the first time across diverse tropical ecosystems. Using trees from 20 plots varying in elevation, precipitation, and ecosystem type (savanna-forest transitions) across the paleo- and neo-tropics, we test the relationship between crown dimensions and tree size. By analyzing these scaling relationships across environmental gradients, biogeographic regions, and phylogenetic distance, we extend Metabolic Scaling Theory (MST) predictions to include how local selective pressures shape variation in crown dimensions. Across all sites, we find strong agreement between mean trends and MST predictions for the scaling of crown size and shape, but large variation around the mean. While MST explained approximately half of the observed variation in tree crown dimensions, we find that local, ecosystem, and phylogenetic predictors account for the half of the residual variation. Crown scaling does not change significantly across regions, but does change across ecosystem types, where savanna tree crowns grow more quickly with tree size than forest tree crowns. Crowns of legumes were wider and larger than those of other taxa. Thus, while MST can accurately describe the central tendency of tree crown size, local ecological conditions and evolutionary history appear to modify the scaling of crown shape. Importantly, our extension of MST incorporating these differences accounts for the mechanisms driving variation in the scaling of crown dimensions across the tropics. These results are critical when scaling the function of individual trees to larger spatial scales or incorporating the size and shape of tree crowns in global biogeochemical models.

## 2 INTRODUCTION

The sizes and shapes of tree crowns are conspicuous and fundamental attributes of the organisms that comprise the structure of Earth’s forests. Despite the key roles tree crowns play in processes including growth and competition at the individual scale, and community assembly (e.g. Iwasa et al., 1985) and function (e.g. Sapijanskas et al., 2014) at the stand scale, the forces that determine their sizes and shapes are still debated.

The forms of tree crowns and their roles in tree growth and stem form have interested foresters for over a century (Larson, 1963). Ecological interest in tree crown form gained momentum in the 1970s. Architectural (Hallé et al., 1978) and ecological niche perspectives (Horn, 1971;Ashton, 1978;Givnish, 1984) were followed by considerations of mechanical constraint (King and Loucks, 1978;King, 1981) and life history strategy (e.g. Poorter et al., 2006). Ecologically, crown shape has been seen as the result of predictable, genetically-programmed branching processes on the one hand, and stochastic disturbance and competitive ones on the other (Busgen and Munch, 1929). In contrast to early focus on context-dependent ecological strategy, more recent theory, including Metabolic Scaling Theory (MST; West et al., 1997), competitive convergence (Iida et al., 2011;MacFarlane et al., 2017), and sphere packing (Taubert et al., 2015), has focused on the determinants of crown size from first principles. MST, for example, does not directly address the roles of niche or ontogeny, and competitive convergence and sphere packing models steer further away from ecological and physiological perspectives, given their absence of environmental and evolutionary factors. In this respect, these more recent perspectives might be classified as “general” theories, in contrast to “environmental” and “ecological” theories that privilege variation attributable to abiotic conditions in the former case, and species, niche, and life history strategy in the latter (e.g. Sapijanskas et al., 2014). In accordance with our classification, we refer to “general” and “ecological” classes of theories below.

Ecologically, tree crown size and shape may be subject to multiple tradeoffs, including lateral extension for light interception and competitor suppression versus mechanical risk, leaf exposure versus drought risk, and fast versus slow growth strategies (Verbeeck et al., 2019). The above general theories do not deny mechanical and other constraints. Instead, adaptive differentiation either does not figure importantly in them or plays a secondary role. Nonetheless, general theory provides a baseline by which to assess their underlying assumptions and hypothesized drivers of variation in canopy dimensions.

Ecological theory appears best able to explain sapling crown form, when variation in crown shape and form is determined more by genetics than stochastic and local processes (e.g. King, 1998;Sterck and Bongers, 1998). Understanding variation in adult crown form, however, has proven more difficult for a number of reasons. These reasons likely include the fact that numerous architectural paths can lead to similar forms (e.g. Fisher and Hibbs, 1982); the measurement of adult crowns is more difficult than that of juvenile crowns; and general and stochastic processes may overpower competitive and genetic ones, making the latter effects difficult to detect.

Mixed support exists for general hypotheses of crown size, including MST (Muller Landau et al., 2006;Enquist et al., 2009;Iida et al., 2011;Pretzsch and Dieler, 2012;Antin et al., 2013;Taubert et al., 2015;Blanchard et al., 2016) and competitive convergence (Iida et al., 2011). Even when scaling theories in general, and MST in particular, are confirmed, variance around the mean fit can exceed an order of magnitude (Poorter et al., 2015). We consider how the inclusion of information on ecosystem context, biogeographic region, and phylogenetic position can help explain this large variance.

Beyond their fundamental importance in understanding plants and forests, tree crown shapes and sizes are of increasing interest to communities developing dynamic global vegetation models (DGVMs). Representations of canopy structure in these models range from 1-D “single leaf” models, to 2-D multilayer models (e.g. perfect plasticity approximation Purves et al., 2007), to efforts to fuse individual-based models with DGVMs to represent the influence of variable crown allometry on ecosystem dynamics (Fischer et al., 2019). Crown allometries play a role in how trees assemble and compete in forests, so as DGVMs become more complex and attempt to represent individual dynamics, models for crown allometry and plasticity of those allometries are needed.

In this study we examine general patterns in tree crown size (width, depth, surface area, and volume), shape (relative depth), and allometry across an original dataset of 1144 individual tree crowns of 281 species spanning one elevation transect, one savanna-forest-precipitation gradient, and one savanna-forest gradient with constant precipitation, across the paleo- and neo-tropics. We evaluate predictions and underlying assumptions of MST theories in light of our data, and then examine the variance around MST fits from an ecological vantage point. To evaluate the relative contributions of general and context-specific processes, we ask if ecological predictors can improve general models. We pose ecological hypotheses across ecosystems, space, biogeography, and evolutionary history, and test them in conjunction with general predictors (Figure 1, Table 1.). To our knowledge, this study represents the first test across ecosystems, and the first test across environmental gradients, of crown size and shape in the tropics.

**Figure 1.**
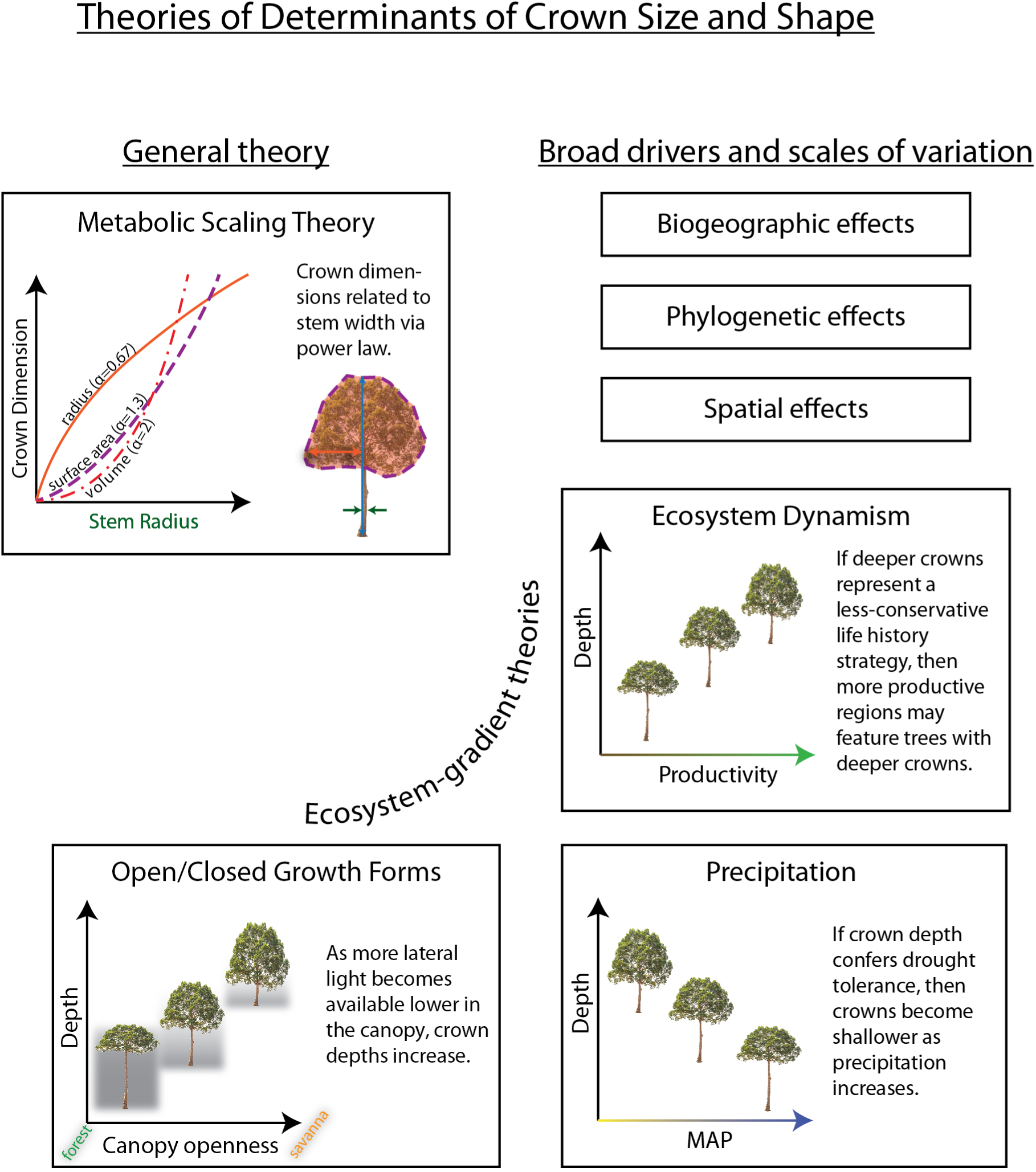
Hypotheses explaining aspects of crown size and shape tested in this study. Results are summarized in Table 1.

**Figure 2.**
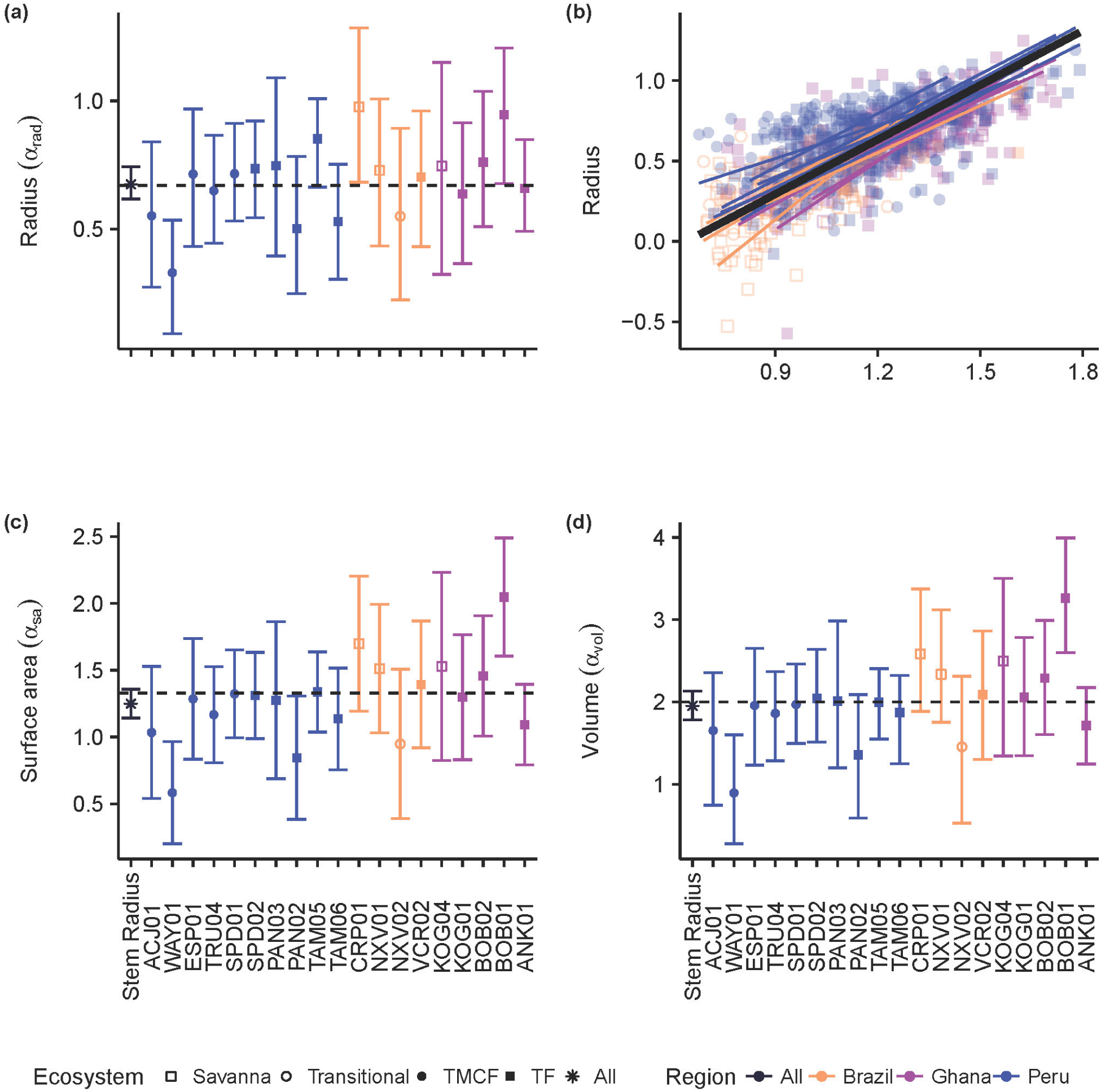
Crown vs stem radius LMM scaling exponents for crown radius (a), surface area (c), and volume (d). Data and SMA fits shown in panel (b) for crown radius scaling (surface area and volume fits are similar, and can be found in SI). “Stem Radius” equates to *α*_*dim*_ in Eq. 3. Black dotted lines indicate *α* predicted by Metabolic Scaling Theory. Site x stem radius interactions are indicated by the site labels below, and per-site intercepts are not shown (see Figure S 9., Figure S 15., and Figure S 20. for β terms and Figure S 14., Figure S 19. for SMA fits vs data). Model includes random intercepts and slopes per species. Error bars indicate 95% confidence intervals fit by likelihood surface profiling. Categorical site variables were encoded as deviation contrasts, which fits the magnitude of the site coefficients as the mean of the slope at that site minus the grand mean slope across all individuals. In order to aid interpretation, we added the grand mean to each site effect and its confidence interval. The Stem Radius effect represents the main effect of stem radius on crown radius, and in this context is the grand mean across all individuals, taking the species random effects into account.

**Table 1.**
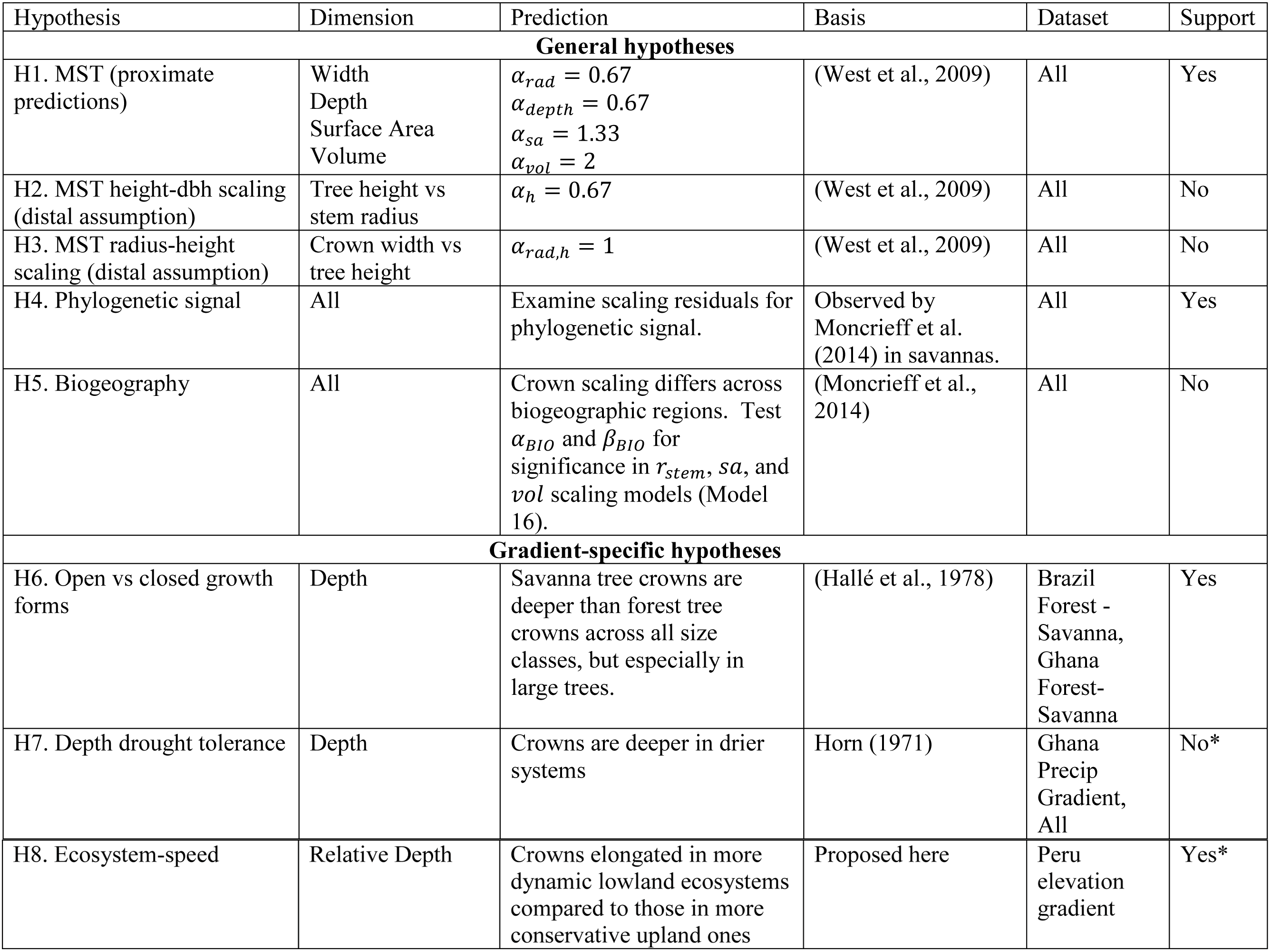
Hypotheses tested in this study. * indicates equivocal support or lack thereof.

| Hypothesis | Dimension | Prediction | Basis | Dataset | Support |
| --- | --- | --- | --- | --- | --- |
| **General hypotheses** |  |  |  |  |  |
| H1. MST (proximate predictions) | Width<br>Depth<br>Surface Area<br>Volume | $\alpha_{rad} = 0.67$<br>$\alpha_{depth} = 0.67$<br>$\alpha_{sa} = 1.33$<br>$\alpha_{vol} = 2$ | (West et al., 2009) | All | Yes |
| H2. MST height-dbh scaling (distal assumption) | Tree height vs stem radius | $\alpha_h = 0.67$ | (West et al., 2009) | All | No |
| H3. MST radius-height scaling (distal assumption) | Crown width vs tree height | $\alpha_{rad,h} = 1$ | (West et al., 2009) | All | No |
| H4. Phylogenetic signal | All | Examine scaling residuals for phylogenetic signal. | Observed by Moncrieff et al. (2014) in savannas. | All | Yes |
| H5. Biogeography | All | Crown scaling differs across biogeographic regions. Test $\alpha_{BIO}$ and $\beta_{BIO}$ for significance in $r_{stem}$, $sa$, and $vol$ scaling models (Model 16). | (Moncrieff et al., 2014) | All | No |
| **Gradient-specific hypotheses** |  |  |  |  |  |
| H6. Open vs closed growth forms | Depth | Savanna tree crowns are deeper than forest tree crowns across all size classes, but especially in large trees. | (Hallé et al., 1978) | Brazil Forest - Savanna, Ghana Forest-Savanna | Yes |
| H7. Depth drought tolerance | Depth | Crowns are deeper in drier systems | Horn (1971) | Ghana Precip Gradient, All | No* |
| H8. Ecosystem-speed | Relative Depth | Crowns elongated in more dynamic lowland ecosystems compared to those in more conservative upland ones | Proposed here | Peru elevation gradient | Yes* |

### 2.1 HYPOTHESES

MST’s predictions of crown allometry start with assuming the principle of elastic similarity that posits the relationship between the height of a tree is proportional to the 2/3 power of stem radius (McMahon and Kronauer, 1976):

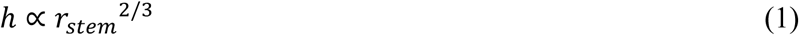

Second, West et al. (2009) assume that crown radius scales isometrically with tree height:

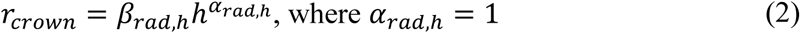

Third, by substitution, the relationship between crown and stem radius is derived as:

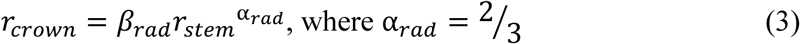

We refer to the first two assumptions (Eqs 1, 2) as distal assumptions, and the derived predication of *r*_*crown*_ (Eq 3) as the proximate prediction. By further assuming a spherical, Euclidean-uniform (as opposed to fractal; Voss, 1988) crown, West et al. (2009) extend the relationship to crown surface area,

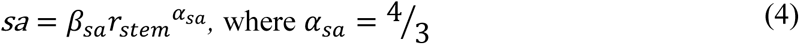

and to crown volume,

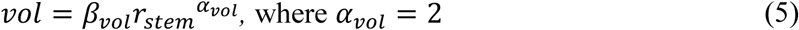

See SI for an illustration (Figure S 1) and more detailed derivation.

MST does not make specific predictions for crown depth, but as we show in the supplemental material, uniform Euclidean assumptions about crown shapes lead to the same prediction for crown depth as for crown radius: *α*_*depth*_ = 0.67. MST would furthermore predict a scaling exponent for relative crown depth (depth/width) as ∝_*reldepth*_=∝_*depth*_−∝_*rad*_ = 0, which posits a lack of relationship between tree size and relative crown depth.

We first test the proximate predictions for crown scaling parameters above against our entire dataset (H1). We also test the crown radius versus height (Eq. 1, H3) and height versus stem radius (Eq. 2, H2) distal assumptions that underlie the scaling predictions (Price et al., 2012). Next, using the residuals of our scaling models, we examine how the variance from the mean fit is partitioned across taxonomic (genus and family), ecosystem type, spatial (plot), and biogeographic (regional) groups. We then examine those groups in more detail. Finishing our treatment of MST, we show how additional modifications of MST can be used to further assess the origin of variation in tree crown scaling.

We then turn to examine variation in crown scaling across environmental gradients and phylogeny to help explain the residual model variance. Specifically, we test whether crown scaling changes as a result of biogeography (H5) and phylogenetic relationship (H4) in our dataset. The variation of crown shapes across biogeographic regions and evolutionary history has not been a focus of extensive research. Blanchard et al. (2016) examined crown allometry across the wet tropics and found little biogeographically-structured variation in scaling exponents. Moncrieff et al. (2014) found that differences between crown shapes in African and Australian savannas were associated with biogeographic region and evolutionary history, not environmental variation. Different evolutionary processes and rates of traits do not lead to distinct phylogenetic patterns (Revell et al., 2008). Nonetheless, as we expect that crown size and shape is variable, adaptive, and that aboveground resources can be partitioned, we do expect to observe a phylogenetic signal. We do not expect to see a strong biogeographic signal given the large range of ecosystems we sample across each biogeographic region.

Across the individual environmental gradients, we test specific explanations of crown shape. “Open” growth form trees are characterized as being shorter and having wider, deeper, and more hemispherical crowns than trees of equivalent girth in the closed forest (Hallé et al., 1978;Harja et al., 2012). Does crown allometry reflect open vs closed growth forms across savanna – forest gradients (H6)? Horn (1971) hypothesized that deep crowns confer drought tolerance due to the self-shading of leaves. Are crowns of similarly sized trees deeper in drier systems as compared to wetter ones (H7)? And finally, Horn’s (1971) framework posits that fast growing trees should have deeper crowns and lower density wood than slower shade-tolerants. Do crowns become relatively shallower, and hence more conservative, in systems with slower nutrient and carbon turnover (H8)?

## 3 METHODS

### 3.1 STUDY SITE

Different environments may select for different crown geometries. For example, differences in resource competition, climate, fire, and herbivory, may confer advantage to crowns of different shapes or sizes. We assessed the scaling of crown shape across 3 environmental gradients: a 3300m elevation gradient extending from the Andes down to the Amazon in Peru; a savanna-forest transition in Brazil; and a savanna-forest precipitation transition in Ghana. In addition to continuous environmental variables, our study distinguishes between categorical ecosystem types: tropical forests (TF), tropical montane cloud forests (TMCF), savannas, and transitional sites between savannas and forests.

Our study employs 20 one-hectare plots across the three environmental gradient transects above (Table S 1): ten plots from the Peruvian elevation gradient, four plots from the Brazilian savanna-forest transition, and six plots from the Ghanian savanna-forest precipitation transition. These plots comprise part of the Global Ecosystems Monitoring Network (GEM; http://gem.tropicalforests.ox.ac.uk).

The ten Peruvian plots (Figure S 2) belong to a group of permanent 1-ha plots in the departments of Cusco and Madre de Dios in SE Peru. Six of the plots are montane plots in the Kosñipata Valley in the province of Paucartambo, department of Cusco (Malhi et al., 2016), spanning an elevation range 1500 - 3500 m (Malhi et al., 2010), two are submontane plots located in the Pantiacolla front range of the Andes (range 600 - 900 m), and two plots are located in the Western Amazon lowlands in Tambopata National Park (range 200 - 225 m). All the Peruvian plots are operated by the Andes Biodiversity Ecosystems Research Group (ABERG, http://www.andesconservation.org). From Feb 2013 to Jan 2014, mean annual air temperature (MAT) varied from 9°C to 24.4°C and mean annual precipitation (MAP) ranged from 1560 mm y^-1^ to 5302 mm y^-1^ across all sites along the gradient (Table S 1). Precipitation peaks strongly at ∼1500m elevation. Productivity and ecosystem carbon cycle data for these sites are reported by Malhi et al. (2017).

The Brazilian forest–savanna gradient was located near Nova Xavantina, Mato Grosso. Two well-defined seasons occur in the region: hot and wet from October to March, and cool and dry from April to September (Marimon et al., 2014). The gradient spans Amazon forest and Cerrado biomes, transitions rapidly from an Amazonian transitional dry forest to *cerrado sensu stricto* (South American woody savanna); abrupt transitions in vegetation type accompany changes in soil physical and chemical properties (Marimon Junior and Haridasan, 2005). Three of the four plots are located in Parque Municipal do Bacaba, and represent three distinct vegetation types of progressively decreasing woody biomass and stature (cerradão, cerrado tipico, and cerrado rupestre, see Marimon Junior and Haridasan, 2005). Despite fire exclusion by park management, the first two Bacaba sites burned in 2008, while the last has not burned in over twenty years. The fourth site is located in a semi-deciduous forest, located 25 km away (SE) from Nova Xavantina in Fazenda Vera Cruz, and is not subject to a fire regime (Marimon Junior and Haridasan, 2005;Marimon et al., 2014). The close proximity of all plots suggests that variation in vegetation type is driven by edaphic properties rather than by variation in rainfall.

The Ghanaian forest-savanna precipitation gradient spans three sites. Three plots are found in the Kogyae Strict Nature Reserve, in the northeast of the Ashanti region, with MAP of 1360mm and MAT of 28°C (Janssen et al, in revision). Two semi-deciduous forest plots are in the Bobiri Reserve, 25 km from Kumasi, with a MAP of 1500mm. The two final plots, moist evergreen and swampy evergreen forests, are in the Ankasa Reserve in southwestern Ghana with a MAP of 2000mm (Chiti et al., 2010). The soils in the Ghana transect are more fertile than those in the Brazil sites (Malhi unpublished data).

### 3.2 FIELD SAMPLING

From April – November 2013, we measured plant traits as part of the CHAMBASA project in Peru, from March – May 2014 as the BACABA project in Brazil, and from October 2014 – March 2015 as the KWAEEMMA project in Ghana. Based on the most recently available census and diameter data, a sampling protocol was adopted wherein species were sampled that maximally contributed to plot basal area (a proxy for plot biomass or crown area). We aimed to sample the minimum number of species that contributed to 80% of basal area, although in the diverse lowland forest plots we only sampled species comprising 50-70% of plot basal area. Within each species, 3-5 individual trees were chosen for sampling (5 trees in upland Peruvian sites and 3 trees in lowland Peruvian, Brazilian, and Ghanaian sites; Table S 2). If 3 trees were not available in the chosen plot, we sampled additional individuals of the same species from an area immediately surrounding the plot.

Tree climbers sampled a fully sunlit canopy branch at least 1 cm diameter of each tree, from which simple leaves, or individual leaflets from compound-leaved species (both referred to as ‘leaf’ below), were removed and measured. In the case of compound leaves, the entire compound leaf was also collected for whole-leaf area calculations. Branches and leaves were chosen with minimal damage (e.g. from herbivory). Five leaves per branch were sampled in Peru, and three per branch were sampled in Brazil and Ghana.

### 3.3 CROWN MEASUREMENT

Tree crowns were measured using a laser hypsometer (TruPulse 360/360R, Laser Technology Inc., Colorado, USA). In “Missing Line” (ML) mode, the TruPulse 360 returns horizontal distance (HD), vertical distance (VD), and azimuth (AZ) between two points in space as determined by 2 laser pulse returns that determine distance coupled with azimuth measurements from the unit’s internal compass (TruPulse 360/360B User’s Manual, Laser Technology Inc., Colorado, USA). We took ML measurements between stem base and crown top, crown bottom, and usually 6-20 points around the circumference of the crown perimeter depending on the complexity of its shape. Because we were interested in tree function (e.g. gas exchange capacity), we defined the crown base as the lowest significant foliage that was not a resprout or otherwise relatively spurious, instead of using the first primary branch. We applied a convex hull to this set of points to yield a 3-D polyhedron from which crown dimensions were extracted. We estimate crown volume and surface area as the volume and surface area of the convex hull. We estimate average crown radius as the radius of a circle with the same area as the 2-D convex hull of the points when projected onto a horizontal plane.

Manual measurements of crown widths and tree heights were taken using clinometer and tape measure in a subset of Ghanaian trees to verify that results from the laser hypsometer method are comparable to those from more widely used techniques. Dimensions from both methods corresponded closely (N = 20; width R^2^_adj_ = 0.83, SE = 1.06; height R^2^_adj_ = 0.74, SE = 2.77), and are within the range for the tangent method reported by Larjavaara and Muller-Landau (2013). Larjavaara and Muller-Landau (2013) recommend the laser hypsometer method as opposed to manual clinometers for tree height measurement, especially for cross-site studies where instrument operators are different. They further note that while errors are inherent in both methods, those introduced by the laser hypsometer may lend greater weight to leaf versus fine wood structure, which largely corresponds with our goals here.

### 3.4 INTRASPECIFIC VS INTERSPECIFIC MODELS

MST’s predictions of allometric scaling assume that the terminal vascular characteristics are approximately the same within and across species (West et al., 1997;West et al., 1999). These characteristics include: petiole conducting area, petiole length, and flow rate (West et al., 1997;Savage et al., 2008). In addition, to scale up to crown characteristics, West et al. (2009) assume that leaf area densities (LADs) of tree crowns are equivalent across species and tree size. If these assumptions do not hold across species then we must control for this variation in one of two ways: either measure the key characteristics in each individual or species, or evaluate scaling parameters within (intraspecific), not across (interspecific) species. In effect, without independent measurements of these characteristics, MST’s assumptions compel us to model scaling intraspecifically. In this study, we implement intraspecific LMMs by including species as a random effect. Random effects enable us to account for variation between species with sample sizes of 3-5 individuals per species. In our standardized major axis (SMA) regressions (see Appendices), we group by species, and then examine the estimated overall fit.

### 3.5 NOTATION

Our notation of model coefficients and parameters uses α when referring to exponential (or scaling) parameters, and *β* when referring to linear (or normalization) parameters. Subscripts may include a comma, such as α_*y,x*_, in which case the first term (*y*) indicates the dependent variable in the model, and the second term (*x*) indicates the independent variable that coefficient is associated with. Thus, α_*depth,h*_ indicates the exponential parameter of tree height when predicting crown depth. Since *r*_*stem*_ is the most common predictor, we omit commas for parameters associated with *r*_*stem*_.

### 3.6 STATISTICAL AND PHYLOGENETIC MODELS

We employed both LMM and SMA regressions to estimate scaling parameters and test hypotheses. Our primary analyses are conducted with LMM, and confirmed with SMA in the Appendices. The LMM approach is advantageous when accounting for variation across many groups (e.g. species) via random effects, and it is appropriate for fitting allometry models (Kilmer and Rodríguez, 2016). SMA regression is often used to estimate scaling coefficients as it allows for measurement error in both dependent and independent variables (Warton et al., 2006), and while less flexible, we include it as convention dictates.

LMM grand means were estimated by deviation (or sum-to-zero sensu Crawley, 2012) coding of categorical grouping factors (usually grouped by site), which effectively fits the main effect as the grand mean across all individuals while taking random effects into account. In order to aid interpretation of figures, we add the grand mean to each site effect and its confidence interval unless indicated otherwise. All analyses were performed in the R programming environment (R Core Team, 2016), accessing data using the GEMTraits database and R package (Shenkin et al., 2017), using the lme4 R package (ver 1.1-13; Bates et al., 2015) to fit the LMM models, and the afex R package (ver 0.16-1; Singmann et al., 2016) to estimate variable importance. 95% parameter confidence intervals were computed using profiling methods unless indicated otherwise.

To test the MST predictions above, we fit log-log linear models to estimate the linear (normalization) and exponential (scaling) parameters that relate the crown dimension to the size of the tree. Specifically, if *Y* is the crown dimension and *X* the tree size scalar (e.g. stem radius) in the equation *Y* = *βX*^*α*^, the slopes predicted by these models correspond to the scaling exponents α_*rad*_, α_*sa*_, α_*vol*_, and α_*depth*_, and the intercepts to the normalization factors *β*_*rad*_, *β*_*sa*_, *β*_*vol*_, and *β*_*depth*_ (Eqs. 1-5; Table 2, Models 5 - 12; see Appendices for formula and model derivations).

**Table 2.**
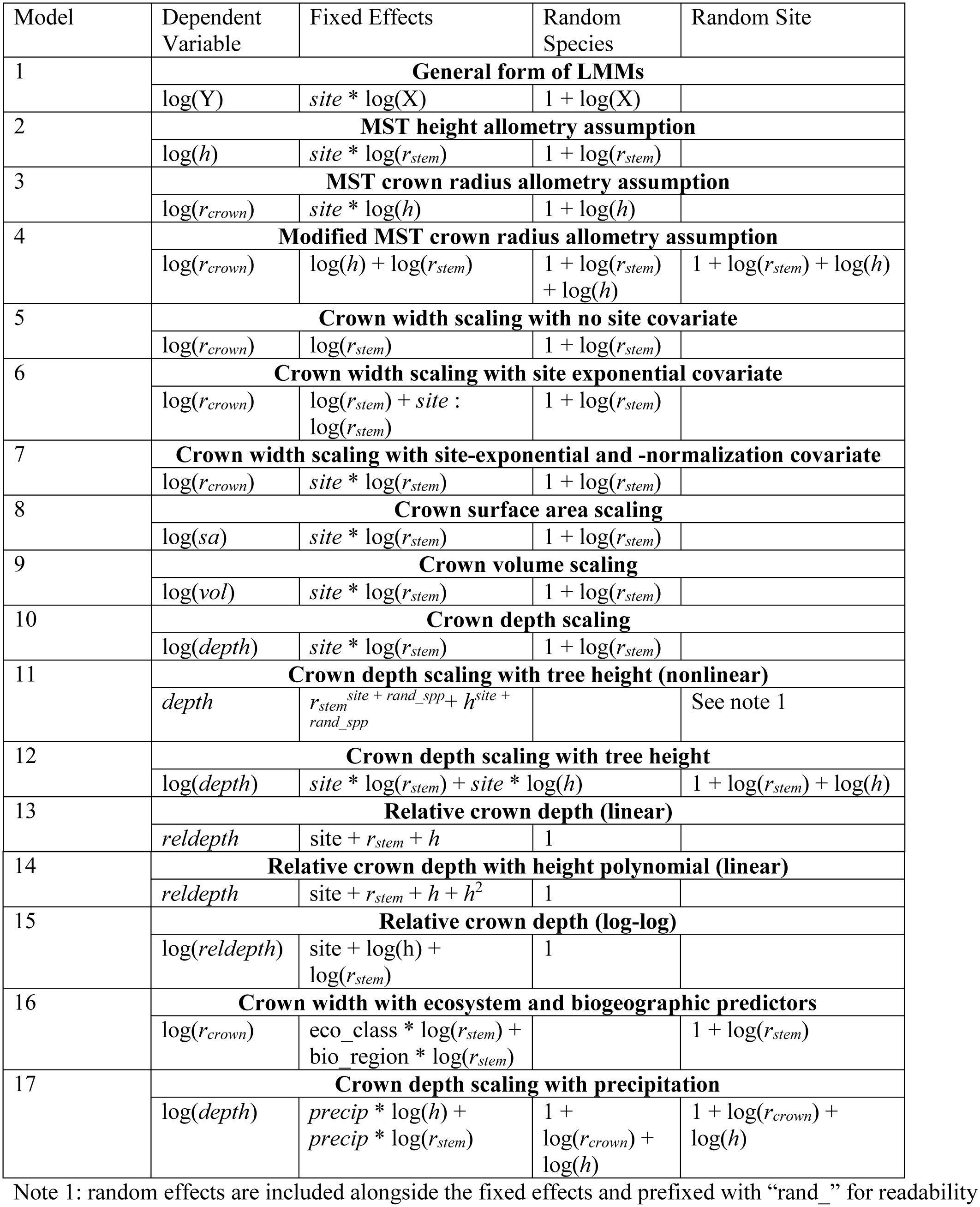
List of modelling equations employed in this study.

LMM log-log models suffice for scaling tests with single predictors, but when multiple independent non-linear predictors are necessary, one must either accept a multiplicative relationship between predictors, or resort to non-linear mixed models (NLMM). We use NLMMs below when testing multiple non-linear predictors that cannot be expressed as a modification to an existing coefficient (see SI). When using NLMMs below, we utilize the nlme R package (Pinheiro et al., 2016), and we include species-level random effects for each predictor unless indicated otherwise. When NLMMs would not converge, we resorted to LMMs.

MST predicts a lack of relationship between tree size and relative crown depth. We therefore fit both linear and log-log models when evaluating relative depth (Table 2, Models 13 - 15; Table S 6). For readability, not all models tested in Table S 6 are listed in Table 2.

To examine whether ecosystem type or biogeography influenced crown size and shape allometries, we fit LMM scaling models for crown width, depth, surface area, and volume with biogeography (Peru, Brazil, and Ghana) and ecosystem type (TF, TMCF, Savanna, and Transitional) as covariates. We illustrate the equation for crown width (Table 2, Model 16), but omit the full suite of models tested (Table S 8) for readability. While MST LMMs included species as a grouping factor, LMMs used for predictions across ecosystems and biogeography (Table 2, Model 16) included site but not species as a random effect, since we were interested in community-level responses. Additionally, inclusion of species-level random effects in the ecosystem and biogeographic models resulted in an excessive number of predictors.

To test the crown depth – drought tolerance hypothesis (H7), we modeled precipitation as a scaled covariate in the crown depth scaling model (Table 2 Model 17). We test the effect of precipitation both across the Ghana precipitation gradient and across the entire study.

Partitioning of variance was computed by fitting crossed and nested random effects for the categorical variables of interest in linear mixed models using the lme4 (Bates et al., 2011) package, extracting the variance-covariance matrix, and computing the variance (*Provarp*) associated with each level (*l*_*i*_) as 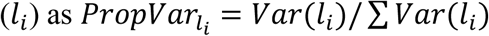. Residuals used in this analysis were produced by fitting models similar to those used for MST tests, but with site-level random intercepts and slopes instead of fixed effects. Residuals were then calculated based on model predictions with random effects set to zero.

We created our phylogeny using Phylocom (Webb et al., 2008) with an APGIII mega-tree (The Angiosperm Phylogeny, 2009; megatree R20100701; available at https://github.com/camwebb/tree-of-trees/blob/master/megatrees/R20100701.new) and ages file (Gastauer and Meira-Neto, 2013). Node values and confidence intervals were computed with maximum likelihood methods using fastANC in the phytools R package (Revell, 2012).

## 4 RESULTS

### 4.1 ALLOMETRIC SCALING OF TREE CROWNS

Here we fit MST models to our data and compare our empirical scaling exponents with theoretical predictions (H1 - H3). Model equations are specified in Table 2. SMA results largely confirm LMM results, and details are included in the appendices.

#### 4.1.1 ASSESSING METABOLIC SCALING THEORY’S DISTAL ASSUMPTIONS

The two assumptions underlying MST crown scaling, namely 2/3 scaling between stem radius and height and isometric scaling between height and crown radius, are both violated in our dataset. Instead, the constant stress model (*α*_*h*_ = 1/2) better explains the relationship between stem radius and height (Model 2; *α*_*h*_ = 0.46, 95% CI 0.42 – 0.51;Figure S 3), and crown radius scales with height as α_*rad,h*_ = 0.63 (CI 0.53 – 0.73, Model 3, Figure S 5), with variation approaching an order of magnitude (Figure S 4, Figure S 5). Thus, H2 (MST height – stem radius scaling assumption) and H3 (MST crown radius – height assumption) are not supported.

The assumption of height-crown isometric scaling (Eq. 2), in particular, lacks empirical underpinnings. Instead, we propose an alternative assumption to better account for the drivers of tree form and function. Specifically, we allow crown radius to depend on both *r*_*stem*_ and *h* independently by modifying the second assumption (Eq. 2) to include *r*_*stem*_ (Model 4; see Appendices for derivation). Thus, the modified assumption becomes:

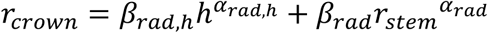

With plot as a random effect, this modification resulted in α_*rad,h*_ = 0.16 (SE = 0.05) and 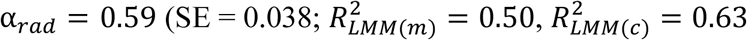; see Appendices for details). Parameter estimates were similar across the various formulations tested, including those without plot as a random effect.

##### 4.1.1.1 Assessing Metabolic Scaling Theory’s Proximate Predictions

We tested three models for predicting crown scaling (Models 5 - 7), and adopted the more complex formulation (Model 7) from here onwards because it fits nearly as well as the best model according to AIC, and is a more conservative choice for hypothesis testing (Bolker et al., 2009) (for details, see Scaling model formulation in Appendices).

Despite the failure of the assumptions underlying MST crown scaling (see above), we found that when aggregated by species (i.e. intraspecific scaling), our models closely agree with MST predictions for crown width, surface area, and volume scaling (Table 3), and thus support hypothesis H1 (MST predictions).

**Table 3.**
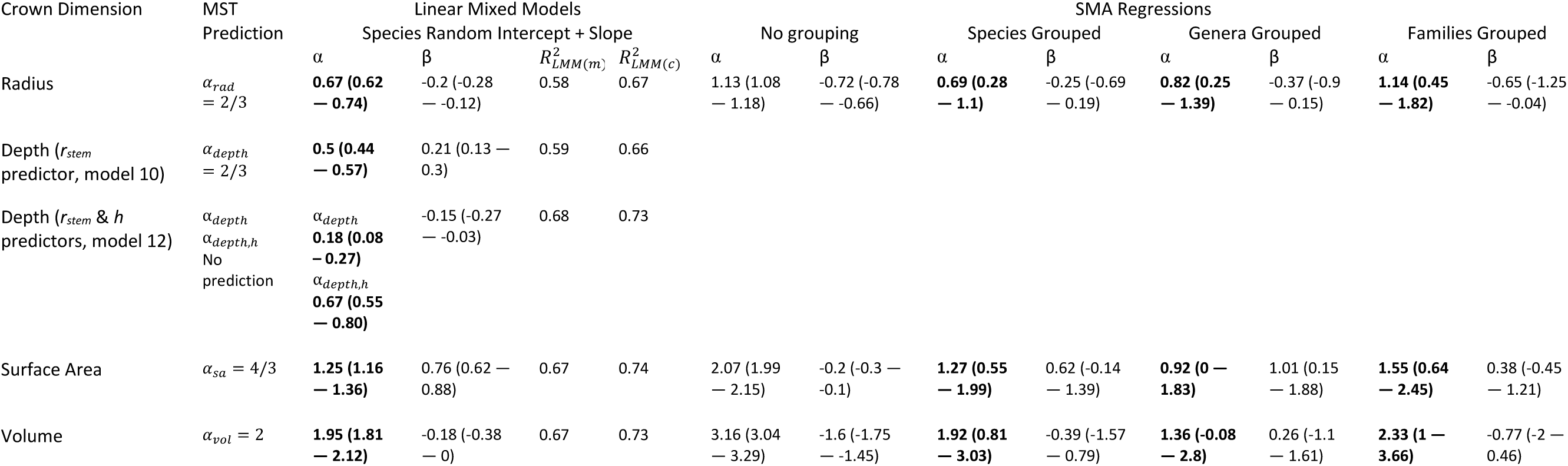
Metabolic scaling parameters as fit by LMMs and SMA regressions.

##### 4.1.1.2 Crown depth scaling

Crown depth was better predicted by tree height than by stem radius, but both predictors were important (AICc of model with just stem radius [10] minus that with both predictors [12] = 18; see Models for crown depth in Appendices for details). We therefore use the model with both predictors, but report the results from themodel with just stem radius for copmrison with the scaling models for other crown dimensions.

Crown depth scales with stem radius as α_*depth*_ = 0.50 (Model 10, Figure S 21, Table 3) and as α_*depth*_ = 0.18, α_*depth,h*_ = 0.67 when also including tree height as a predictor (Model 12, Table 3). Remarkably, in this formulation, crown depth scales exactly with tree height as crown width scales with stem radius. We do not support MST’s prediction that crown depth scales with stem radius as 2/3. Rather, our data suggest that crown depth scales with tree height as 2/3.

### 4.2 EXPLAINING RESIDUAL VARIATION IN SCALING

Taking the residual errors of the MST fits (*ε*_i,j_), we examine how this variation is structured across taxonomy, space (plots), biogeography (region), and ecosystem type (Figure 3, Table S 7). We find that residual variation in crown width leaves ∼50% unexplained, while biogeography accounts for 28%, taxonomic family for 12%, and spatial (between plot) variation for 11%. Crown volume and surface area residuals are similarly structured: unexplained variation accounts for about half of the total, spatial variation account for ∼20%, ecosystems for ∼15%, biogeography for ∼10%, and taxonomic family for 7%. Crown depth residuals (Model 12) were structured across space (24%) and biogeographic region (17%), with no taxonomic or ecosystem signal..

**Figure 3.**
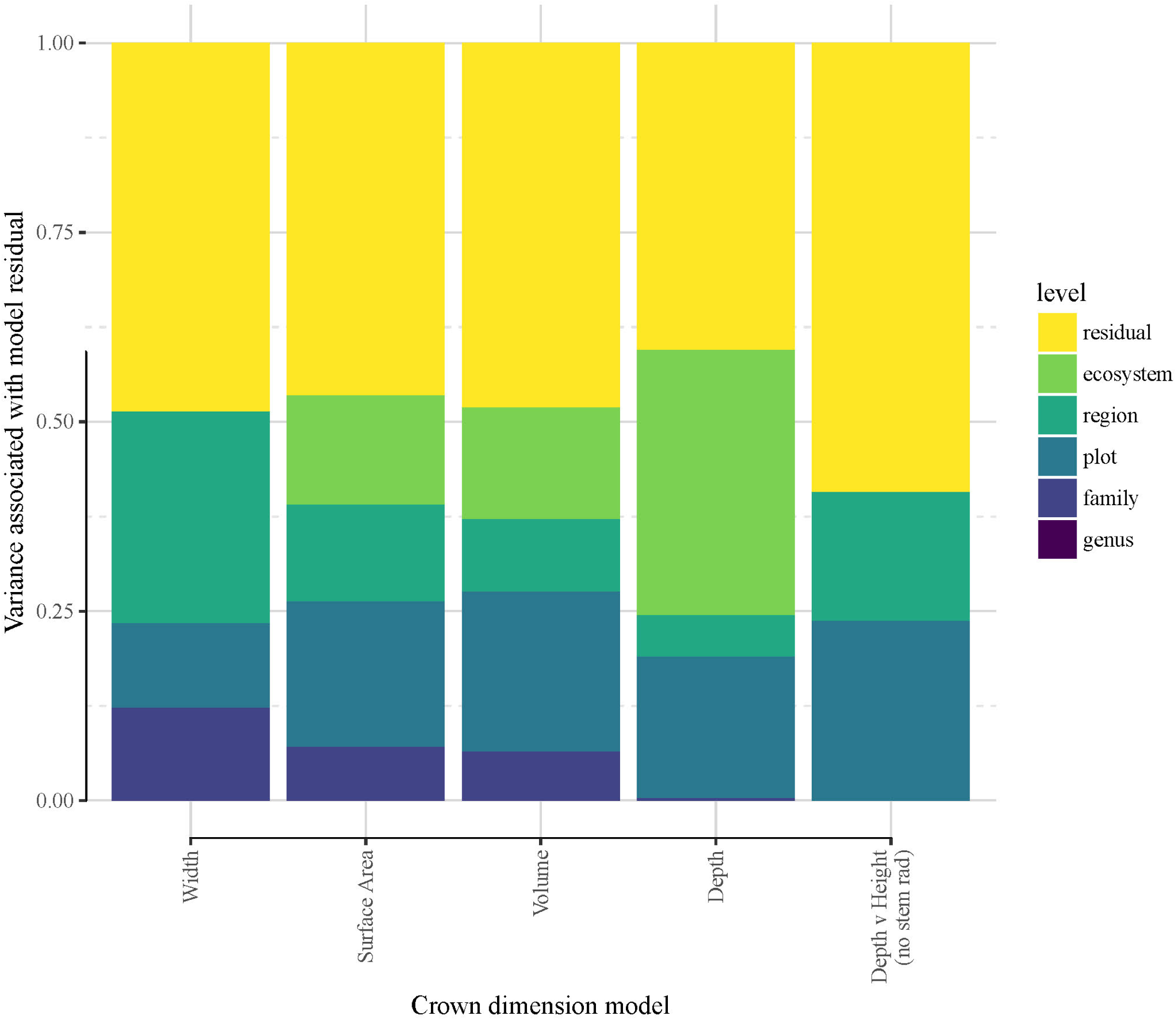
Variance associated with species’ deviation from mean crown scaling relationship The quantity analyzed here is the mean species residual, or deviation, from the crown_*d*imension = *βr*_*stem*_^*α*^ scaling model. LMM models were re-fit with site as a random effect, and residuals were obtained with the random effects removed.

#### 4.2.1 PHYLOGENETIC STRUCTURE OF CROWN SCALING

Crown scaling of most families do not differ significantly from each other. The crowns of Fabaceae, and particularly those of the Mimosoideae and Papilionoideae subfamilies, however, are consistently larger for a given stem radius than those of other taxa across all three study regions (Model 7, Figure 4, Figure S 22, Figure S 23). This clear phylogenetic signal supports our hypothesis that such a signal exists (H4 (Phylogenetic signal)). Crown surface area and volume allometries show similar phylogenetic patterns, with phylogenetic differences in volume being the strongest, surface area less so, width weaker, and depth lacking a Fabaceae signal (Model 10, Figure S 25). Other clades with large crowns include *Celtis* and *Tapira*. Crowns of trees in the Chrysobalanaceae and Primulaceae, and some Moraceae taxa, tend to be smaller than expected. These patterns do not seem to be structured by biogeography. That is to say, clades with significantly larger or smaller crowns are largely comprised of species from all three regions.

**Figure 4.**
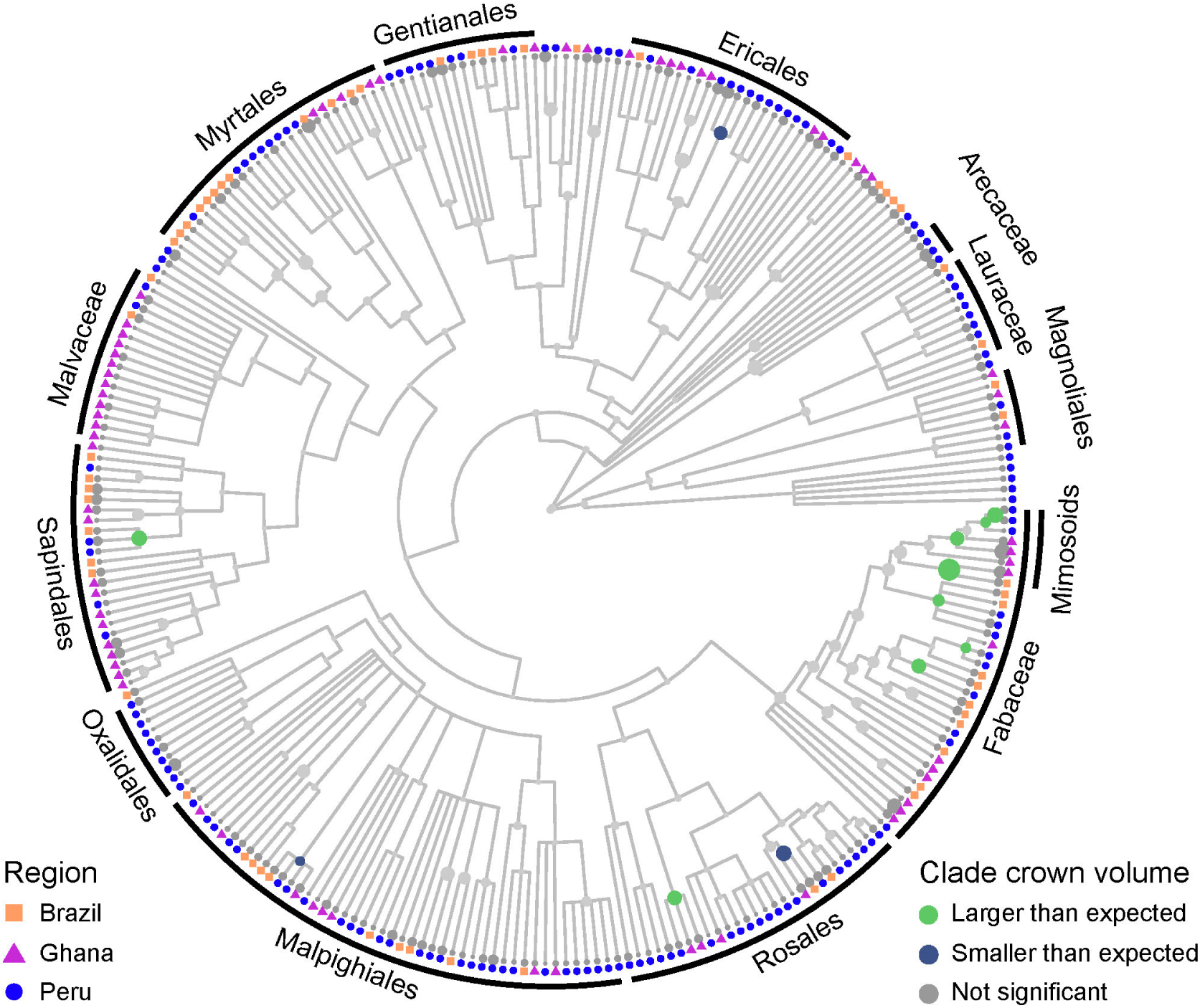
Mean per-species residuals from the crown volume vs stem radius scaling LMM with species as a random effect mapped onto a tree species phylogeny (see Methods for details of phylogeny construction). A residual in this instance implies a difference in intercept, not slope, in the model. Size of circle corresponds to size of residual. Internal node states were determined using fastANC (see Methods), with colors indicating direction and confidence of internal node estimates: 95% confidence interval of model residuals of grey nodes intersects zero, green nodes indicate clades with larger than expected crowns, and blue nodes clades with smaller than expected crowns. Tips are not evaluated for significance and are therefore not colored. Similar plots with species names are included in SI (Figures S 22 - S 25).

**Figure 5.**
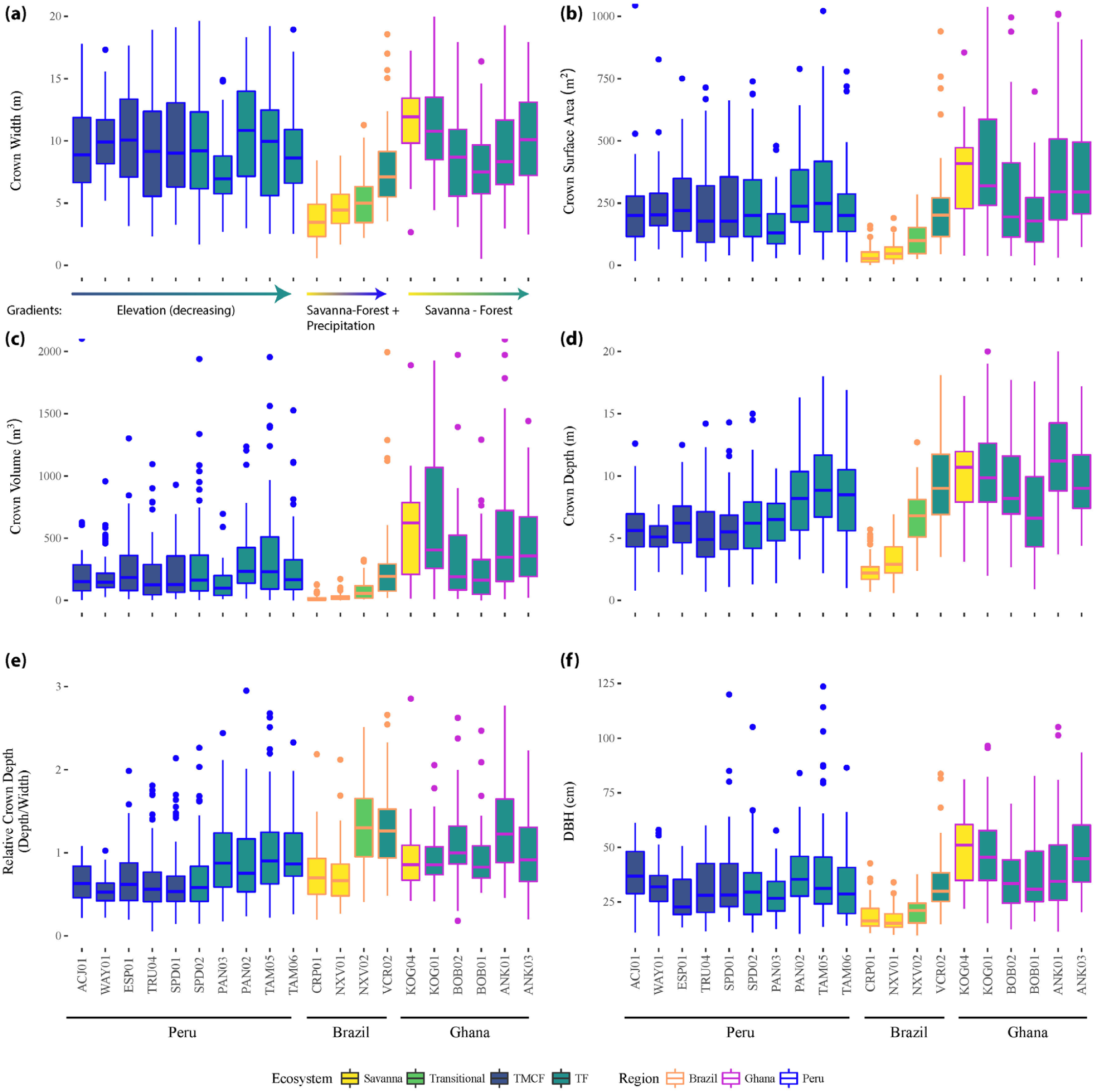
Observed crown volume (a), radius (b), depth (c), and relative depth (d) across sites (x axis), ecosystems (fill color), and regions (outline color and annotation). See tree height in SI (Figure S 30.). Crowns with large relative depth values are elongated, and those with low relative depth are flatter. The data here are descriptive and have not been modeled or corrected for tree size or any other variable.

### 4.3 TRENDS ACROSS ECOSYSTEMS AND BIOGEOGRAPHIC REGIONS

Here we examine patterns and test hypotheses of crown scaling patterns across environmental gradients (H6 - H8) and biogeography (H5). We discuss our observations of these trends in detail in the Appendices (see Observed Trends across ecosystems and biogeographic regions).

Both ecosystem normalization (*β*_*eco*_) and exponential (*α*_*eco*_) covariates significantly improve surface area (Kenward-Roger approximation; P(*α*_*sa,eco*_) = 0.02, P(*β*_*sa,eco*_) = 0.003) and volume models (P(*α*_*vol,eco*_) = 0.01, P(*β*_*vol.eco*_) = 0.001). Ecosystem type is only significant in crown width (Kenward-Roger approximation; P(*α*_*rad,eco*_) = 0.02, P(*β*_*rad,eco*_) = 0.005) and depth (Kenward-Roger approximation; P(*α*_*depth,eco*_) < 0.0001, P(*β*_*depth,eco*_) = 0.004) in models with biogeography removed. Tree height but not stem radius predictors is significant in these crown depth models. Predicted crown dimensions across ecosystems are presented in Figure 7.

**Figure 6.**
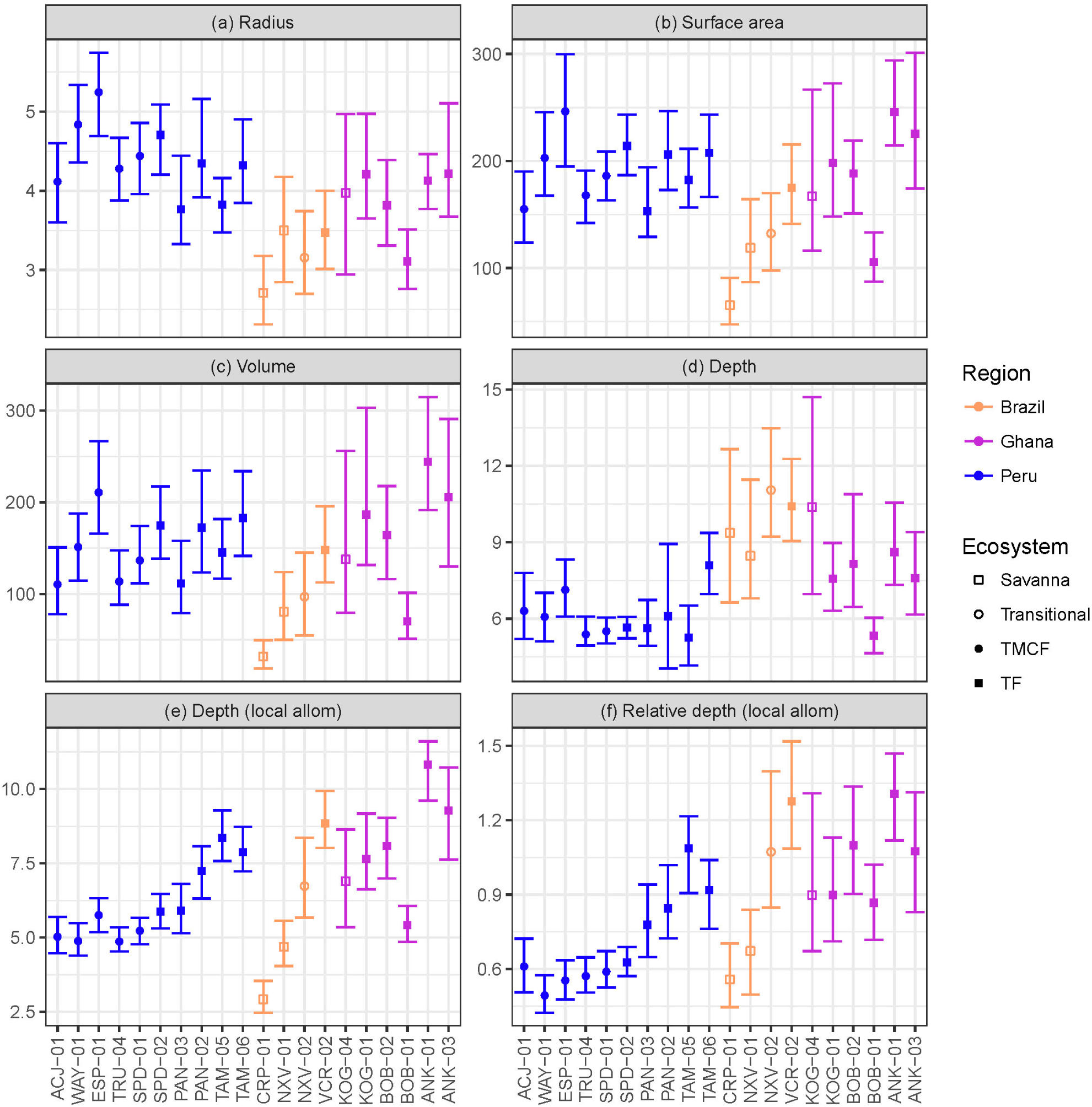
Modeled crown dimensions and 95% CIs for a 30cm DBH tree. The crown depth model also included tree height, which was fixed in (d) using our height vs stem radius allometry (see SI) and averaged across sites so the same DBH and height were used for each site, and allow to vary in (e) and (f) based on the mean per-site height – DBH allometry (Model 2). Predictions are derived from the following LMM models described above: (a) Model 7, (b) Model 8, (c) Model 9, (d) Model 12, (e) Model 12, (f) Model 15. CIs estimated using a bootstrap algorithm with 100 iterations. Units are m, m^2^, and m^3^. See Figure S 31 for predictions for other tree sizes.

**Figure 7.**
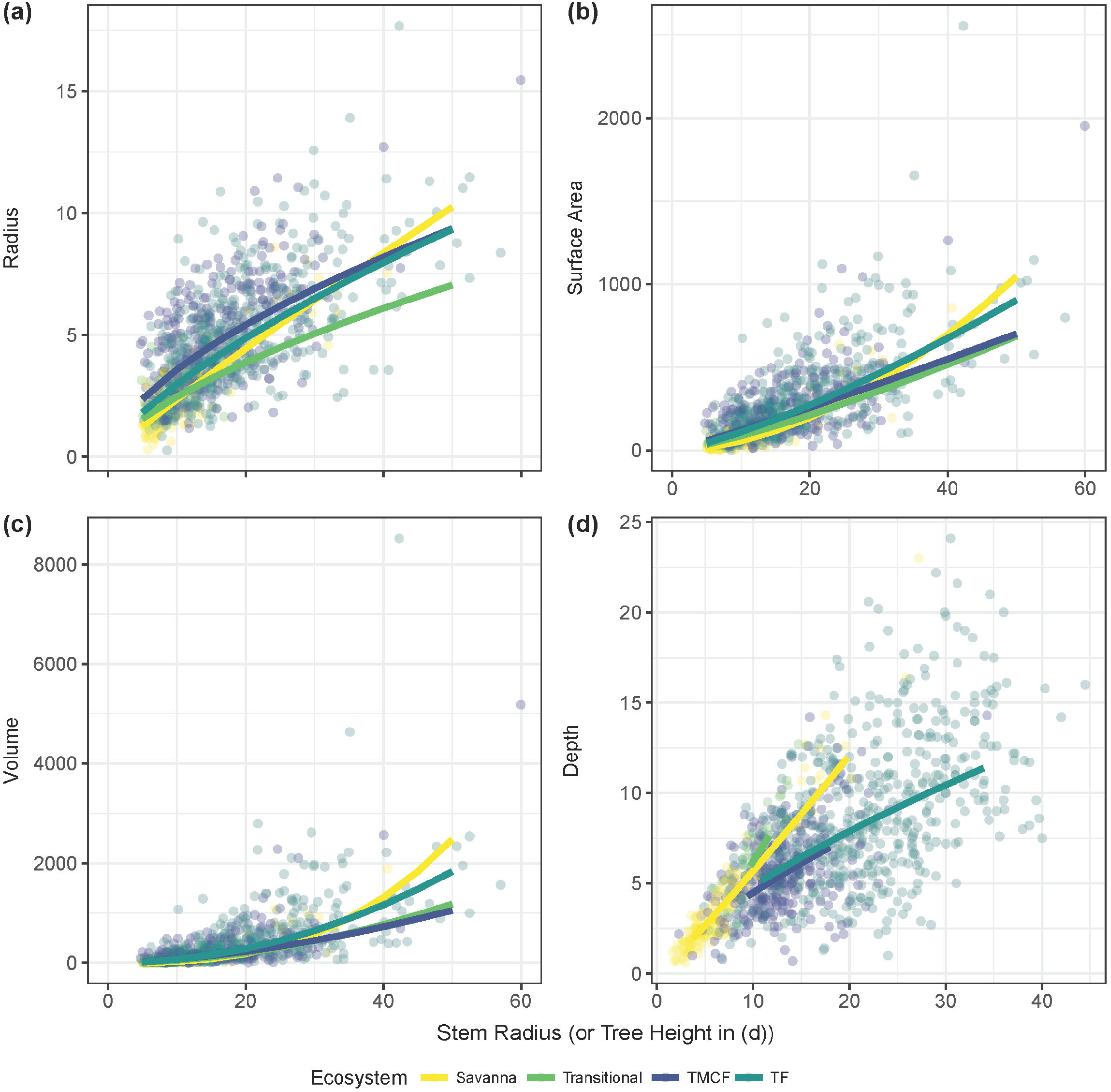
LMM predictions (lines) and individual trees (points) across all regions, grouped by ecosystem type. The crown depth model includes both stem radius and tree height, and is plotted against tree height. For predictions, ecosystem-specific DBH-height allometries were constructed and used.

Our linear and variance partitioning models disagree about the role of biogeography in crown scaling. Linear models were generally unimproved by the addition of the biogeography predictor (Table S 8). The variance partitioning models, however, attribute substantial variance to biogeography, and especially in the case of crown width scaling (Figure 3). The unusual formulation necessitated by log-log models with covariates (see Appendices) may explain this disagreement. We conclude that while biogeography is associated with some variance, it is a weaker influence than ecosystem type.

#### 4.3.1 ENVIRONMENTAL GRADIENT HYPOTHESES

The observed trends across environmental gradients are described in detail in the Appendices (see Observed trends across gradients within regions). Here we evaluate our specific hypotheses related to the environmental gradients our study spanned.

##### 4.3.1.1 Savanna-Forest Transitions

Savanna trees are shorter than forest trees for the same girth (Figure S 33). Consequently, because crown depth scales principally with tree height, a 30cm DBH savanna tree is shorter and has a shallower crown than a 30cm DBH forest tree (Figure S 32). When controlling for tree height however, a 15m-tall savanna tree crown is more than 50% deeper than a 15m-tall forest tree (Figure 7). Stout savanna trees with deep crowns are consistent with our hypothesis H6 (Open growth form).

##### 4.3.1.2 Precipitation Gradient

We tested for effects of precipiation on crown depth across the Ghanaian transect, and across our entire study. No precipitation covariates were significant across the Ghanaian transect (Table S 9, Table S 10). Modeling the effect of precipiation on crown depth across all sites, just one precitation covariate was marginally significant (*P*α_*precip,h*_ = 0.07; Table S 9, Table S 10). Taken together, our analyses do not support hypothesis H7 (Depth drought tolerance).

##### 4.3.1.3 Elevation Gradient

We hypothesized that crowns will become deeper as one moves from the low productivity sites in the Andes down to the high productivity sites in the Amazon (H8 (Ecosystem speed)). Crown widths decreased going downslope for small (10cm) and mid-sized (30cm) trees (Figure 6, Figure S 33a), and predictions from the linear model (Model 13) indicated that relative crown depth increased downwards through the transect. The elongation of crowns moving downslope is apparent when taking local height allometries into account (Figure 6e,f), but less so when assuming constant height allometries across sites (Figure 6d, Figure S 28). Our ecosystem speed hypothesis H8 is therefore supported.

## 5 DISCUSSION

Our findings (Table 1) support the view that general MST processes describe a central tendency for the scaling of tree crowns (Muller Landau et al., 2006;Enquist et al., 2009;Pretzsch and Dieler, 2012;Antin et al., 2013;Taubert et al., 2015;Blanchard et al., 2016;Farrior et al., 2016), and that this tendency is influenced by ecosystem and evolutionary context. Scaling did not change significantly across regions, but it did across ecosystem types with savanna crowns growing more quickly with tree size than those of forests. We found that crown radius scales with stem radius (King, 1996) whereas crown depth scales with tree height, and because tree height differs more across ecosystems than stem girth, crown depth varies more across gradients that crown depth. Controlling for tree size, crowns of legumes were wider and larger than those of other taxa. Implications of specific results are discussed below.

### 5.1 METABOLIC SCALING THEORY

MST has been challenged on a number of grounds for not corroborating some empirical data sets and for not providing the most parsimonious explanation (e.g. Muller Landau et al., 2006). Perhaps the strongest criticism from forest ecologists stems from the absence of competitive and environmental considerations in the theory (Coomes and Allen, 2009;Coomes et al., 2011;Stark et al., 2011;Pretzsch and Dieler, 2012), though some attempts have been made to incorporate these influences (Price et al., 2010;Price et al., 2012;Stark et al., 2015). Our results here, tested across contrasted of tropical ecosystems, environmental gradients, and biogeographical contexts, serve as robust yet qualified confirmation of MST predictions for the relationship between crown dimensions and stem radius.

While previous studies have disagreed on whether MST provides an accurate explanation for crown scaling, those that implement interspecific models show poor fits between theory and data; those that use intraspecific models show generally good fits (e.g. Muller Landau et al., 2006;Pretzsch and Dieler, 2012;Blanchard et al., 2016) (but see Tredennick et al., 2013). Because MST assumes that certain key characteristics are constant across individuals (assumption 7 in Savage et al., 2008), intraspecific models are the appropriate framework for tests of MST in the absence of measurements of those key characteristics. Indeed, our own intraspecific models strongly differed from our interspecific ones. Overall then, this study adds its support of MST crown scaling across environmental gradients and biogeographic regions to the general support MST crown scaling finds in other studies.

#### 5.1.1 MODIFICATION OF MST CROWN SCALING ASSUMPTIONS

Despite MST’s almost exact prediction of our empirical *α*_*rad*_= 0.67, neither of West *et al.’s* (2009) component assumptions of stem radius vs tree height and tree height vs crown radius relationships hold. Both empirical relationships were found to have significantly smaller exponents than those predicted by MST. While West et al. (1999) acknowledge that height allometries vary (Niklas, 1995;Nogueira et al., 2008;Feldpausch et al., 2011;Banin et al., 2012), the principal MST crown scaling prediction relies on a value of *α*_*h*_= 0.67.

MST predictions rely on chains of relationships. In this case, the linkage is between stem radius, stem height, and ultimately crown width. We modified MST’s stem height/crown width assumption to include stem radius (Model 4), and found that the modification resolved the previously-incongruent chain of scaling exponents (see Appendices, Resolving the incongruence of MST distal assumption deviation and MST proximate prediction accuracy). We propose that the following assumptions be used instead of equations 1 and 2:

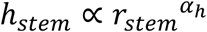

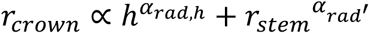

where *α*_*h*_ = 1/2. Empirically, we find that *α*_*rad,h*_ = 0.16 and *α*;_*rad*_′ = 0.59, but these values should be informed by further development of theory before offering them as MST assumptions.

### 5.2 PHYLOGENETIC VARIATION

While crown scaling did not vary amongst most families, we do find a strong, consistent, and biogeographically-wide phylogenetic signal in tree crown allometry in the large Fabaceae crowns, and in particular, in the Mimosoideae and Papilionoideae subfamilies. What is it about these subfamilies that could lead to their particularly wide crowns?

Of the three legume subfamilies, rhizobial associations that enable nitrogen fixation in root nodules are common in the Mimosoideae and Papilionoideae, but less so in the Caesalpinioideae (Allen and Allen, 1981;Andrews and Andrews, 2017). Given the correspondence between increased crown size and rhizobia occurrence, the simplest hypothesis is that the increased availability of photosynthate in N-fixers enables more carbon allocation to crown mass and hence crown width. Alternatively, even if rhizobia do not affect overall per-tree productivity, they might allow N-fixers to differentially allocate more carbon to crowns than roots. Finally, some evidence suggests that root symbionts may stimulate increased leaf-level photosynthesis through their role as carbon sinks (Kaschuk et al., 2009). Could such a sink-effect be reflected on a whole-plant scale? If crown sizes are matched to the metabolic requirements of the organism, and if these requirements grow as a result of root symbioses, then we might expect this pattern of larger crowns in species that have root symbioses. This question would be an interesting avenue for future research.

Large crowns may be advantageous by increasing carbon uptake due to increased leaf area and light interception, but also by suppressing neighboring crowns. Taylor et al. (2017) found that N-fixing trees inhibited growth of their neighbors and of plots where they were abundant. By harboring large crowns, these N-fixers may be shading out their neighbors.

Does crown scaling differ between species? Iida et al. (2011) found that crown scaling parameters did not differ more than expected from the community tendency in Pasoh, Malaysia. Here we find that scaling in different forest types does differ (e.g., Figure 2.d), and that species differ as well (e.g. Figure S 18). Indeed, there was a clear phylogenetic signal in allometric scaling parameters (Figure 4).

While foresters have been well aware of species-specific crown shape since the inception of their practice (Larson, 1963), the relationship between phylogeny and crown shape has remained largely unexplored until now. Incorporating more studies across an even wider range of ecosystems and environments into larger phylogenies would be likely to yield interesting insights into the evolutionary pressures on the shapes of tree crowns.

### 5.3 ECOSYSTEM, BIOGEOGRAPHY, AND PHYLOGENY

Our community-wide hypothesis that crown depth decreases with elevation (increasing moving downslope) in our Peruvian transect was based on Horn’s (1971) individual-level theory. Thus, while the hypothesis was partially confirmed, a number of mechanisms might underlie the observed pattern, of which we discuss four. First, “faster” ecosystems in the lowlands (see NPP & GPP in Malhi et al 2016, submitted) could favor faster growing, acquisitive-strategy trees which, if Horn’s (1971) relationships hold, would have deeper crowns. Second, if tree height is linked to crown depth, the taller trees in the lowlands would lead to deeper crowns there. Third, greater solar radiation in the lowlands may allow for more shaded leaves to maintain positive carbon balance. Finally, increasing canopy heterogeneity could allow lateral light to penetrate deeper into the canopy, allowing lower leaves to maintain a positive carbon balance, and thus enabling deeper crown shapes. Heterogeneity may result from differences in heights of dominant trees, from topography, or a combination.

The last two mechanisms are not supported by the data: neither solar radiation (Table S 1) nor canopy heterogeneity (Asner et al., 2014; Table S 1) reflect the pattern in crown depth. The second mechanism, tree height, is indeed closely coupled with both relative and absolute crown depth. It seems clear that the driver of increasing crown depths moving downslope is the lengthening of height allometries from the uplands to the lowlands. This proximate “allometric” driver does not exclude the first mechanism, ecosystem productivity, as a more distal driver. Ecosystem productivity also shows a similar pattern to crown depth, and may be linked tree height allometries. Indeed, NPP and mean vegetation height exhibit similar patterns across the Peruvian transect (Malhi et al., 2017; Table S 1).

Crown depth exhibited a consistent response to ecosystem type. This suggests that crown depth is adaptive, but given that H7 (drought-depth hypothesis) was not supported, precipitation is not the primary factor driving it. We examined height – DBH allometries and crown depth comparisons for growth form differences, and found support for hypothesis H6 (Open growth form). Thus, savanna trees exhibit open-growth forms: they are ultimately shorter and harbor shallower crowns than forest trees of equivalent girth, but have deeper crowns than forest trees of equivalent height.

Biogeography has little effect on crown shape according to our analyses, in contrast to Moncrieff et al. (2014). Thus, we should expect neither deep phylogenetic patterns in nor strong influence of disparate faunal communities on crown shape. Rather, crown shape is likely more structured ecologically and competitively, and if evolution does play a role, we might expect crown architecture to be relatively plastic evolutionarily.

## 6 CONCLUSIONS

In this first look at tropical tree crown allometry across ecosystems, biogeography, and phylogeny, we find that both the lack of patterns in some instances (radius predictions across ecosystems and biogeography) and strong patterns in others (depth predictions across ecosystems, leguminous crown size) spurs further questions. In particular, ecological patterns such as ontogenetic variation and competitive effects on crown size may explain some of the observed patterns. Plasticity of crown dimensions in relation to local competition will be important to quantify for models to effectively simulate local dynamics, and this should comprise future research as well.

Our study lacks data from the Asian tropics. Thus, while we did not find a strong biogeographic signal in crown allometry, future studies that do include them and their especially tall trees may find one.

## 7 ACKNOWLEDGEMENTS

This work is a product of the Global Ecosystems Monitoring (GEM) network (gem.tropicalforests.ox.ac.uk) the Andes Biodiversity and Ecosystems Research Group ABERG (andesresearch.org), the Amazon Forest Inventory Network RAINFOR (www.rainfor.org), and the Carnegie Spectranomics Project (CSP; spectranomics.ciw.edu) research consortia. The field campaign was funded by grants to Y.M. from the UK Natural Environment Research Council (Grant NE/J023418/1), and from a European Research Council Advanced Investigator grant GEM-TRAITS (321131). G.P.A. and the Carnegie team were supported by the endowment of the Carnegie Institution for Science and a grant from the National Science Foundation (DEB-1146206). A.F.S. and L.P.B. were partially supported by John Fell Fellowships from the University of Oxford. I.O was supported by a Marie Curie Intra-European Fellowship. B.B. was supported by a NSF doctoral dissertation improvement grant and a NERC independent research fellowship. B.S.M. and B.H.M. were partially supported by CNPq research productivity grants (Pesquisador PQ). We thank the Servicio Nacional de Áreas Naturales Protegidas por el Estado (SERNANP) and personnel of Manu and Tambopata National Parks for logistical assistance and permission to work in the protected areas in Peru. We also thank the Explorers’ Inn and the Pontifical Catholic University of Peru, as well as ACCA for use of the Tambopata and Wayqecha Research Stations, respectively. We are indebted to Professor Eric Cosio (Pontifical Catholic University of Peru) for assistance with research permissions and sample storage. Importantly, we are grateful to all fieldworkers (especially tree climbers) who participated in the field campaigns. We are grateful to Stephanie A. Bohlman for advice on study design, and Beisit L. Puma Vilca for manuscript feedback. Brazilian co-authors are grateful to the National Council of Science and Technology of Brazil (CNPq) for financial support for project CNPq/PELD sítio 15 (403725/2012-7), FAPEMAT (164131/2013), and project CNPq/PPBio Phytogeography of Amazonia/Cerrado Transition (457602/2012-0), coordinated by B.S.M. and B.H.M., respectively.

