## Appendices for "The Influence of Ecosystem and Phylogeny on Tropical Tree Crown Size and Shape"

### SUPPLEMENTARY INFORMATION

#### 1 REVIEW OF CROWN ALLOMETRY THEORIES

##### 1.1 PREDICTIONS OF MST

Metabolic Scaling Theory (MST; West et al., 1997) offered an alternative explanation for crown size. Based on the hydraulics of vascular architecture (West et al., 1997), MST is a framework that explains how metabolic rate varies with body size (Kleiber's Law; Kleiber, 1932), and makes general predictions about the relationships between body size, organ size, and the rates of biological processes (West et al., 1999;Savage et al., 2008). Extensions to the theory predict the relationships between tree size and crown size (Enquist et al., 2009;West et al., 2009).

Scaling relationships take the following general form:

$$D = \beta\, I^{\alpha}\text{, or}$$

$$D \propto I^{\alpha}$$

where $D$ is the dependent variable, $I$ the independent predictor, β the normalization term, and α the scaling exponent. Most MST theory focuses on the value of α. Extending MST beyond metabolic and biomass predictions, West et al. (2009) derive a scaling relationship between stem radius and crown radius, surface area, and volume based on three assumptions (see Hypotheses).

Tests of MST crown scaling to date have been confined to individual sites, or when tested across sites, have been confined to the same ecosystem type, and have resulted in equivocal support. Enquist et al. (2009) found a canopy radius scaling exponent of 0.684 (95% CI 0.6457–0.7195, MST prediction is 2/3) using Reduced Major Axis regression (equation: $r_{crown} = 59.43\ D^{0.684}$) in a forest in Costa Rica. Muller Landau et al. (2006) found a crown area scaling exponent of 1.19 (95% CI 1.09–1.29; whether ground-projected or surface area is unclear; MST prediction is 4/3 for either) for Barro Colorado Island, while Farrior et al. (2016) found the projected crown area scaling exponent to be 1.28 for the same forest. Pretzsch and Dieler (2012) found intra-specific crown cross sectional area scaling exponents of 1.46 (95% CI 1.40 to 1.52) for all species, and 1.41 (95% CI 1.30 - 1.52) for angiosperms (MST prediction is 4/3), in forest plots in Germany, though inter-specific exponents fit MST less well. Antin et al. (2013) found good inter-specific concordance with MST scaling for crown radius, projected area, and volume in an Indian monsoon rainforest. Blanchard et al. (2016) found that 95% confidence intervals of projected crown area scaling exponents across pan-tropical lowland rainforest sites all contain the MST-predicted 4/3.

##### 1.2 SPACE, ECOSYSTEM, BIOGEOGRAPHY, AND EVOLUTION

Lateral competition may constrain crown widths within sites (Takahashi, 1996;Iida et al., 2011;Dieler and Pretzsch, 2013) (but see Rouvinen and Kuuluvainen, 1997), leading to similar widths for similarly-sized trees. This "competitive convergence" hypothesis posits that crown shape does not respond adaptively, but is only squeezed more or less depending on how close neighboring trees are packed together. Stem densities in our sites vary an order of magnitude, from 178 stems/ha in a Ghanaian savanna to 1627 stems/ha in a transitional Brazilian forest.

If crown allometries are adaptive, on the other hand, then we would expect to observe niche partitioning (Poorter et al., 2006) and therefore phylogenetic signal. Since a number of traits potentially related to crown dimensions are indeed phylogenetically structured (e.g. wood anatomy), we expect to observe a phylogenetic signal in crown size and shape (H4 (Phylogenetic signal)).

##### *1.2.1 ECOSYSTEMS AND BIOGEOGRAPHIC REGIONS*

Research on the influence of ecosystem type on crown shape is sparse. Examining crown shapes of *Acacia karroo* across savannas, forests, and arid shrublands in South Africa, Archibald and Bond (2003) found that trees in a fire-affected savanna grow tall before branching, leading to thin and deep crowns. Crown shapes across biogeographic regions are similarly understudied. Blanchard et al. (2016) examined crown allometry across the wet tropics and found little biogeographically-structured variation in scaling exponents. Examining savanna trees across Australia and Africa, Moncrieff et al. (2014) found differences in crown shape related more to biogeographic region than environmental variation across those regions. Underlying this biogeographic signal was a signal from evolutionary history driven by a few important taxa: *Vachellia* and *Senegalia* in South Africa (both belonging to the Mimosoid Fabaceae sub-family) versus *Eucalyptus* and *Corymbia* (both belonging to the Myrtaceae family) in Australia. We examine the variation of crown allometric scaling across biogreographic regions, and while we expect scaling to vary between regions (H5 (Biogeography)), we do not hypothesize a direction.

Across our three environmental gradients, we pose non-null ecological hypotheses that do not depend on competitive explanations.

###### 1.2.1.1 Forest-savanna transitions

Savannas often host fewer and more loosely packed trees than forests. “Open” growth form trees are characterized as being shorter and having wider, deeper, and more hemispherical crowns than trees of equivalent girth in the closed forest (Hallé et al., 1978). Thus while both the neutral competitive convergence and growth form hypotheses predict inverse relationships between stem density and crown widths, only the growth form hypothesis predicts deeper savanna crowns and shorter trees (H6 (Open growth form)).

Horn (1971) hypothesized that deep crowns confer drought tolerance due to self-shading of leaves. Therefore, when controlling for tree size, we expect that trees in drier sites will have deeper crowns than those in wetter sites (H7 (Depth drought tolerance)).

###### 1.2.1.2 Elevation gradient

Many characteristics of plant communities change across elevation gradients (Malhi et al., 2010;Feakins et al., 2016). We focus on shifts in productivity, though we examine the influence of shifts in height allometry below as well.

Ecosystems become more conservative with increasing elevation. As one moves upslope, most aspects of NPP decrease (total, stem, canopy, and fineroot; Girardin et al., 2010;Malhi et al., 2017), and leaf traits involved in the leaf economics spectrum (Wright et al., 2004) such as leaf mass per area (LMA) become more conservative (Feakins et al., 2016). Thus, more conservative life history strategies may be adaptive with increasing elevation. Horn's (1971) framework posits that fast growing trees should have deeper crowns and lower density wood than slower shade-tolerants. We therefore might expect slower, more conservative ecosystems to feature shallower crowns when compared to more dynamic ecosystems. Thus, across an elevation transect, this logic predicts narrower and deeper crowns in more productive lowland sites, versus wider and shallower crowns in colder, cloudier, and less dynamic upper montane sites (H8 (Ecosystem speed)). We note that while empirical data show that tropical pioneers tend to have shallow crowns (e.g., Poorter et al., 2006), the general relationship between crown dimensions and growth rate has not been conclusively determined. We therefore base our hypothesis on Horn's (1971) theory.

### 2 DETAILED RESULTS

#### 2.1 OBSERVED TRENDS ACROSS ECOSYSTEMS AND BIOGEOGRAPHIC REGIONS

Examining observed crown dimensions across ecosystems and regions, we find that crowns in the cloud zone of the Andes and in the savannas of Brazil are relatively shallow, while lowland forests in Peru, Brazil, and across all of Ghana (including the savanna) are relatively deep. Crowns in the Brazilian savanna and transitional plots are small, though their shapes (relative depths) fall within the ranges of the other sampled plots (Figure 5). Crowns in the Brazilian savanna are orders of magnitude smaller than those in Ghana and Peru; less so but still small when accounting for tree size differences (Figure 6).

Crown volume allometry does differ between sites: trees in ESP01 have relatively voluminous crowns for a given tree size, whereas those in the Brazilian savannas (CRP01 and NXV01) are small. However, we find that the large crowns we see in some Ghanaian plots (KOG04 and KOG01) are partially due to the presence of large trees, and not entirely to allometric differences (Figure 2d, Figure 6, Figure S 31).

Crown widths in the transitional plot increased the slowest with stem radius, and depth there increased the fastest, likely due high stem densities (Figure 7, Table S 1). Crowns in TMCF occupied a region between the transitional and TF ecosystems in terms of volume and surface area (Figure 7). Relative crown depth decreased with tree size across all ecosystems (Figure S 29).

#### 2.2 OBSERVED TRENDS ACROSS GRADIENTS WITHIN REGIONS

Here we examine and describe patterns across environmental gradients within each region. We examine both the structure of the sampled data (Figure 5) and fitted model parameters and predictions that account for tree size (Figure 6, others; see Allometric Scaling of Tree Crowns for model description). Hypotheses related to these patterns are addressed in the main text (see Environmental Gradient Hypotheses).

##### *2.2.1 PERU*

In Peru, moving downslope from the Andes to the Amazon, crown volumes and depths exhibit a step increase in the foothills between PAN03 and PAN02. The crowns in lower-elevation plots are larger and deeper both overall (Figure 5), and when controlling for tree girth and height (Figure 6), though these patterns fade for larger tree sizes (Figure S 31). Trees of the same girth are taller in the lowlands than the highlands (Figure S 3), and when local tree height allometries are used to set the local height in the crown model rather than equalizing height across sites, the depth signal reappears (Figure 6, Figure S 31). Thus, crown depth trends are largely a reflection of stem allometry trends across the elevation gradient.

Crown widths as observed do not exhibit strong trends with elevation (Figure 5). When accounting for stem radius however, a slight trend for 10cm DBH trees (Figure S 31a) and a step change for 30cm trees (Figure 6) at the cloud base between SPD-02 and PAN-03 is apparent, with crown width constant or increasing downslope until SPD-02, and dipping in the lowlands (Figure 6, Figure S 31). This trend disappeared for larger trees however (Figure S 31b,c), and ecosystem predictors were not significant in overall width models (see Trends across ecosystems and biogeographic regions above). Crown depth exhibits a step increase going downwards through the foothills between PAN03 and PAN02 (Figure 5). This step in crown depth is retained when accounting for stem radius (Figure S 32), but disappears when also accounting for tree height (Figure 6, Figure S 31). Tree height allometries mirror this pattern of elongation in the lowlands (Figure S 33), suggesting that changing height allometry is responsible for the increased lowland crown depths.

These patterns in crown width and depth result in a step change of relative crown depth (depth/width) from relatively flat crowns upslope to relatively deep crowns downslope as one emerges from the cloud base between SPD01 and PAN03. This pattern holds in both observed data (above) and in models that account for tree size (Figure 5, Figure S 26, Figure S 28), and is accentuated when local height-DBH allometries are included in the predictions (Figure 6, Figure S 31). Relative crown depth of smaller trees in lowland sites is more variable than in upland sites (Figure 6, Figure S 28, Figure S 31). Moving downslope, absolute depth increases earlier, at a slightly higher elevation, than relative depth due to unusual crowns in PAN03: they are relatively small, particularly narrow, and while crown depths there are comparable to other high-elevation sites, their relative depths resemble those of lowland plots. PAN03 sits on a wind-exposed ridge (Figure S ), with nutrient-poor white sand and clay soils derived from quartzite. Crowns may be narrow there due exposure or soils. Anecdotal observation suggests that crowns from our lowland sites in Peru (TAM05 and TAM06) may be shallower than those in more central Amazonian regions (Chavana-Bryant pers comm).

##### *2.2.2 BRAZIL*

In Brazil, moving from the savanna through the forest, crowns become larger as width and depth increase monotonically (Figure 5). These trends largely disappear when accounting for tree size with two exceptions: the true savanna (CRP01, *cerrado rupestre*) exhibits narrow crowns for small trees and the transitional site (NXV02, *cerradão*) exhibits narrow crowns for large trees (Figure 6, Figure S 31). Both of these sites have high stem counts, which likely accounts for their narrowness.

Relative depth of crowns in Brazil exhibits a steep step increase between the savanna and transitional/forest sites, even when accounting for tree size (Figure S 26, Figure S 28). Crowns are very shallow in the savanna plots, and are similar to those in the tropical montane forests of Peru in that regard, though they are by far the smallest crowns in the entire pan-tropical study.

##### *2.2.3 GHANA*

In Ghana, the savanna crowns are exceedingly large, dwarfing even the crowns of the forest trees in Brazil and much of Peru (Figure 5, Figure 6). Crowns shrink as one moves from the savanna to the transitional site of BOB01, increasing again towards the closed-canopy forest of ANK03. Crowns grow deeper moving from the savanna to the forest, and this drives the observed patterns in crown volume; crown width remains largely constant. BOB01 stands out as an exceptional site with particularly small crowns, especially when tree size is accounted for (Figure 6), perhaps linked to heavy liana infestation in that plot. Across all of Ghana, the relative depth of crowns remains largely constant (Figure 5), though the savanna (KOG04) and relatively stem-dense semi-deciduous forest (BOB02) harbor relatively deep crowns per tree size (Figure S 28).

Crown volume distributions are skewed more to the right (towards larger volumes) than the other dimensions. The skewness increases with decreasing elevation, such that the largest crowns become larger as one moves downslope. Crown depth is also skewed to the right in the Peruvian Andean sites and the Brazilian *cerrado rupestre* (CRP01). Crown relative depth is accordingly skewed to the right in the Andes, but not so in in the *cerrado rupestre*.

##### 2.3 Observations across regions and ecosystems

Crown radius scaling changes significantly across sites (Figure 2a; Appendices). Sites across the Peru transect generally exhibit shallower crown-stem scaling than in Ghana and Brazil. While Peruvian mid-elevation sites exhibited steeper scaling than high- and low-elevation sites, we do not find any strong or consistent trends in $\alpha_{rad}$ across elevation in Peru. The savanna sites exhibited steeper scaling than the transitional and tropical forest sites in Brazil, but no such savanna-forest difference was observed in Ghana.

##### 2.4 Scaling model formulation

We tested three models for predicting crown width scaling exponents, each with different covariates: (1) no site covariate (Model 5), (2) site exponential covariates ($\alpha_{site}$; Model 6), and (3) both site exponential and normalization covariates ($\beta_{site}$; Model 7). All models contain a random exponential ($\alpha_{spp}$) and normalization ($\beta_{spp}$) term per species. While Model 6 was deemed the most parsimonious model by corrected-AIC ($\Delta AICc_{2-1} = 4.21$), likelihood ratio tests (LRT) indicate that inclusion of site normalization ($\beta_{site}$) significantly improves the model (Table S 5). Furthermore, because removal of model terms based on their significance is a subject of current debate (Bolker et al., 2009), we consider the inclusion of site normalization terms to constitute a conservative choice. We therefore use Model 7 from here onwards. Because the fixed parameter estimates for α, β, and per-site deviations did not vary appreciably between the 3 models, we have some confidence in the robustness of our results.

###### *2.4.1 Models for crown depth*

Nonlinear mixed models with just tree height (1) were more parsimonious than those with just stem radius (2) or both stem radius and tree height (3; $\Delta AICc_{1-2} = 164$, $\Delta AICc_{1-3} = 181$; Model 11), though model (3) was reduced in complexity to converge (no per-site nor random normalization effects). Formulated as LMMs, model (3) without reduced complexity was the most parsimonious ($\Delta AICc_{3-1} = 18$, $\Delta AICc_{3-2} = 215$; Model 12), and every term significantly improved the model (LRT; $P(\gamma_j) = 0.04$, $P(\alpha_{depth}) = 0.0001$, $P(\alpha_{depth,h}) < 0.0001$, $P(\delta_j) = 0.0004$, $P(\theta_j) = 0.0002$). Despite the unusual formulation (see Appendices), we focus on the best LMM model (3; Model 12) with tree height and stem radius as the predictors of crown depth.

#### 2.5 MST Distal Assumptions

The first assumption of MST crown scaling (West et al., 2009) is that stem diameter and height are related via elastic similarity (Eq. 1; McMahon and Kronauer, 1976). Our data, however, support a relationship based on the constant stress theory prediction of $\alpha_h = 1/2$ (Model 2; $\alpha_h = 0.46$, 95% CI 0.42 – 0.51; Figure S 3.). Thus, H2 (MST distal 1) is not supported.

The second assumption of MST crown scaling (West et al., 2009) is that crown dimensions scale isometrically with tree height (Eq. 2, $r_{crown} \propto h^{\alpha_{rad,h}}$, $\alpha_{rad,h} = 1$). Our data show that crown radii vary nearly an order of magnitude for any particular tree height (Figure S 4, Figure S 5). For the scaling equation, the LMM fit (Model 3) results in $\alpha_{rad,h} = 0.63$ (CI 0.53 – 0.73, $R^2_{LMM(m)} = 0.45$, $R^2_{LMM(c)} = 0.66$), or $\alpha_{rad,h} = 0.54$ (CI 0.46 – 0.61, $R^2_{LMM(m)} = 0.31$, $R^2_{LMM(c)} = 0.66$) with site covariates removed. These scaling exponents are significantly less than the MST assumption of $\alpha_{rad,h} = 1$ (Table 3, Figure S 5), and do not support H3 (MST distal 2).

Our models consistently found a strong relationship between $r_{crown}$and $r_{stem}$ independent of $h$. Therefore, we found it reasonable to modify this second assumption (Eq. 2) to include $r_{stem}$ (Model 4; see Appendices for derivation). Thus, the modified assumption becomes:

$$r_{crown} = \beta_{rad,h} h^{\alpha_{rad,h}} + \beta_{rad} {r_{stem}}^{\alpha_{rad}}$$

Site was included as a random effect because models with fixed site effects failed to converge (fit parameter values were very similar between fixed site, random site, and no site models regardless). This modification resulted in $\alpha_{rad,h} = 0.16$ (SE = 0.05) and $\alpha_{rad} = 0.59$ (SE = 0.038; $R^2_{LMM(m)} = 0.50$, $R^2_{LMM(c)} = 0.63$; profiling & bootstrap for CIs failed). We include parameters for the same model but with site terms removed as we were able to compute confidence intervals via profiling methods for it: $\alpha_{rad,h} = 0.16$ (95% CI 0.085 – 0.238) and $\alpha_{rad} = 0.52$ (95% CI 0.524 – 0.664). We discuss the motivation for and implications of this modified assumption below.

#### 2.6 Resolving the incongruence of MST distal assumption deviation and MST proximate prediction accuracy

We abbreviate our notation here: $r_s \coloneqq r_{stem}$, $r_c \coloneqq r_{crown}$, $\beta_c \coloneqq \beta_{rad,h}$, $\alpha_c \coloneqq \alpha_{rad,h}$, and $\alpha_{cr_s} \coloneqq \alpha_{rad}$.

The first assumption is that tree height is related to stem radius by elastic similarity, or by a scaling exponent of 2/3. Hence,

$$h = \beta_h {r_s}^{\alpha_h} \qquad (1)$$

where $\alpha_h = 2/3$.

The second assumption is that crown radius scales isometrically with tree height:

$$r_c = \beta_c h^{\alpha_c} \qquad (2)$$

where $\alpha_c = 1$.

The third assumption, which we do not examine in depth here, is that the shape of the crown is Euclidean.

Substituting gives

$$\begin{aligned} r_c &= \beta_c (\beta_h r_s{}^{\alpha_h})^{\alpha_c} \\ &= \beta_c \beta_h{}^{\alpha_c} r_s{}^{\alpha_h \alpha_c} \end{aligned} \qquad (3)$$

The proximate MST prediction for crown radius derives from equations 1 and 2. If,

$$r_c = \beta_{cr_s} r_s{}^{\alpha_{crs}}$$

then,

$$\beta_{cr_s} = \beta_c \beta_h$$

$$\alpha_{cr_s} = \alpha_h \alpha_c \qquad (4)$$

Empricially,

$$\alpha_h = 0.46$$

$$\alpha_c = 0.54$$

$$\alpha_{cr_s} = 0.67$$

$\alpha_h$ and $\alpha_c$ lie notably below their predicted values, while $\alpha_{cr_s}$ is predicted well by MST. The incongruence arises in equation 4 since,

$$0.46 \times 0.54 = 0.25 \neq 0.67$$

To attempt to resolve this problem, we investigate whether $\beta_c$ is itself a function of $r_s$. In other words, if variation in $r_s$ for a given $h$ influences $r_c$ significantly, then the assumptions as formulated in equations 1 and 2 may not constitute a sufficient basis for prediction.

Thus, if

$$\beta_c = \beta_c{}' r_s{}^{\alpha_c{}'}$$

Then equation 2 becomes

$$r_c = \beta_c{}' r_s{}^{\alpha_c{}'} h^{\alpha_c} \qquad (5)$$

Then from equation (3),

$$\begin{aligned} r_c &= \beta_c \beta_h{}^{\alpha_c} r_s{}^{\alpha_h \alpha_c} \\ &= \beta_c{}' r_h{}^{\alpha_c{}'} \beta_h{}^{\alpha_c} r_s{}^{\alpha_h \alpha_c} \end{aligned}$$

$$= \beta_c{}'\beta_h{}^{\alpha_c} r_s{}^{\alpha_h \alpha_c + \alpha_c{}'}$$

Our new equivalence for $\alpha_{cr_s}$, from equation 4, becomes

$$\alpha_{cr_s} = \alpha_h \alpha_c + \alpha_c{}' \qquad (6)$$

To determine empirical values for $\alpha_c$ and $\alpha_c{}'$ from equation 5, we fit the following LMM:

$$\log(r_c) = \log(\beta_c{}') + \alpha_c{}' \log(r_s) + \alpha_c \log(h)$$

We also include species-level random intercepts as and slopes across both log(h) and log(rc) (in lme4 this is specified as `(1 + log10(h_tree)|Species) + (1 + log10(r_stem)|Species)`). This model results in the following empirical scaling exponents:

$$\alpha_c = 0.16$$

$$\alpha_c{}' = 0.59$$

We can evaluate whether the exponents of the new theoretical assumptions correspond with empirical observations by substituting empirical values in equation 6. Thus, empirically,

$$0.67 \cong 0.46 \times 0.16 + 0.59 = 0.66$$

We find that by including the dependence of crown radius on stem radius, in addition to height, our new assumptions are able to explain the empirical observations of crown radius scaling.

### 3 SMA REGRESSION RESULTS

SMA regressions were performed with the smart R package (Warton et al., 2012). Mean SMA factor estimates were computed by taking factor means across the grouping variables (site, species, genus, or family) and computing confidence intervals as 1.96 times the standard deviation of those means.

Testing isometric scaling of crown radius with tree height (i.e. $\alpha_{rad,h} = 1$), the SMA fit yields $\alpha_{rad,h} = 0.56$ (95% CI -2.86 – 3.98). SMA regressions and LMM fits agree on the slope of the relationship, but differ in their confidence intervals (Table 3).

#### 3.1 CROWN WIDTH SCALING

When aggregated by species (intraspecific scaling), genus, or family, crown radii are largely as predicted by MST by SMA regressions (aggregated by: species, $\alpha_{radius} = 0.69$ (95% CI = 0.28 — 1.1), genus $\alpha_{radius} = 0.82$ (0.25 — 1.39), family $\alpha_{radius} = 1.14$ (0.45 — 1.82); Table 3. , Figure S 6., Figure S 7., Figure S 8.). When we remove the phylogenetic grouping in SMA models and thereby test interspecific scaling, SMA regressions result in the crown radius scaling exponent appearing larger than the $\alpha = 2/3$ predicted by MST ($\alpha_{radius} = 1.13$ (1.08 — 1.18); Table 3. and Figure S 10.). Because MST assumes key parameters such as LAD are constant across individuals, controlling for these parameters by aggregating phylogenetically provides a better estimate. SMA models support our conclusions based on Figure 2.a that crown radius scaling changes significantly across sites (SMA test of differing slopes $p < 0.001$).

#### 3.2 CROWN SURFACE AREA SCALING (PROXIMATE)

SMA results agree with LMM results for crown surface area scaling. Intra-specific, -generic, and –familial SMA regressions fit the MST prediction of $\alpha_{sa} = 4/3$ well (eq. 2, Figure S 11., Figure S 12., Figure S 13., Figure S 14., and Table 3. ). Interspecific SMA regressions again predict a much larger scaling exponent than MST, and are not in agreement with the intra-taxa models (Table 3. ).

#### 3.3 CROWN VOLUME SCALING (PROXIMATE)

SMA regressions again agree with LMM fits for crown volume scaling. Intra-specific, -generic, and –familial SMA regressions fitting $\alpha_{vol}$ confirm MST predictions (Eq. 3, Figure S 16., Figure S 17., Figure S 18., Figure S 19., and Table 3. ). Interspecific SMA regressions again predict a much larger scaling exponent than MST, and are not in agreement with intra-taxa models. Per-site deviations of $\alpha_{vol}$ from the MST prediction are again similar to those of $\alpha_{rad}$. Species-specific scaling fits vary significantly (Figure S 16., Figure S 18.).

#### 3.4 DERIVATION OF MST SCALING PREDICTIONS

Based on a set of eight fundamental assumptions relating to hydraulic architecture, West et al. (1997) derive the general result that an organism's metabolic rate, $B$, is related to body mass $M$ by the well-known Klieber's Law, $B = B_0 M^{\alpha}$, where $\alpha$=3/4. Extending this theory to tree crowns, West et al. (2009) first took McMahon and Kronauer's (1976) derivation of elastic similarity for the relationship between stem height and stem radius:

$$h_{stem} \propto r_{stem}^{2/3}$$

Note that Price et al. (2007) derive a more general relationship from self-similarity principles and by assuming constant wood density as $h_{stem} \propto r_{stem}^{b/a}$, where a = 1/2 and b = 1/3 for a tree conforming to four assumptions: volume-filling branch networks, linear hydrodynamic scaling, independence of leaf properties and tree size, and uniform biomechanical constraints. West et al. (2009) further assume that crown radius scales linearly with tree height:

$$r_{crown} \propto h$$

By further assuming a Euclidean crown, West et al. (2009) derived the relationship between crown and stem radii to be $r_{crown} \propto r_{stem}^{(2a+b)/a}$. While $a$ and $b$ are allowed to vary between species, for the ideal case they are set to $a = 1/2$ and $b = 1/3$ as above, and hence

$$r_{crown} = \beta_{rad} r_{stem}^{\alpha_{rad}}, \text{ where } \alpha_{rad} = {}^{2}/_{3}$$

where $\beta_{rad}$ is a normalization constant. Our notation will omit subscripts when referring to a property of an entire tree, and will include notation when referring to a tree part.

West et al. (2009) extend the predicted scaling from crown radius to crown surface area (sa). Surface area here refers to the area of the crown as a convex hull (i.e. as if wrapped in a plastic sheet). Because surface areas of Euclidean shapes scale as $r^2$ (e.g. $sa_{sphere} = 4\pi r^2$), the crown surface area should scale as

$$a_{crown} \propto r_{crown}^{2} \propto r_{stem}^{4/3}$$

*thus,* $$a_{crown} = \beta_{sa} r_{stem}^{\alpha_{sa}}, \textit{ where } \alpha_{sa} = {}^{4}/_{3}$$

The crown volume, referring to the volume of the polyhedron enclosed in that same plastic sheet, should then scale as

$$v_{crown} \propto r_{crown}^{3} \propto r_{stem}^{2}$$

*thus,* $$v_{crown} = \beta_{vol} r_{stem}^{\alpha_{vol}}, \textit{ where } \alpha_{vol} = 2$$

MST does not make specific predictions for crown depth. It does, however, make general assumptions about Euclidean crown shapes. These assumptions lead to the same prediction for crown depth as for crown radius: $\alpha_{depth} = 0.67$.

#### 4 Scaling Models and Formulation in R

Metabolic scaling models take the following general form:

$$D = \beta_0 + \beta_1 I^{\alpha},$$

where $D$ is the dependent variable, $I$ the independent variable, $\beta_1$the normalization coefficient, and $\alpha$ the exponential scaling factor. Log-transforming both sides of the equation yields a linear equation, which lends itself to linear models:

$$\log D = \log(\beta_0 + \beta_1 I^{\alpha})$$

The intercept, $\beta_0$, is often assumed to be zero. This assumption is mathematically expedient, and also makes some biological sense: a body size of zero should exhibit zero metabolism. This assumption leaves us with the following scaling formulation:

$$\log D = \log(\beta_1 I^{\alpha})$$
$$= \log \beta_1 + \alpha \cdot \log I$$

Therefore, in a linear model, we can interpret the intercept as the logged normalization coefficient and the slope as the exponential scaling factor. In the lm R function, this model is specified as:

```
lm(log(D) ~ log(I))
```

In order to investigate the role of covariates in the normalization and scaling factors, we recast our scaling model as follows:

$$D = \exp(\beta_c C)\, \beta_1 I^{(\alpha + \alpha_c C)}$$

where $C$ is the covariate, $\beta_c$ is the covariate coefficient for the influence of $C$ on the normalization coefficient, and $\alpha_c$ is the covariate coefficient for the additive influence of $C$ on the exponential scaling factor. The form of the normalization covariate, $\exp(\beta_c C)$, is uncommon and not easily interpretable, but does have the beneficial property of simplifying well in log-transformed models. Rank relationships will still be preserved by this formulation. Parameterizing a log-transformed model for additive (e.g. $\beta_1 + \beta_c C$) or multiplicative (e.g. $\beta_1 \cdot \beta_c C$) normalization covariates is not mathematically tractable, and non-linear models should be used if such relationships are deemed important. Log transforming as above results in:

$$\log \mathrm{D} = \log \beta_1 + \beta_c \mathrm{C} + (\alpha + \alpha_c \mathrm{C}) \cdot \log \mathrm{I} \qquad \text{(S1)}$$

The formulation of equation S1 allows parameters to be fit both for the overall scaling coefficients and the covariate coefficients. This model can be specified in R as:

```
lm(log(D) ~ C + log(I) + C:log(I))
```

The intercept term estimated by lm corresponds to $\log(\beta_1)$, the covariate associated with C to $\beta_c$, the covariate associated with log(I) to $\alpha$, and the covariate associated with C:log(I) to $\alpha_c$. When incorporating random effects into linear models via the lme4 R package (Bates et al., 2015), a similar formulation can be used. For a random exponential factor, the following formula can be used:

```
lmer(log(D) ~ (0 + log(I)|R) + log(I))
```

which corresponds to:

$$D = \beta_1 I^{(\alpha + \alpha_R R)}$$

where the coefficients associated with log(I)|R correspond to a random effect $\alpha_R$ term. For both random exponential and normalization factors, the formula becomes:

```
lmer(log(D) ~ (1 + log(I)|R) + log(I)),
```

which corresponds to:

$$D = \beta_1 \exp(\beta_R R)\, I^{(\alpha+\alpha_R R)},$$

where the coefficients associated with R correspond to a random effect $\beta_R$. Finally, we can combine random and fixed effects:

$$D = \beta_1 \exp(\beta_R R) \exp(\beta_C C)\, I^{(\alpha+\alpha_R R+\alpha_C C)} \quad \text{(S2)}$$

Equation S2 is implemented in lme4 as:

```
lmer(log(D) ~ (1 + log(I)|R) + C + log(I) + C:log(I))
```
(8)

Note that C + log(I) + C:log(I)is equivalent to C*log(I), but we present the formula in expanded form to emphasize that either C or C:log(I) can be removed. Equation (8) represents a scaling formula that is extensible by the addition of fixed and random variables as desired.

##### 4.1 R IMPLEMENTATION INSTRUCTIONS

In straightforward terms, when additive scaling exponent ($\alpha$) covariates are desired, one merely adds "+ C:log(I)" to the lme4 model, where C is the covariate of interest. To add multiplicative normalization ($\beta$) covariates, one adds "+ C" to the model. To add additive random effects to the scaling exponent, one adds "+ (0 + log(I)|R)" to the model where R is the random factor of interest; or "+ (1|R)" for multiplicative random normalization effect; or "+ (1 + log(I)|R)" for both exponential and normalization random effects.

##### 4.2 NONLINEAR MIXED MODELS

If we wish to model the normalization coefficients ($\beta_R R$, $\beta_C C$) without the exp( ) transformation, we must turn to nonlinear models. In this paper, we use nonlinear models when examining the role of functional traits in predicting tree crown allometric scaling parameters, since the exponential relationship the linear model forces is not likely to reflect reality. Adding trait covariates to the original scaling equation (eq S1) results in the following:

$$D = (\beta_1 + \beta_t T) I^{\alpha+\alpha_t T},$$

where T represents the trait value, $\beta_t$ the coefficient for the additive effect of the trait on the normalization parameter, and $\alpha_t$ the coefficient for the additive effect of the trait on the scaling parameter. To account for between-species variation as discussed in the main text, we add random effects:

$$D = (\beta_1 + \beta_R R + \beta_t T) I^{\alpha+\alpha_R R+\alpha_t T},$$

Finally, we implement this model in the nlme R package (Pinheiro et al., 2016) as follows:

```
trait_scaling_fun <-
  function(alpha, beta, beta_tau, alpha_tau, trait, ind_var) {
    return((tau_beta*trait + beta) * ind_var^(alpha + tau_alpha*trait))
  }
```

```
              nlme(crown_radius ~ trait_scaling_fun(alpha, beta,
alpha_tau, beta_tau, trait, stem_rad),
                data = trait_df,
                fixed = list(alpha ~ site, beta ~ site,
alpha_tau + beta_tau ~ 1), random = alpha + beta ~ 1|Species,
                start = list(fixed = c(rep(1, site_len), rep(1,
site_len), 1, 1)))
```

,where trait_df is the dataframe containing the trait and crown data with one row per tree, site is the column in trait_df representing the site where the tree is found, and site_len is the total number of sites in the dataframe.

Nonlinear models are useful for their flexibility, but are more statistically complex. Thus, there are no robust goodness-of-fit statistics for nonlinear mixed models as there are for linear mixed models (e.g. pseudo-R2; Nakagawa and Schielzeth, 2013;Johnson, 2014). Hence, we rely on corrected-AIC for model comparison, but go no further than this in quantifying model errors.

#### 4.3 Testing Mechanical Constraints

Models of mechanical stress and optimal tree design often assume that wood mechanical properties are constant (e.g. King and Loucks, 1978). We address this assumption by including per-species random intercepts and slopes in our LMMs, and as a grouping factor in our SMA regressions.

We fit the model height $= \beta \text{DBH}^{\alpha}$ and compared $\alpha$ with theoretical values for three mechanical constraint models: constant stress ($\alpha$ = 0.5), elastic similarity ($\alpha$ = 0.67), and geometric self-similarity ($\alpha$ = 1). Overall, trees conformed most closely to the constant stress model ($\alpha_{\text{grand mean}}$ = 0.47, 95% CI 0.42 – 0.51; Figure S 3.), though there was significant variation between plots (Figure S 3.). Plot-specific models show that most plots have trees that are shorter and squatter than the constant stress model; the most conservative mechanical model of the three.

#### 4.4 R Package Environment

```
> sessionInfo()
R version 3.3.2 (2016-10-31)
Platform: x86_64-w64-mingw32/x64 (64-bit)
Running under: Windows 7 x64 (build 7601) Service Pack 1

locale:
[1] LC_COLLATE=English_United States.1252      LC_CTYPE=English_United States.1252
      LC_MONETARY=English_United States.1252 LC_NUMERIC=C
[5] LC_TIME=English_United States.1252

attached base packages:
[1] grid      parallel      stats  graphics      grDevices utils      datasets
      methods      base

other attached packages:
 [1] tidyr_0.6.3    phytools_0.6-00     maps_3.2.0    ape_4.1       xtable_1.8-2
      ggtree_1.9.1  treeio_1.0.2
 [8] adephylo_1.1-10       ade4_1.7-6    ggthemes_3.4.0      sjPlot_2.3.1
      gridExtra_2.2.1     gdata_2.18.0  scales_0.4.1
```

```
[15] stargazer_5.2  coefplot2_0.1.3.3   coda_0.19-1  AICcmodavg_2.1-0    smatr_3.4-3
      gtools_3.5.0  gemtraits_0.1
[22] geoR_1.7-5.2   memoise_1.1.0 taxize_0.8.4  maptools_0.9-2      sp_1.2-4
      spatstat_1.51-0     rpart_4.1-11
[29] nlme_3.1-131   plyr_1.8.4   geometry_0.3-6      magic_1.5-6  abind_1.4-5
      piecewiseSEM_1.2.1  cowplot_0.7.0
[36] ggplot2_2.2.1  afex_0.16-1  lsmeans_2.26-3      estimability_1.2
      reshape2_1.4.2      lme4_1.1-13  Matrix_1.2-10
[43] phylobase_0.8.4       RPostgreSQL_0.4-1   DBI_0.6-1    XLConnect_0.2-13
      XLConnectJars_0.2-13

loaded via a namespace (and not attached):
  [1] uuid_0.1-2    fastmatch_1.1-0     blme_1.0-4   VGAM_1.0-3   igraph_1.0.1
      lazyeval_0.2.0
  [7] splines_3.3.2 unmarked_0.12-2     rncl_0.8.2   TH.data_1.0-8 digest_0.6.12
      foreach_1.4.3
 [13] htmltools_0.3.6      magrittr_1.5  RandomFieldsUtils_0.3.25 cluster_2.0.6
      tensor_1.5   modelr_0.1.0
 [19] gmodels_2.16.2       sandwich_2.3-4      prettyunits_1.0.2   colorspace_1.3-2
      haven_1.0.0  dplyr_0.7.0
 [25] tcltk_3.3.2   jsonlite_1.5  phangorn_2.2.0      survival_2.41-3     zoo_1.8-0
      iterators_1.0.8
 [31] glue_1.0.0    polyclip_1.6-1      gtable_0.2.0  MatrixModels_0.4-1  seqinr_3.3-6
      sjstats_0.10.0
 [37] sjmisc_2.5.0  car_2.1-4    SparseM_1.77  mvtnorm_1.0-6 Rcpp_0.12.11  plotrix_3.6-5
 [43] RandomFields_3.1.50  progress_1.1.2      merTools_0.3.0      foreign_0.8-68
      spdep_0.6-13  bold_0.4.0
 [49] stats4_3.3.2  DT_0.2 animation_2.5 htmlwidgets_0.8     httr_1.2.1   lavaan_0.5-
23.1097
 [55] modeltools_0.2-21    reshape_0.8.6 XML_3.98-1.7  rJava_0.9-8  nnet_7.3-12
      deldir_0.1-14
 [61] labeling_0.3  rlang_0.1.1  munsell_0.4.3 tools_3.3.2  splancs_2.01-40
      sjlabelled_1.0.0
 [67] broom_0.4.2   stringr_1.2.0 arm_1.9-3    goftest_1.1-1 knitr_1.16   purrr_0.2.2.2
 [73] coin_1.1-3    mime_0.5     quantreg_5.33 adegenet_2.0.1      xml2_1.1.1
      pbkrtest_0.4-7
 [79] spatstat.utils_1.6-0 clusterGeneration_1.3.4    tibble_1.3.3  pbivnorm_0.6.0
      RNeXML_2.0.7  stringi_1.1.5
 [85] lattice_0.20-35      psych_1.7.5  nloptr_1.0.4  permute_0.9-4 vegan_2.4-3
      effects_3.1-2
 [91] msm_1.6.4     stringdist_0.9.4.4  LearnBayes_2.15     combinat_0.0-8
      lmtest_0.9-35 data.table_1.10.4
 [97] raster_2.5-8  httpuv_1.3.3  R6_2.2.1     codetools_0.2-15    boot_1.3-19
      MASS_7.3-47
[103] assertthat_0.2.0     mnormt_1.5-5  multcomp_1.4-6      expm_0.999-2  mgcv_1.8-17
      quadprog_1.5-5
[109] minqa_1.2.4   rvcheck_0.0.8 scatterplot3d_0.3-40       numDeriv_2016.8-1
      shiny_1.0.3
```

#### 5 SI Figures

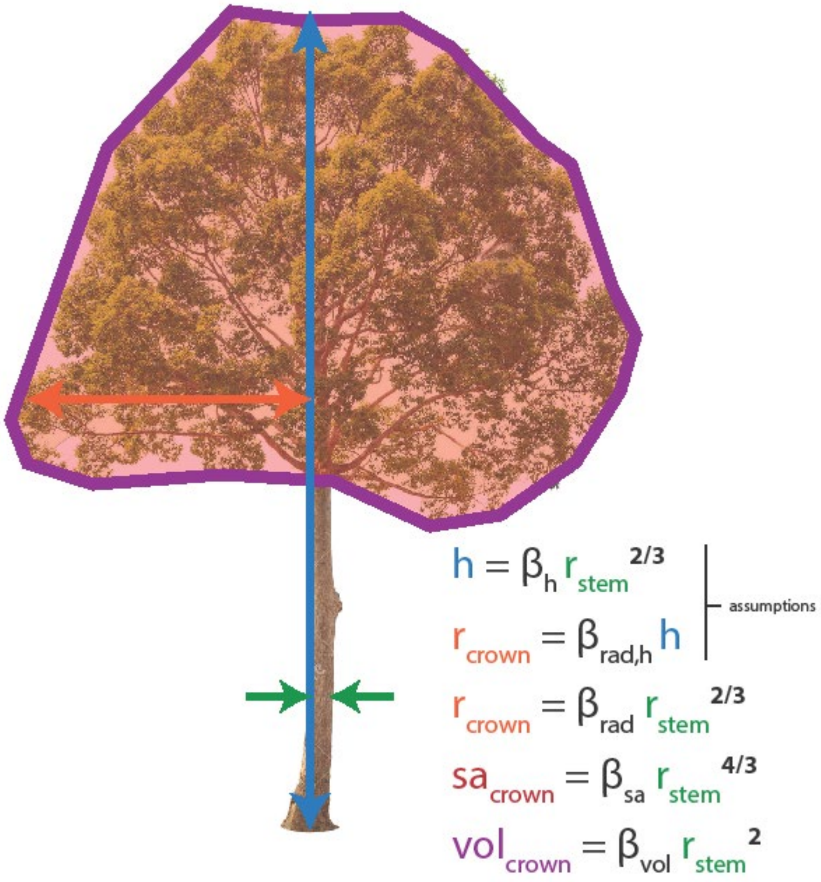


Figure S 1. Illustrated metabolic scaling equations for tree crowns.

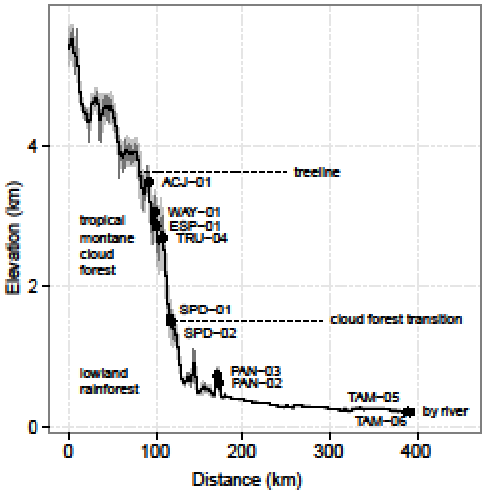


Figure S 2. Location of 10 CHAMBASA forest plots on an elevation transect across the eastern slope of the Andes in southern Peru. Site locations (open circles, site name annotated), vegetation zones, cloud base and river proximity are indicated. Elevation profile: gray line (elevation acquired from the Shuttle Radar Topographic Mission (SRTM) 90 m Digital Elevation Database 4.1 (Reuter et al., 2007), black line (smoothed elevation), gray envelope (topography; ±2r elevation from 1 km wide swath perpendicular to transect, re-centered to smoothed elevation). R code available from https://gist.github.com/ashenkin/7fceb77e78efc33961a8

#### 5.1 MST ASSUMPTION TESTS

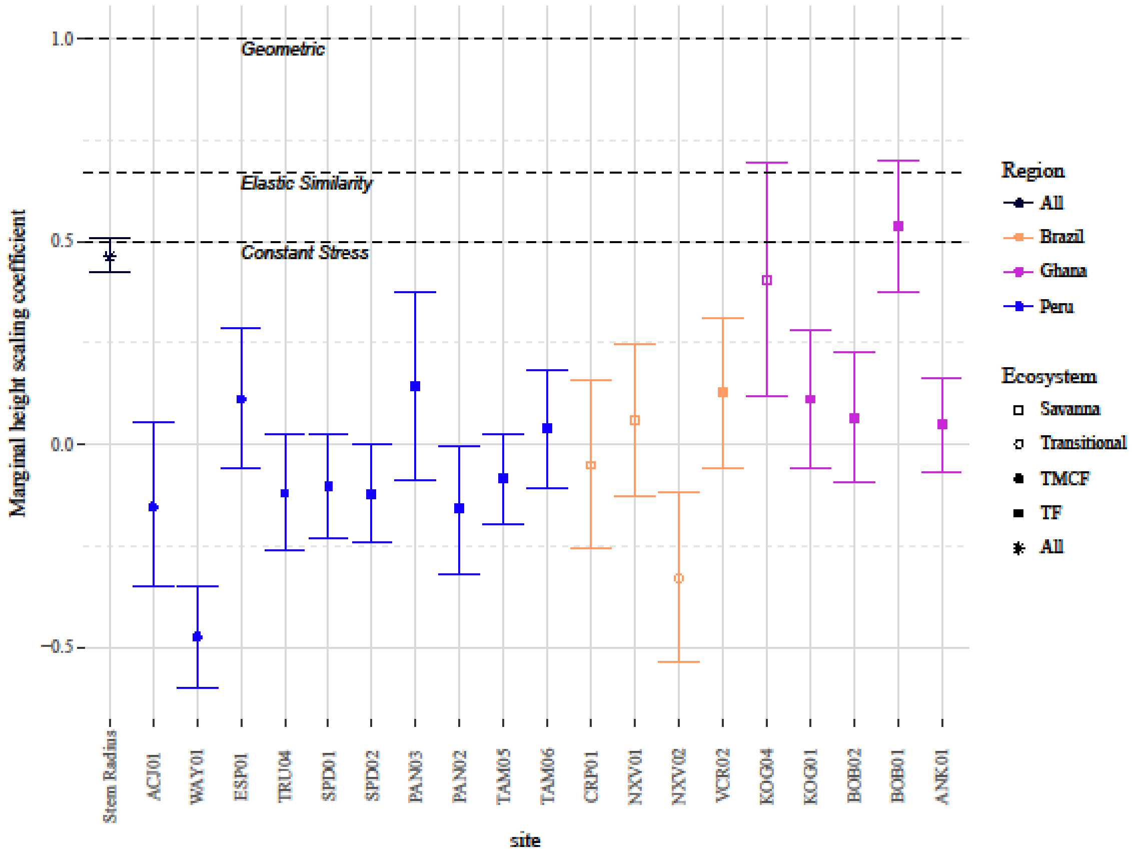


Figure S 3. Height – DBH scaling coefficients from the linear mixed effects model: $\log_{10}$(height) ~ N($\log_{10}$(DBH$_{grand}$ + site + site x DBH), $\sigma^2$). Values correspond to the $\alpha$ parameter in the model $height = \beta DBH^{\alpha}$. Higher coefficients indicate taller and slimmer trees, whereas lower coefficients indicate shorter and squatter trees. Stem_radius is the grand mean effect of DBH on tree height, and the site effects represent the effect of that site on the slope of relationship. Sites with higher values have taller trees for a given DBH, and vice versa. Because deviance contrasts were used, the site effects have a mean of zero, and should be interpreted as an addition to or subtraction from the overall height – DBH slope. Not shown are intercepts and species that were coded as random effects.

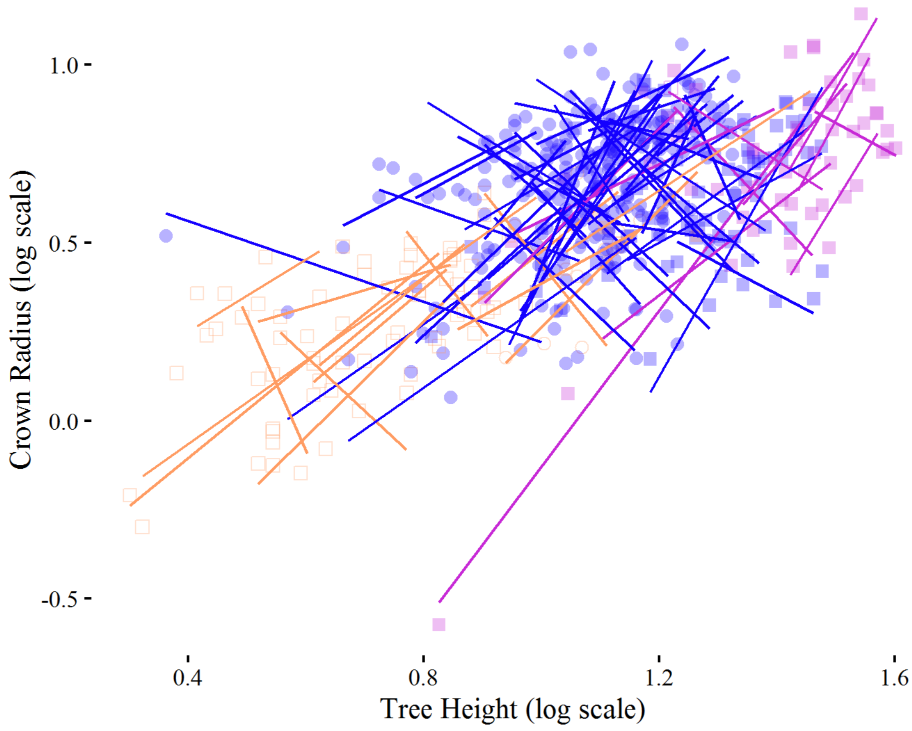


Figure S 4. Tree height versus crown radius. Lines are SMA per-species fits, points are individual observations.

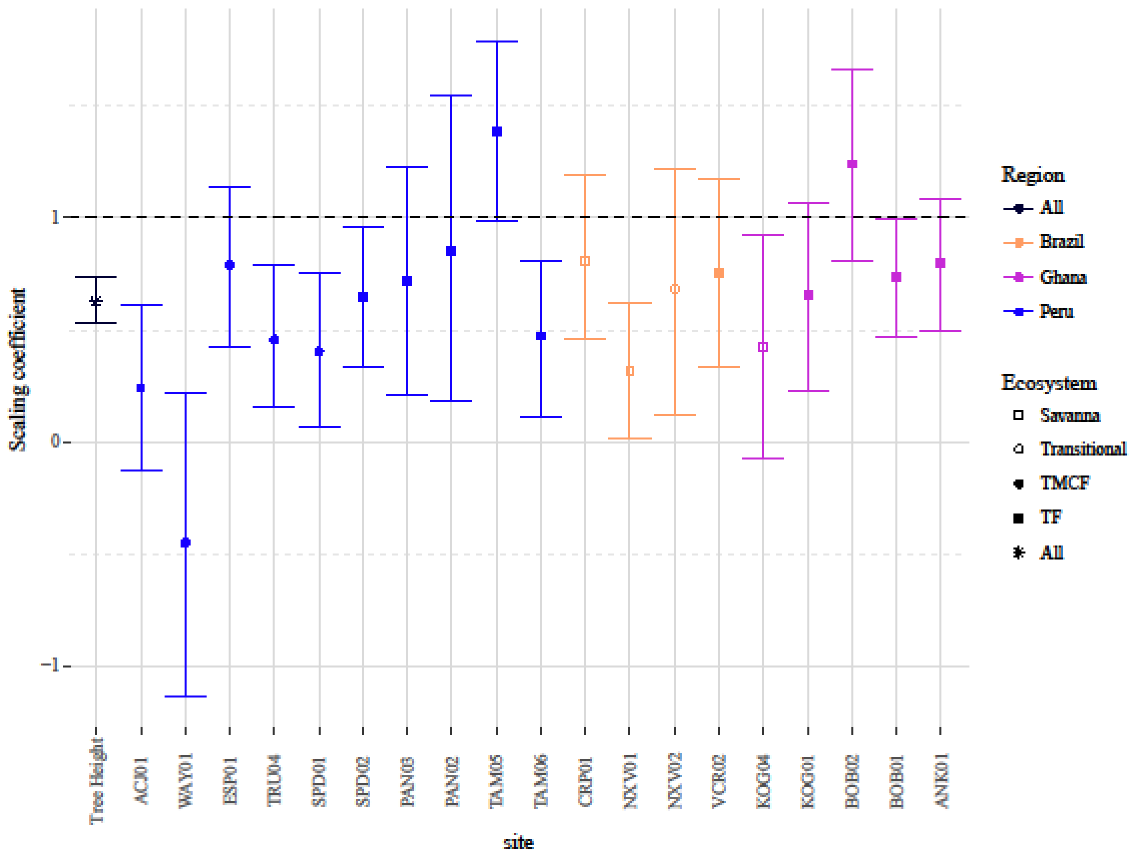


Figure S 5. Tree height versus crown radius slopes as fit by a LMM with species as a random effect.

#### 5.2 MST CROWN RADIUS SCALING

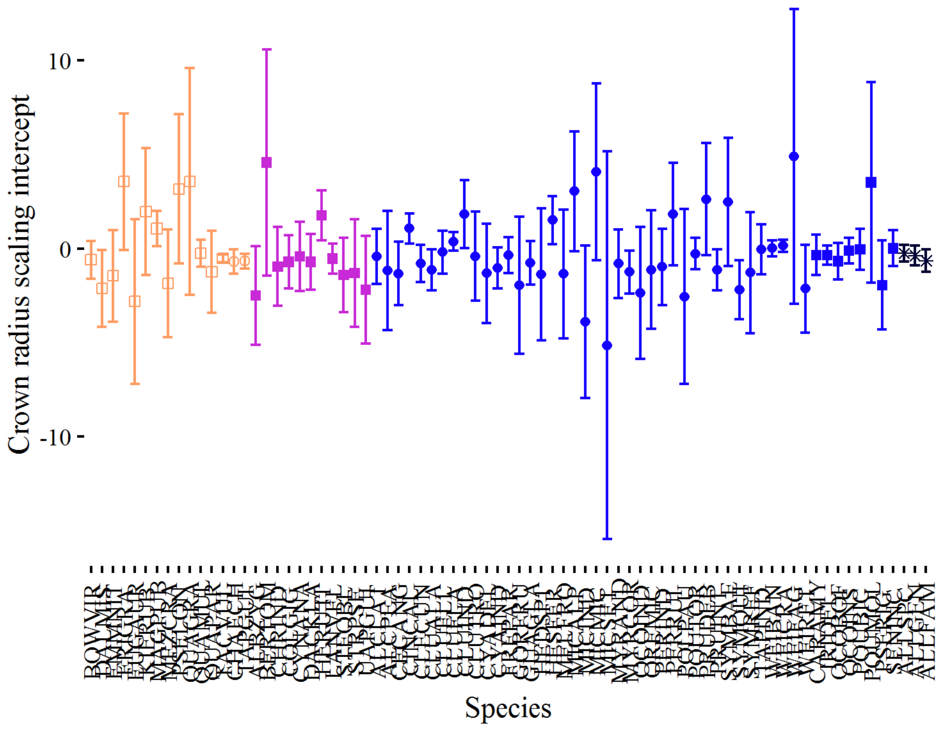


Figure S 6. Crown radius vs stem radius scaling intercepts per species via SMA regression. Each bar represents one species. ALLSPC coefficient filtered for species with 5 or more sampled individuals. TODO: incorporate CI's of means into composite CI.

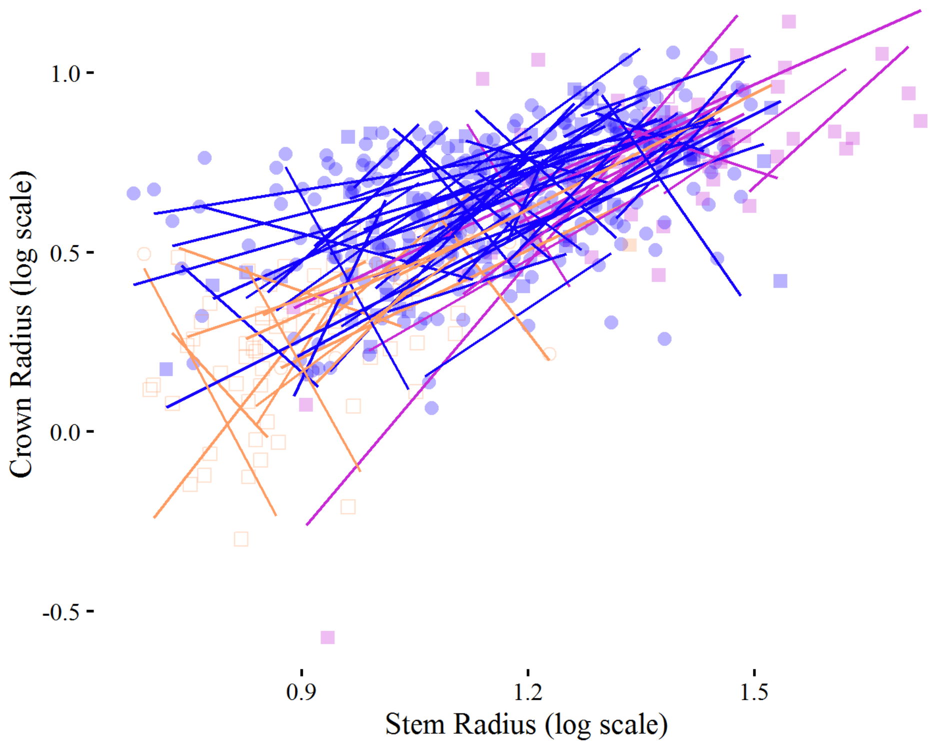


Figure S 7. Crown radius SMA scaling fits versus data. Each line represents one species, and each point represents one individual tree. Data filtered for species with 5 or more sampled individuals.

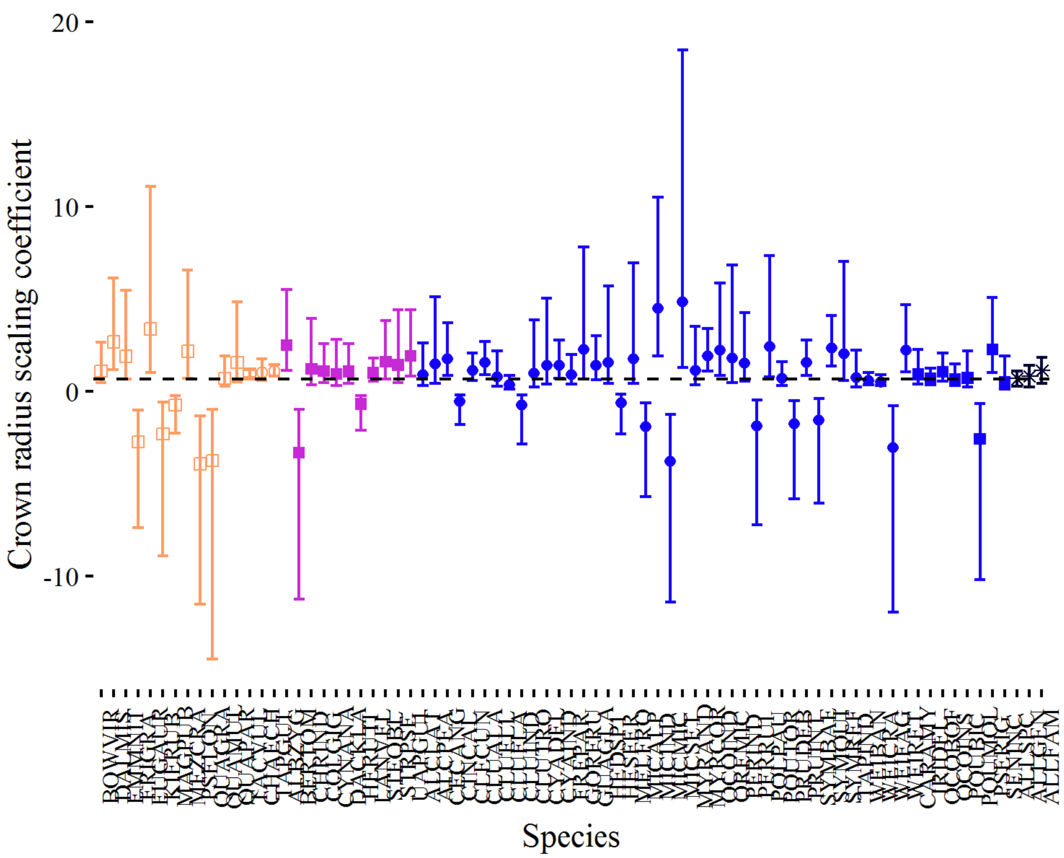


Figure S 8. Crown radius scaling coefficients per species via SMA regression. Each bar represents one species. ALLSPC coefficient filtered for species with 5 or more sampled individuals. The composite "ALL" coefficients comprised of species means.

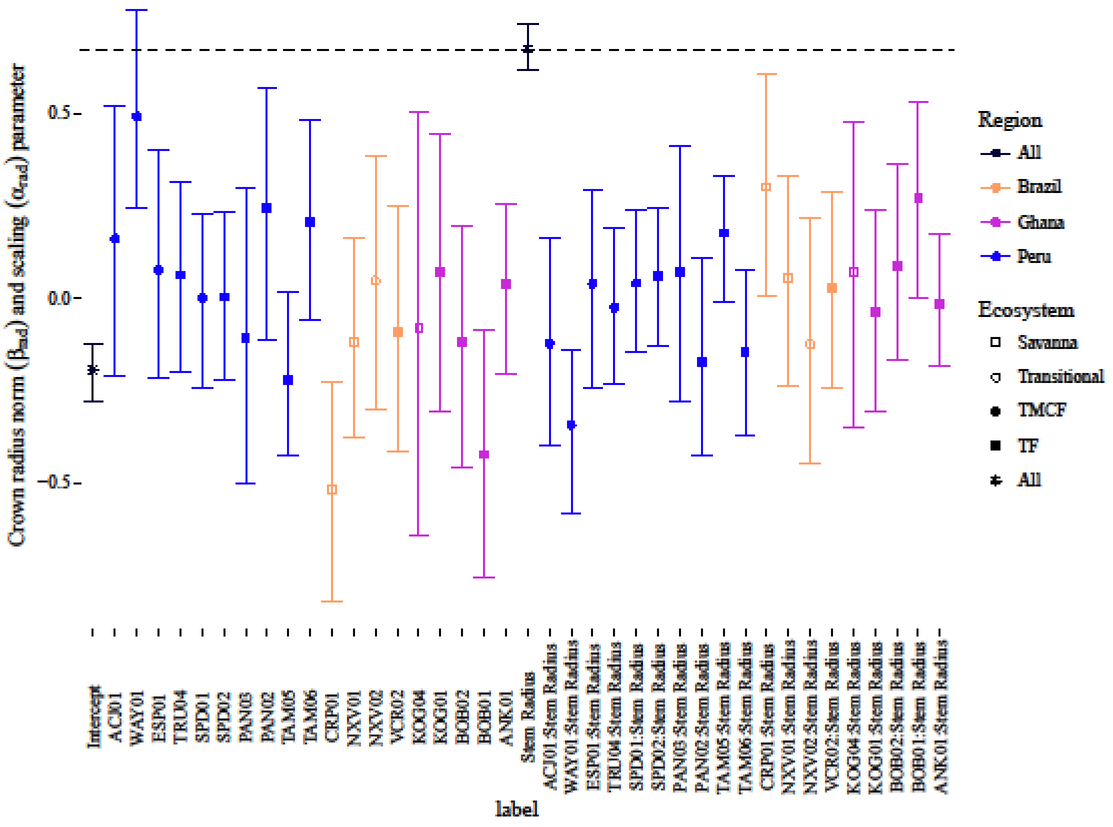


Figure S 9. Crown radius vs stem radius log-log LMM coefficients. As opposed to Figure 2., this figure includes the β terms. Terms with just site codes correspond to deviance-coded β (intercept) terms, and those with the ":Stem Radius" prefix correspond to deviance-coded α (slope) terms. Means have not been added to the site terms here. Hence, the overall mean β for ACJ01, for example, is the "ACJ01" mean plus the "Intercept" mean. The overall mean α for ACJ01 is the "ACJ01:Stem Radius" mean plus the "Stem Radius" mean. The black dotted line indicates the MST prediction for the Stem Radius term.

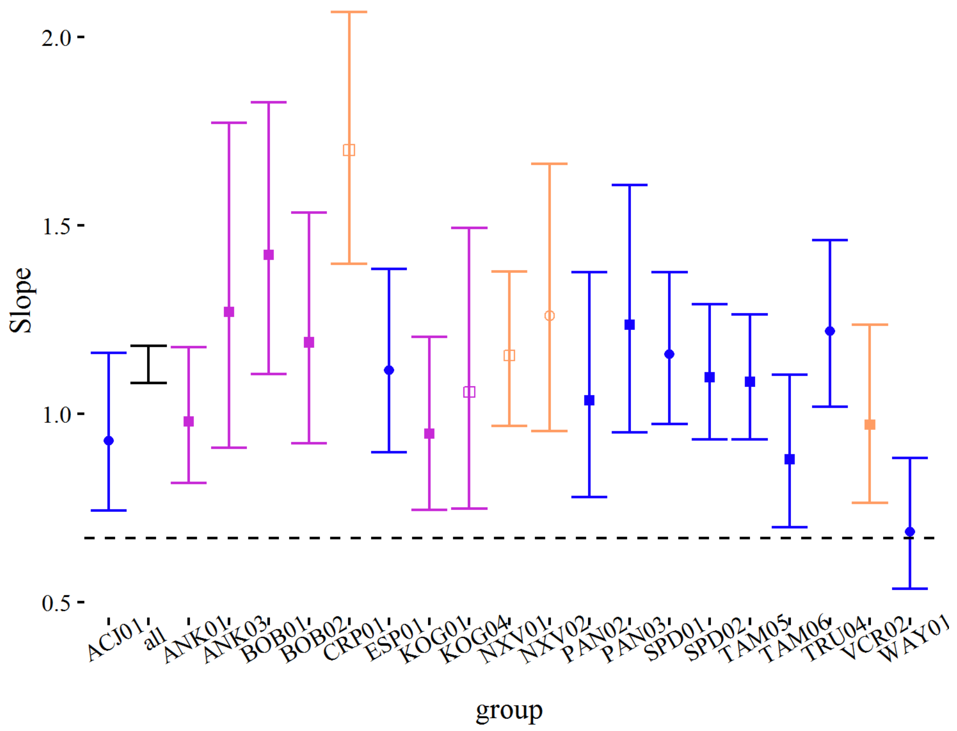


Figure S 10. Slope of log crown radius vs log stem radii per site according to SMA regression. Black bar is for fit with no grouping factors.

#### 5.3 MST CROWN SURFACE AREA SCALING

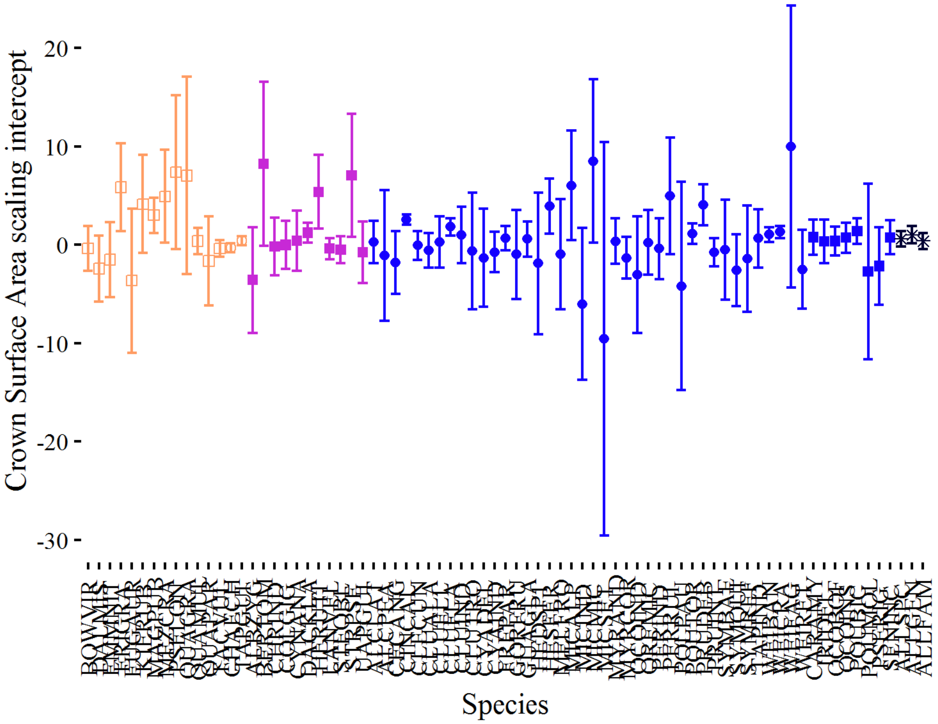


Figure S 11. Crown surface area vs stem radius scaling **intercepts** per species via SMA regression. Each bar represents one species. ALLSPC coefficient filtered for species with 5 or more sampled individuals.

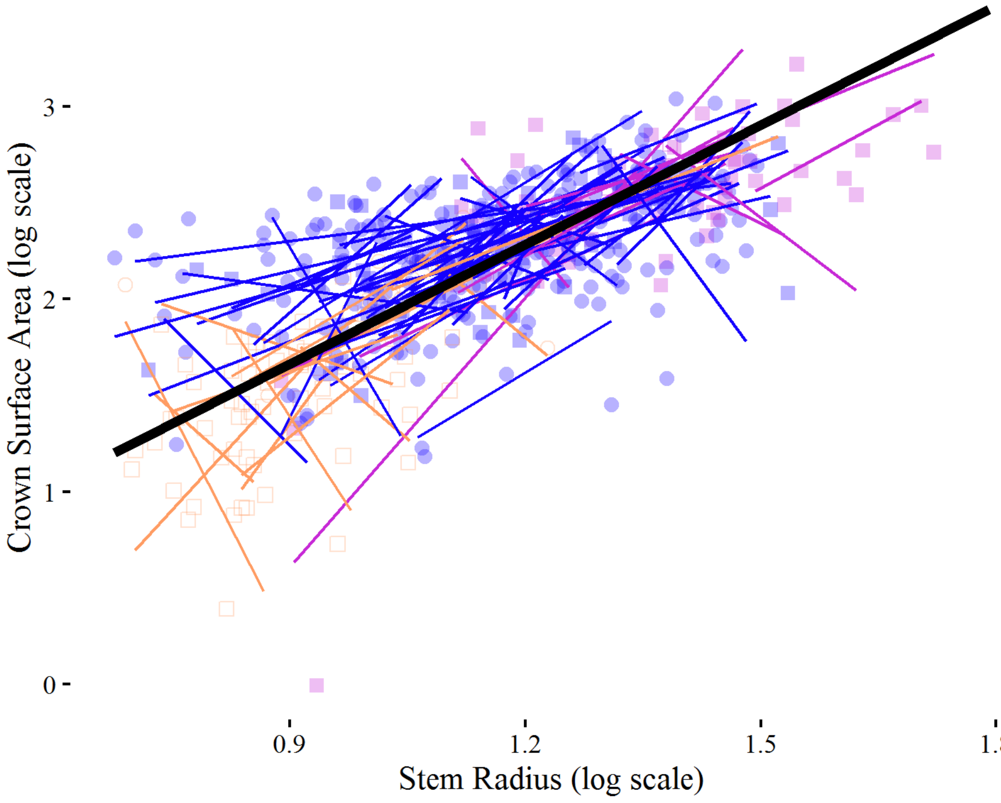


Figure S 12. Crown surface area vs stem radii per species. Colored lines are SMA fits per site, black line is fit ignoring site.

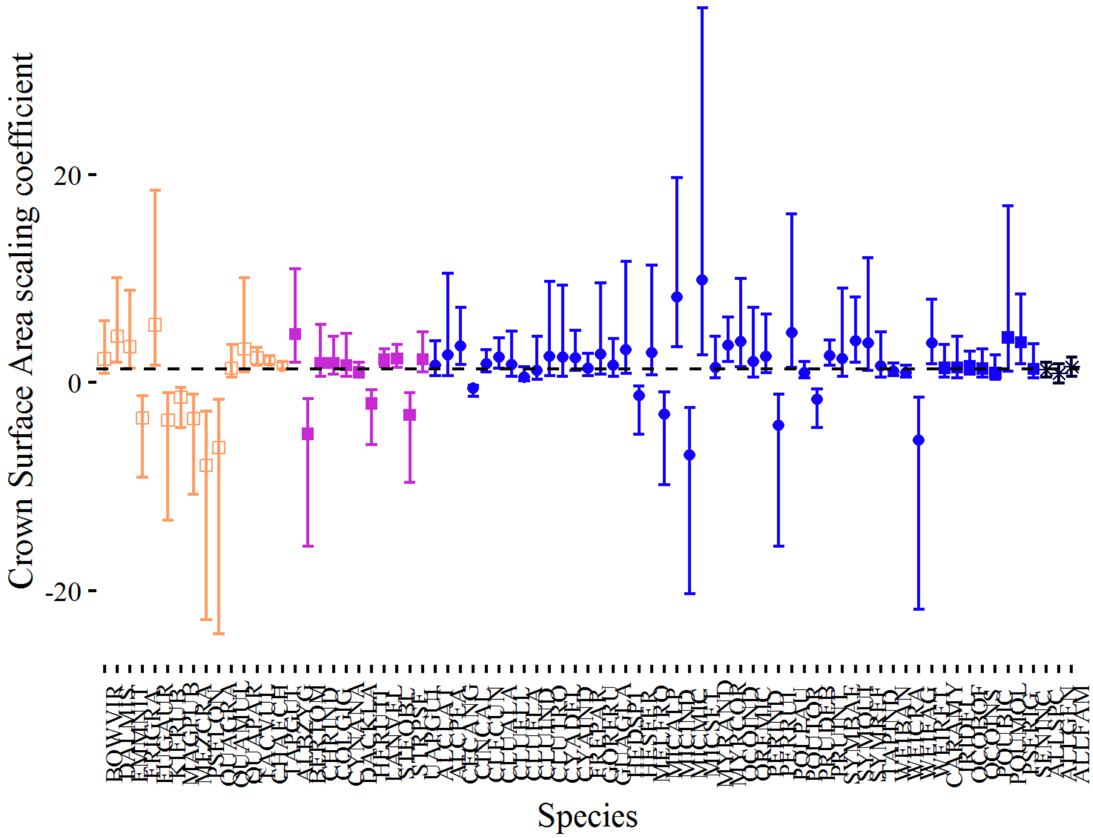


Figure S 13. Crown surface area vs stem radius scaling **coefficients** per species via SMA regression. Each bar represents one species. ALLSPC coefficient filtered for species with 5 or more sampled individuals. The composite "ALL" coefficients comprised of species means.

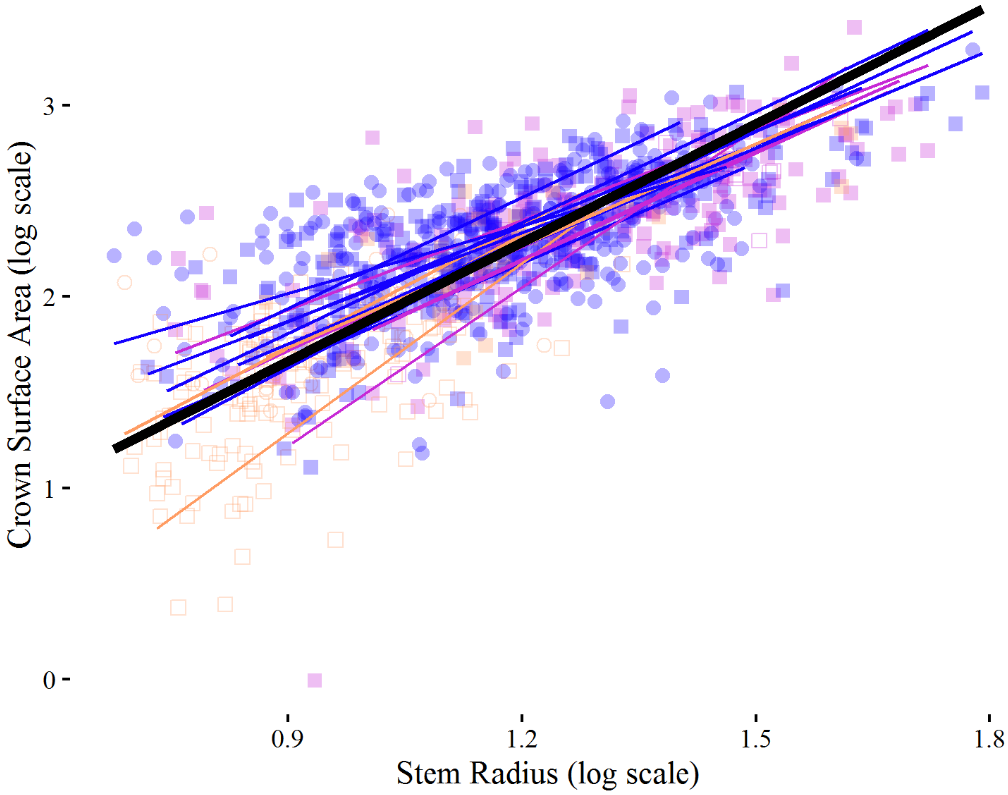


Figure S 14. Crown surface area vs stem radii per site. Colored lines are SMA fits per site, black line is fit ignoring site.

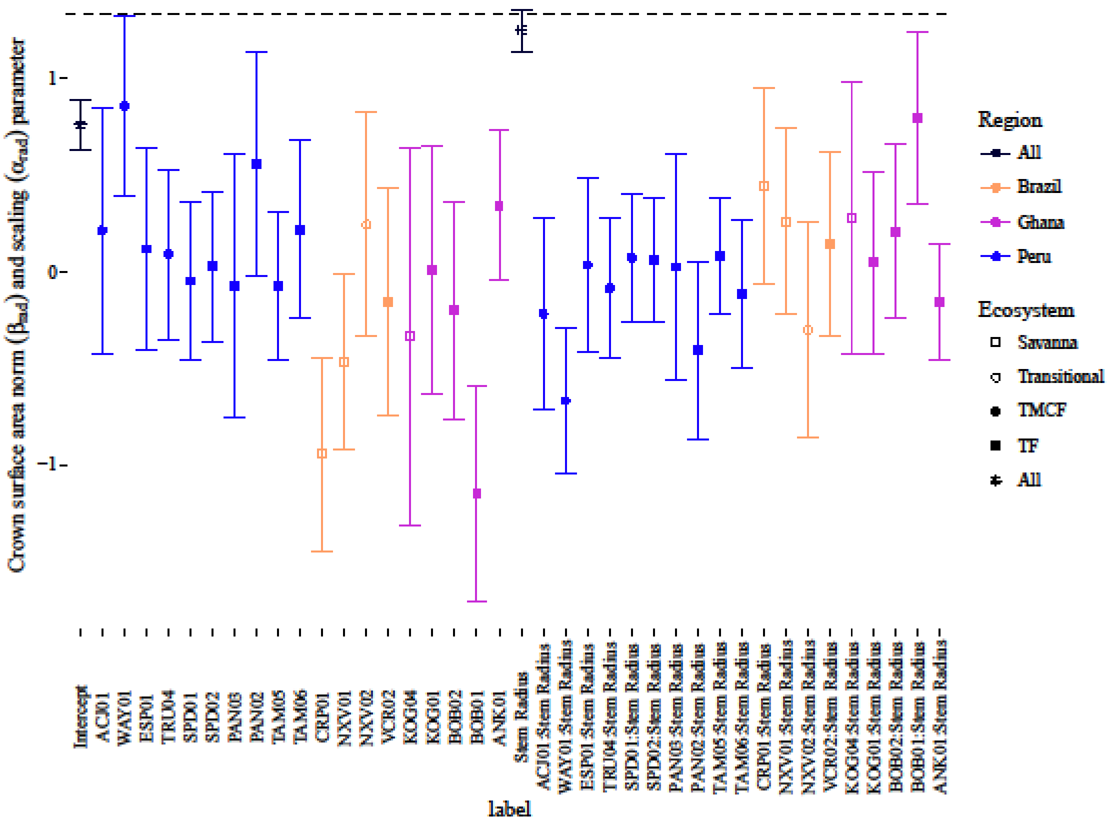


Figure S 15. Crown surface area vs stem radius log-log LMM coefficients. As opposed to Figure 2., this figure includes the β terms. Terms with just site codes correspond to deviance-coded β (intercept) terms, and those with the ":Stem Radius" prefix correspond to deviance-coded α (slope) terms. Means have not been added to the site terms here. Hence, the overall mean β for ACJ01, for example, is the "ACJ01" mean plus the "Intercept" mean. The overall mean α for ACJ01 is the "ACJ01:Stem Radius" mean plus the "Stem Radius" mean. The black dotted line indicates the MST prediction for the Stem Radius term.

#### 5.4 MST CROWN VOLUME SCALING

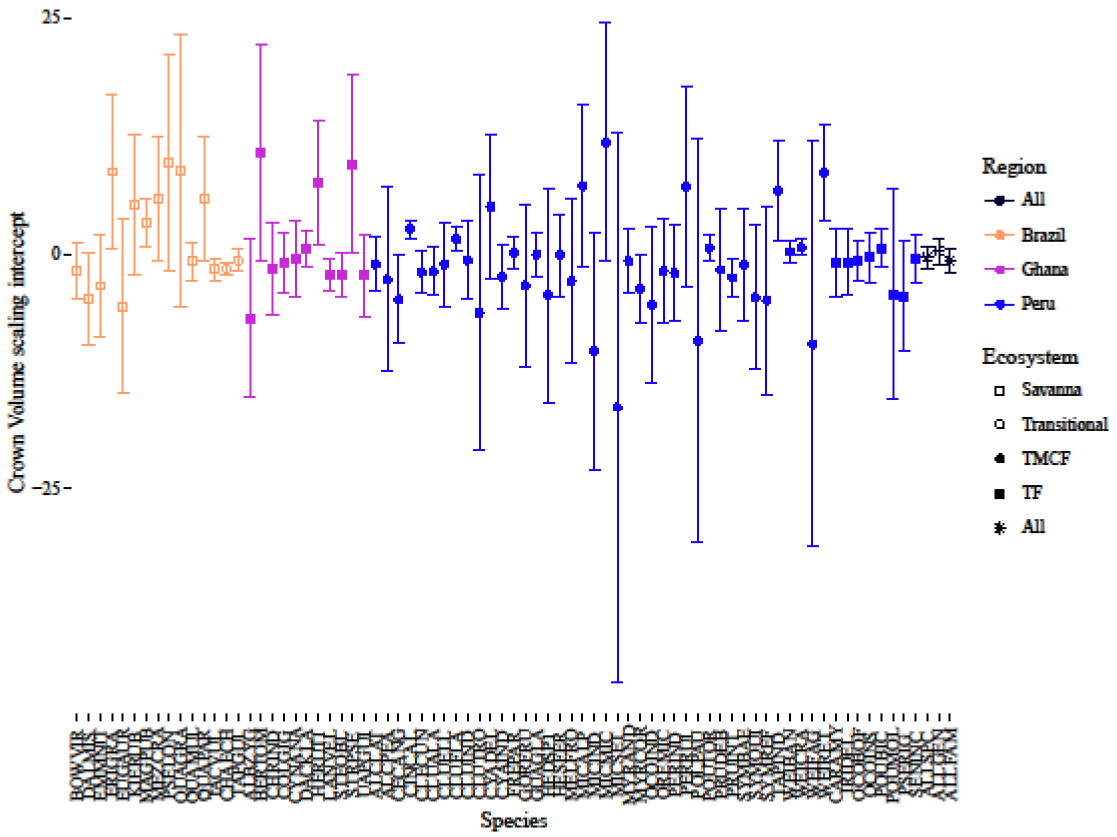


Figure S 16. Crown volume vs stem radius scaling **intercepts** per species via SMA regression. Each bar represents one species. ALLSPC coefficient filtered for species with 5 or more sampled individuals.

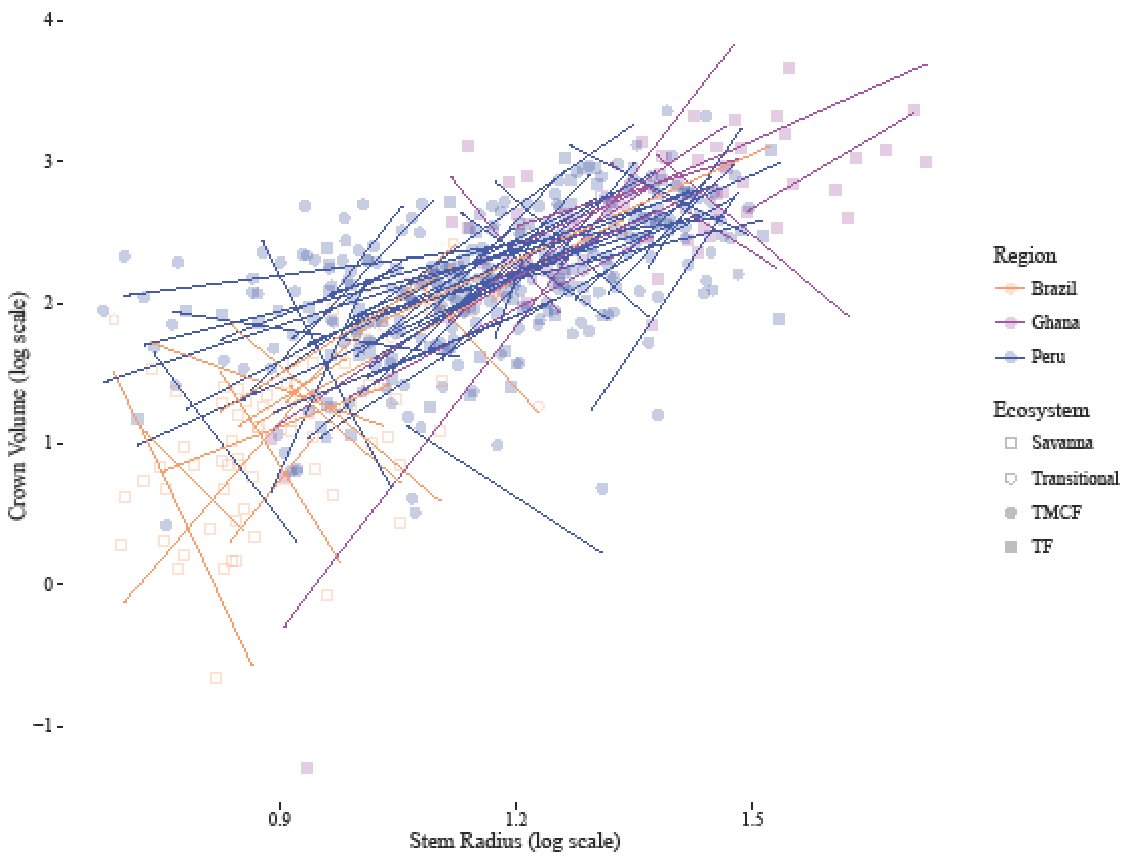


Figure S 17. Crown volume vs stem radii per species. Colored lines are SMA fits per site, black line is fit ignoring site.

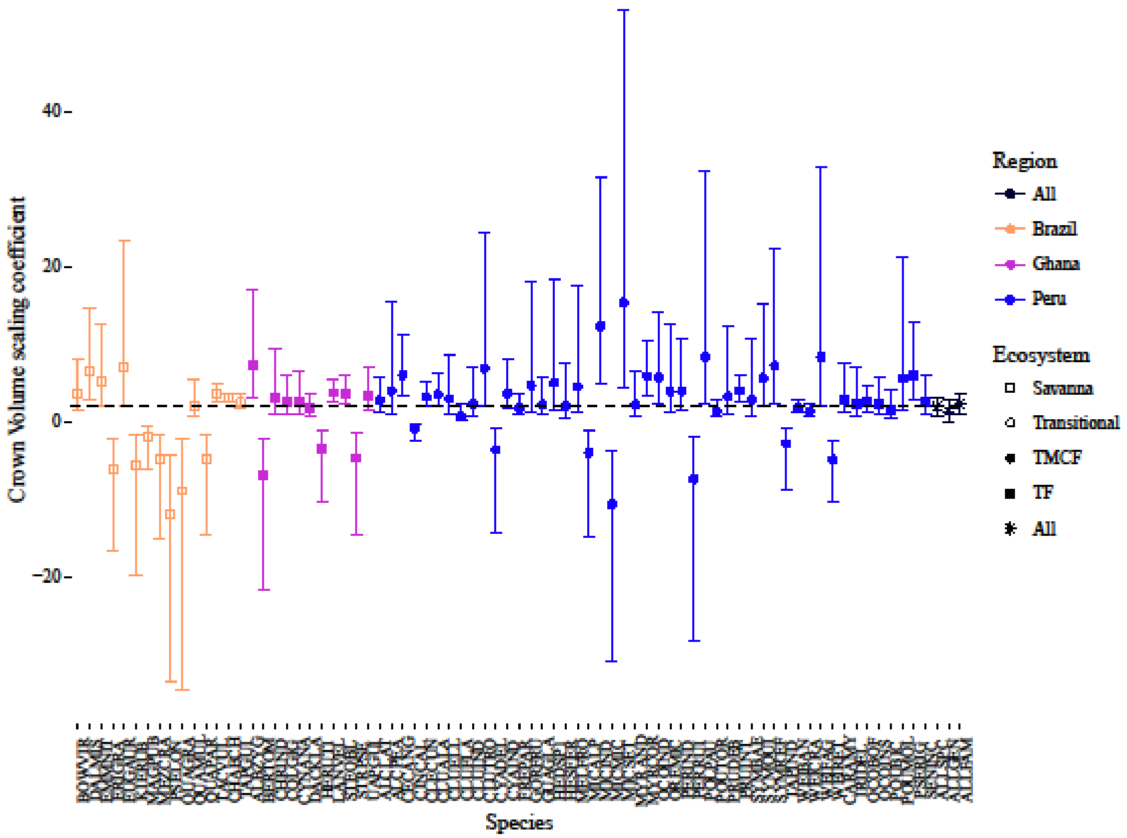


Figure S 18.  Crown volume vs stem radius scaling **coefficients** per species via SMA regression. Each bar represents one species. ALLSPC coefficient filtered for species with 5 or more sampled individuals. The composite "ALL" coefficients comprised of species means.

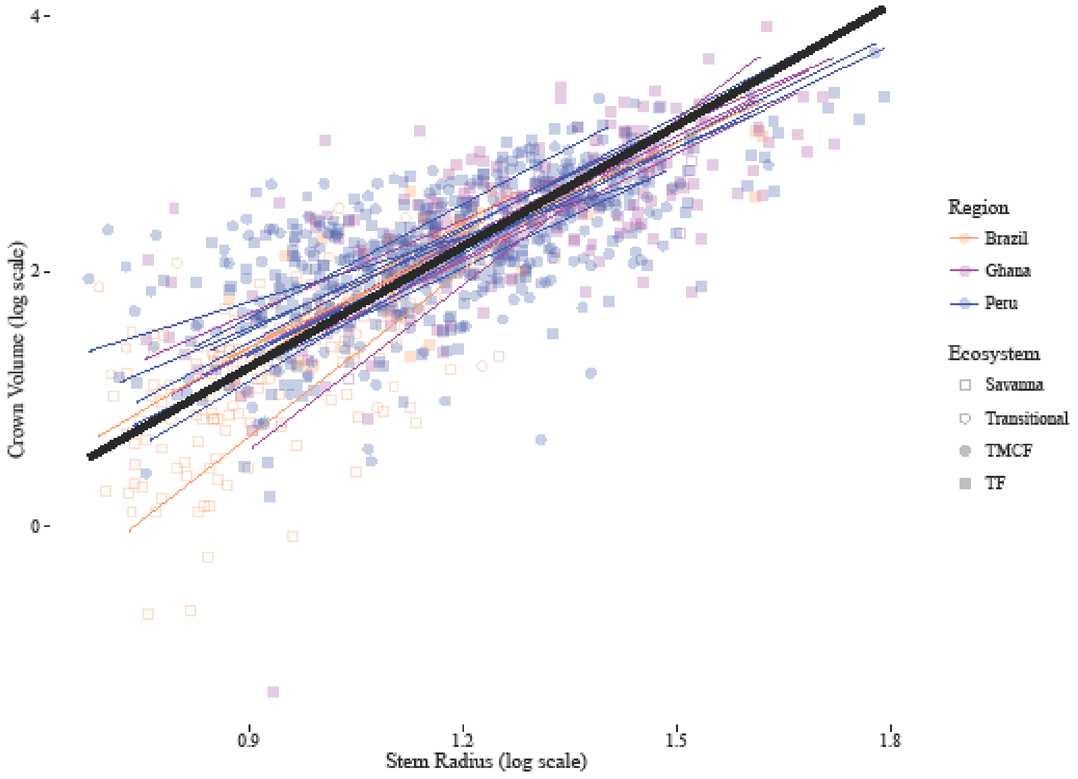


Figure S 19. Crown volume vs stem radii per site. Colored lines are SMA fits per site, black line is fit ignoring site.

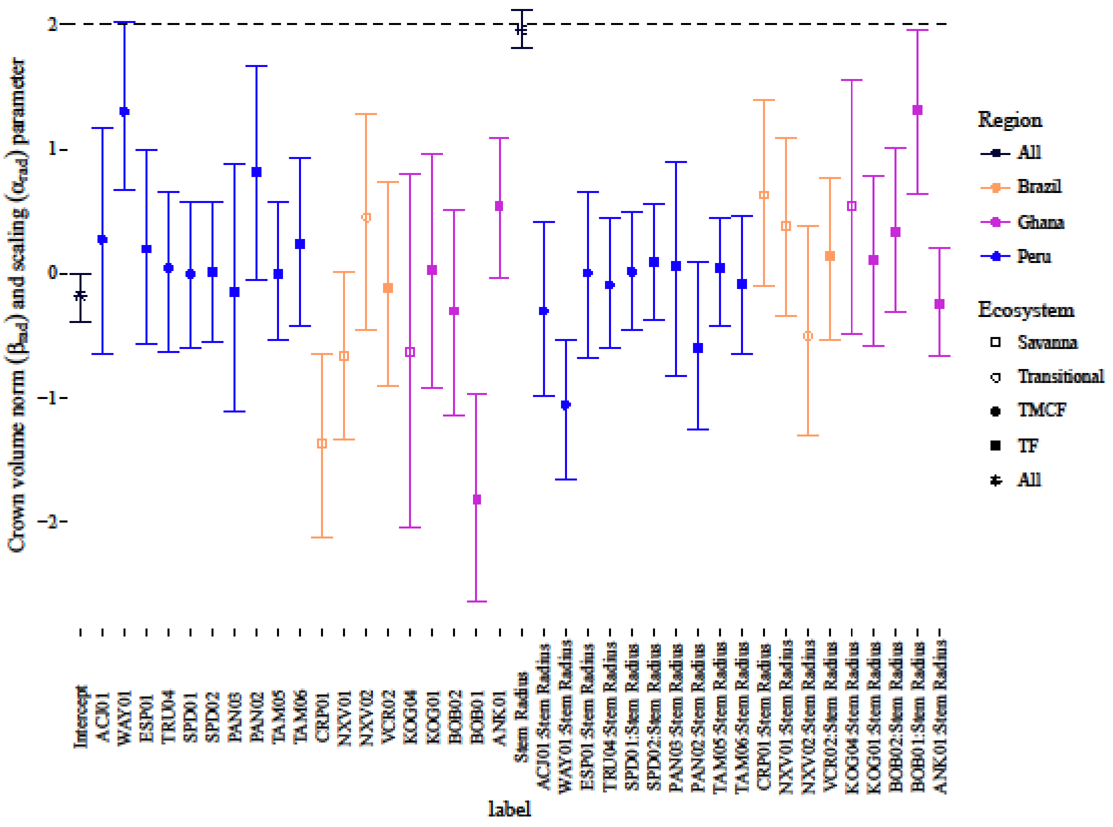


Figure S 20. Crown volume vs stem radius log-log LMM coefficients. As opposed to Figure 2., this figure includes the β terms. Terms with just site codes correspond to deviance-coded β (intercept) terms, and those with the ":Stem Radius" prefix correspond to deviance-coded α (slope) terms. Means have not been added to the site terms here. Hence, the overall mean β for ACJ01, for example, is the "ACJ01" mean plus the "Intercept" mean. The overall mean α for ACJ01 is the "ACJ01:Stem Radius"

#### 5.5 MST CROWN DEPTH SCALING

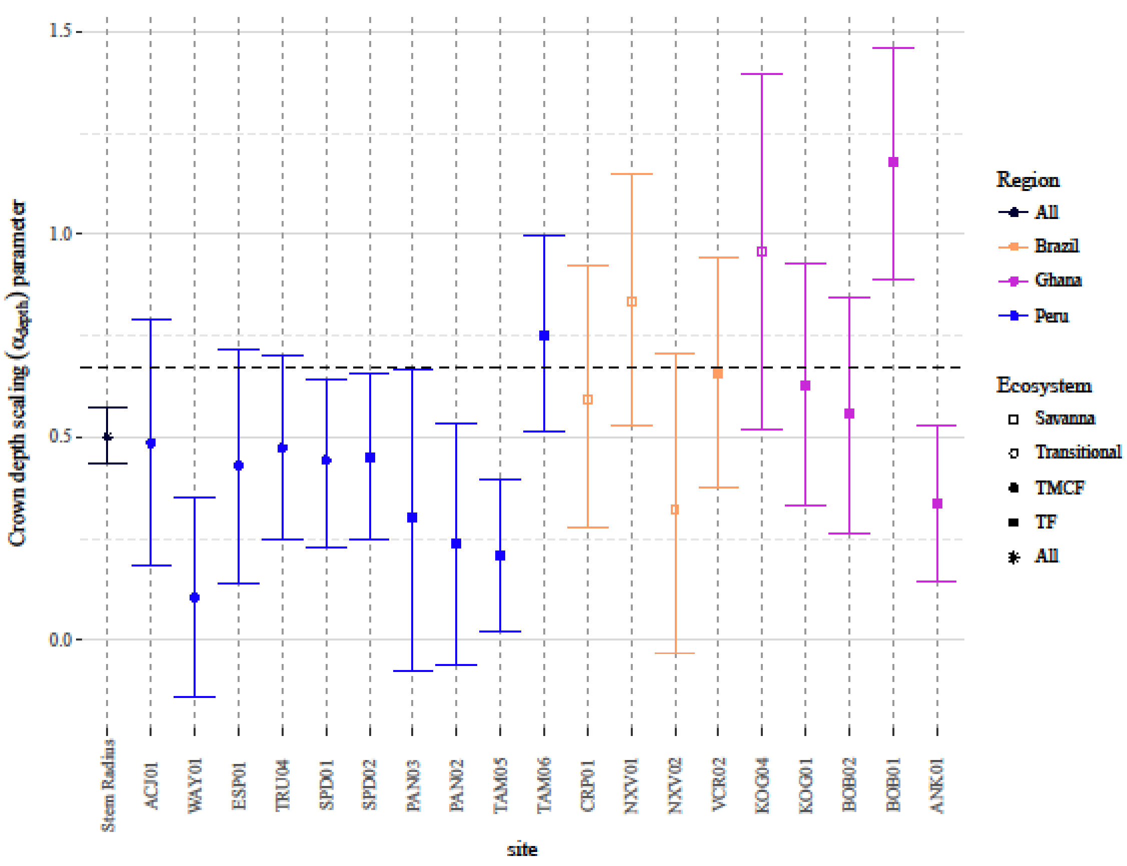


Figure S 21. Crown depth vs stem radius scaling coefficients as fit by LMM. Site coefficients are per-site slope effects with the mean effect added. Dotted line is the MST prediction for crown depth.

#### 5.6 CROWN DIMENSION MODEL RESIDUALS MAPPED ONTO PHYLOGENIES

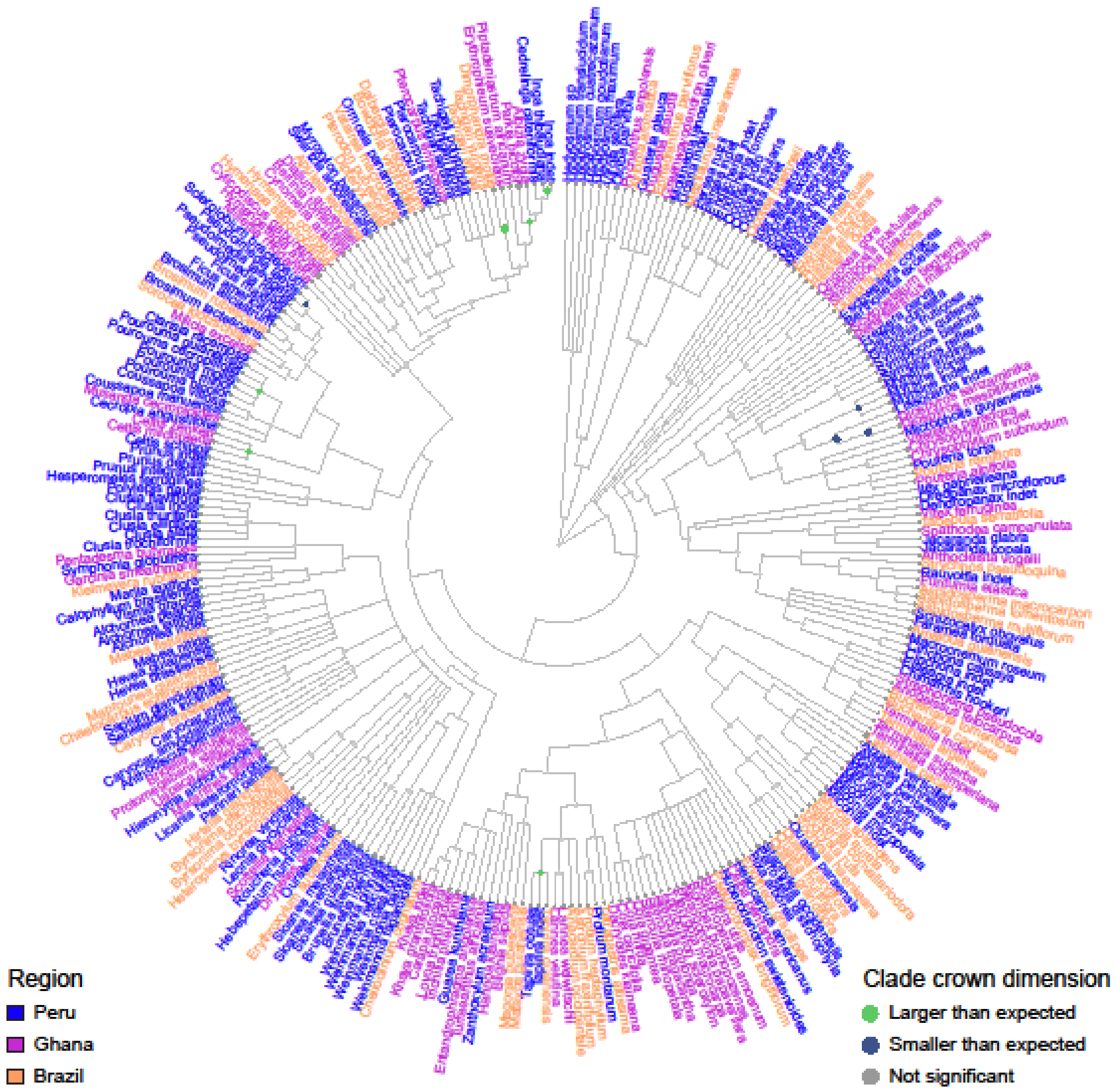


Figure S 22. Crown radius residuals mapped onto phylogeny. Residuals are per-species means from the crown radius vs stem radius scaling LMM with species as a random effect. A residual in this instance implies a difference in intercept, not slope, in the model. Size of circle corresponds to size of residual. Internal node states determined using fastANC (see Methods), with colors indicating direction and confidence of internal node estimates: 95% confidence interval of grey nodes intersects zero, yellow nodes indicate clades with crowns significantly.

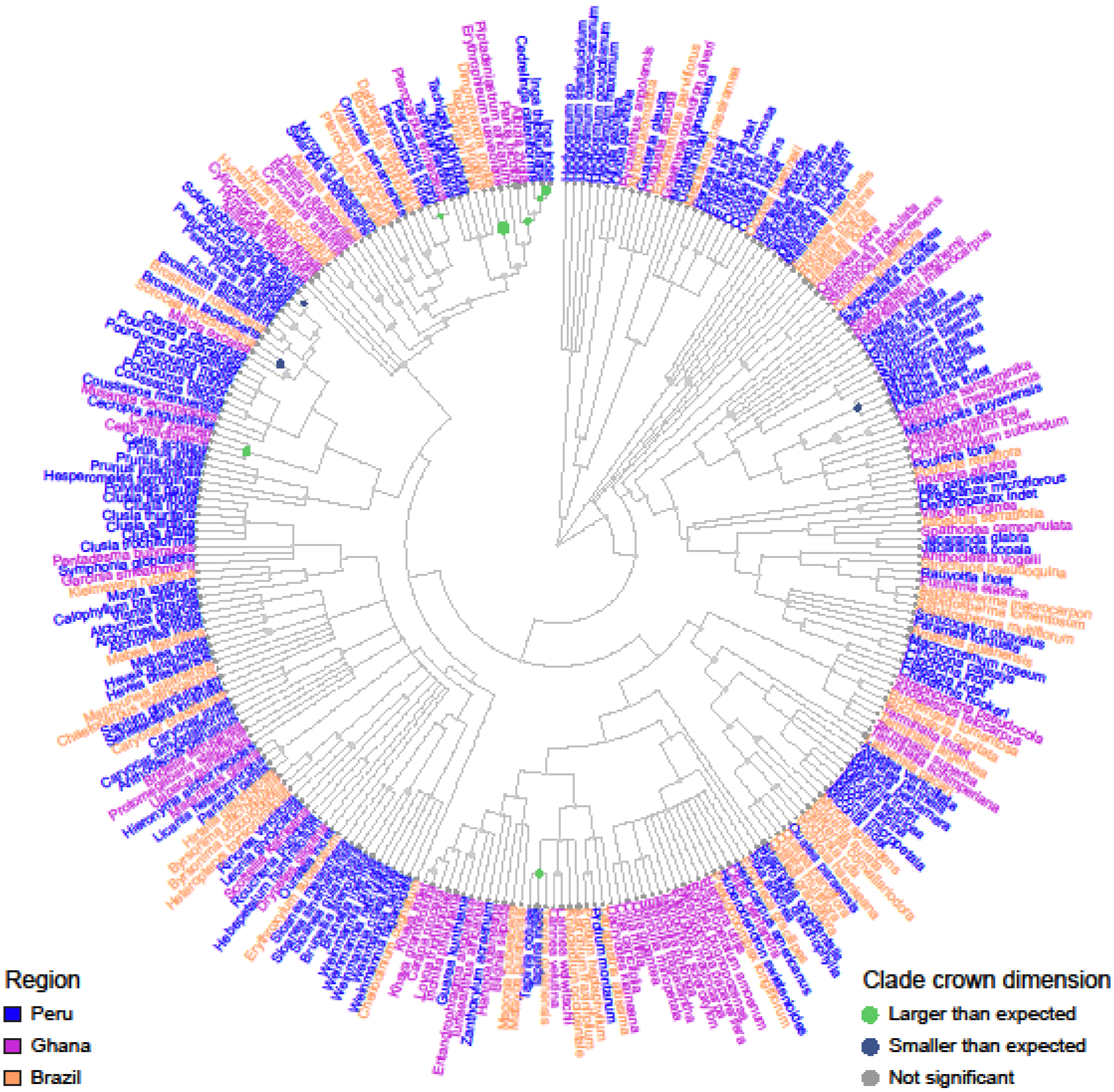


Figure S 23. Crown surface area residuals mapped onto phylogeny. Residuals are per-species means from the crown surface area vs stem radius scaling LMM with species as a random effect. A residual in this instance implies a difference in intercept, not slope, in the model. Size of circle corresponds to size of residual. Internal node states determined using fastANC (see Methods), with colors indicating direction and confidence of internal node estimates: 95% confidence interval of grey nodes intersects zero, yellow nodes indicate clades with crowns significantly.

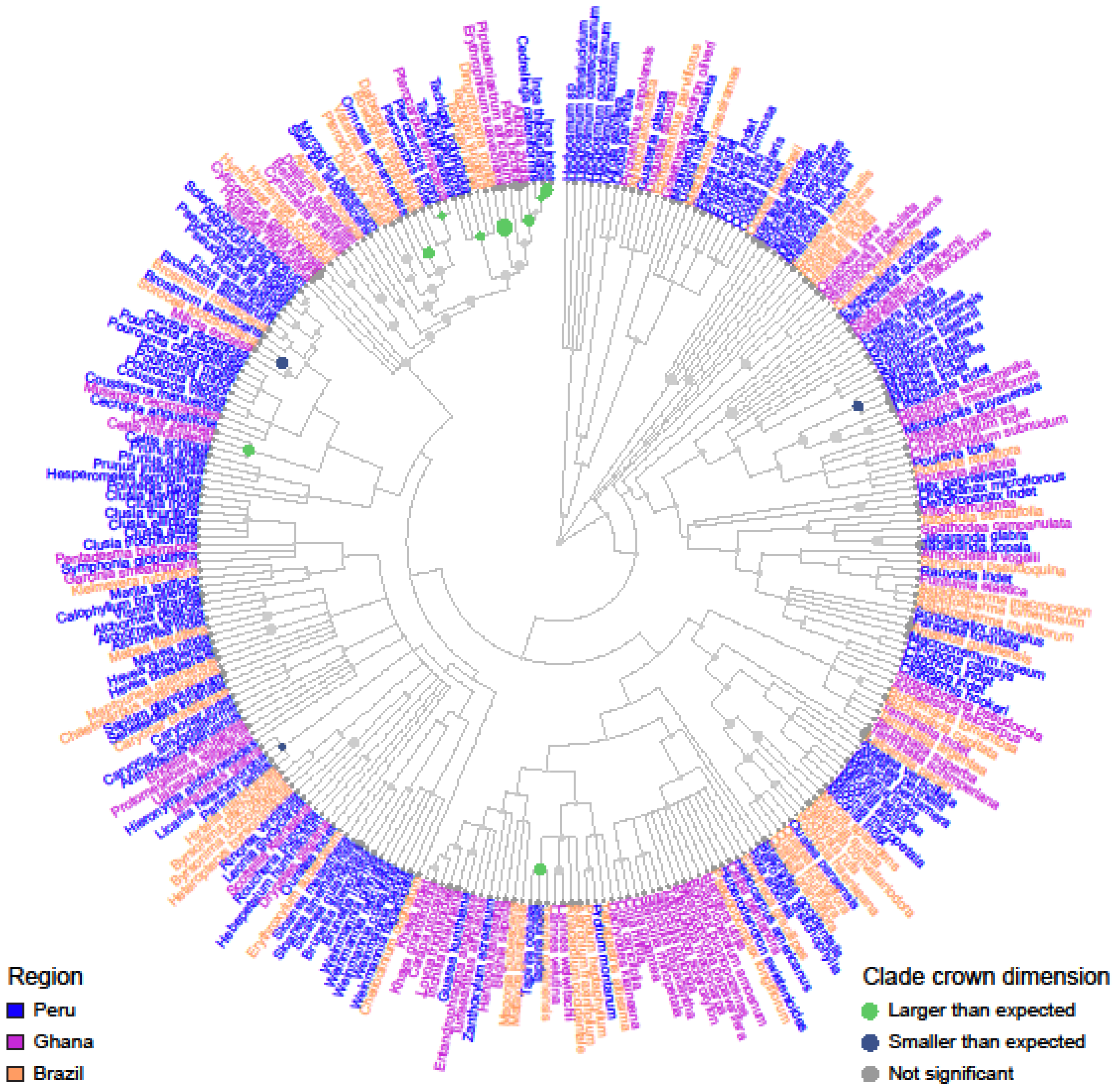


Figure S 24. Crown volume residuals mapped onto phylogeny. Residuals are per-species means from the crown surface area vs stem radius scaling LMM with species as a random effect. A residual in this instance implies a difference in intercept, not slope, in the model. Size of circle corresponds to size of residual. Internal node states determined using fastANC (see Methods), with colors indicating direction and confidence of internal node estimates: 95% confidence interval of grey nodes intersects zero, yellow nodes indicate clades with crowns significantly.

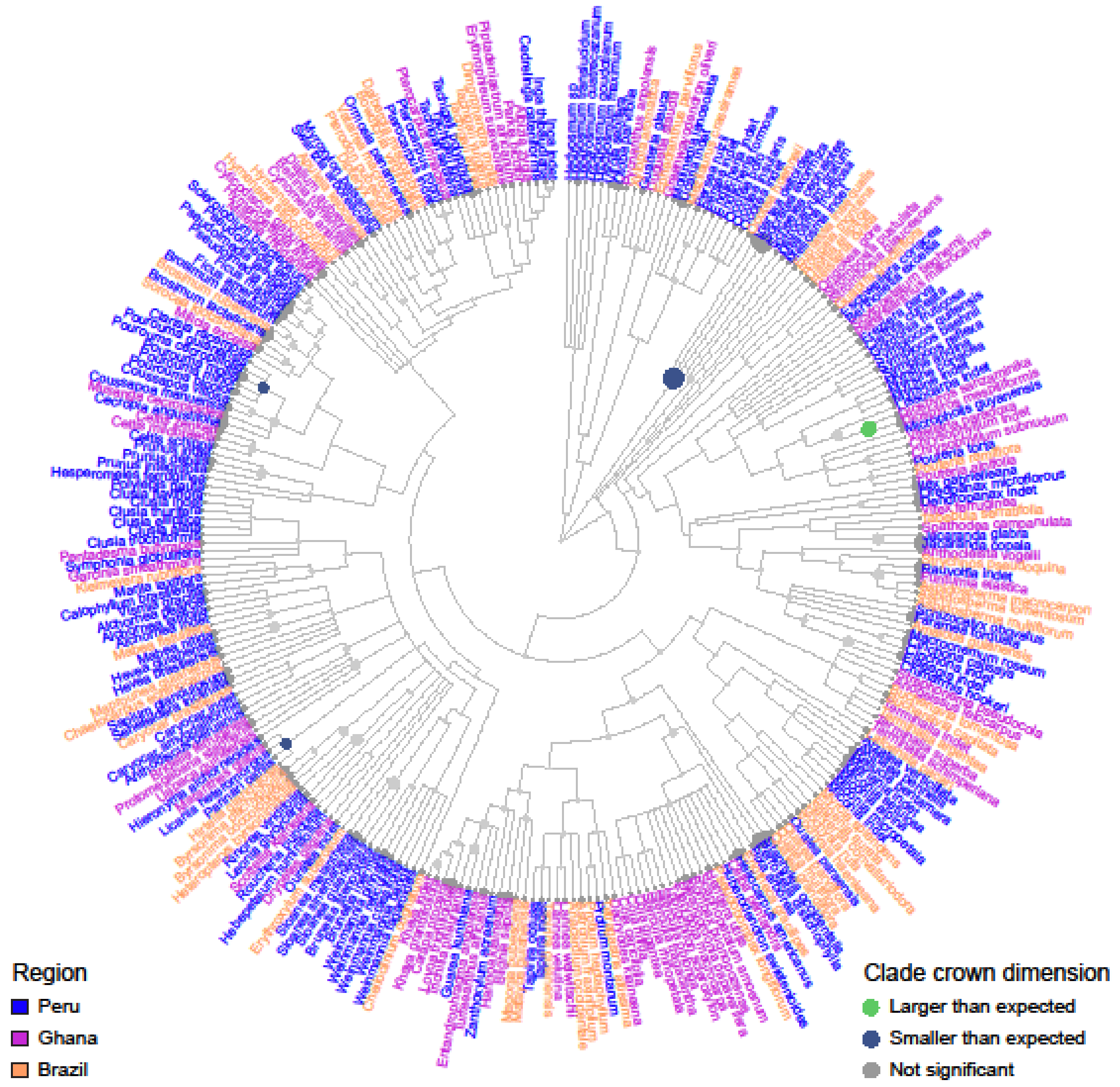


Figure S 25. Crown depth residuals mapped onto phylogeny. Residuals are per-species means from the crown depth vs stem radius + tree height scaling LMM with species as a random effect. A residual in this instance implies a difference in intercept, not slope, in the model. Size of circle corresponds to size of residual. Internal node states determined using fastANC (see Methods), with colors indicating direction and confidence of internal node estimates: 95% confidence interval of grey nodes intersects zero, yellow nodes indicate clades with crowns significantly.

#### 5.7 Crown relative depth models and predictions

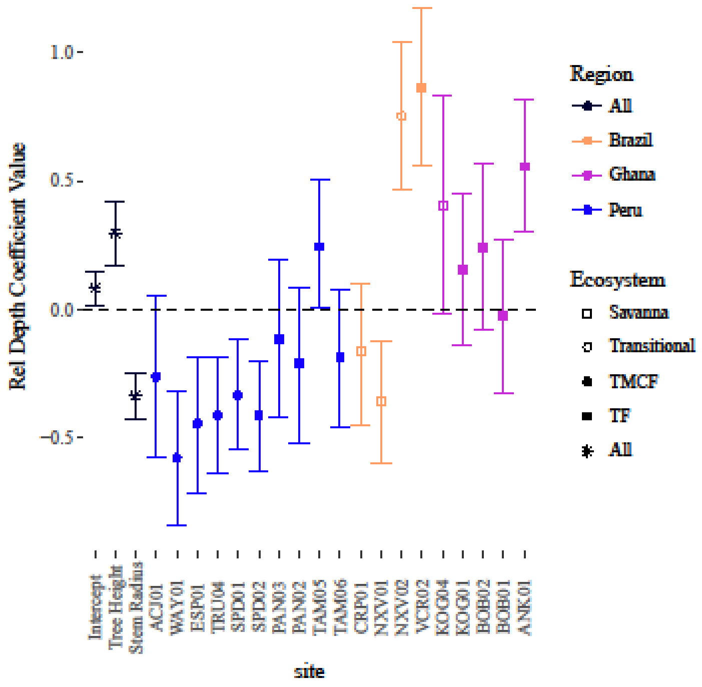


Figure S 26. Crown relative depth linear LMM predictors including tree height. As opposed to other similar figures here, the site effects are direct effects, and do not include any other added effects. These coefficients are derived from the following model: reldepth ~ (1|Species) + tree_height + stem_radius + site (Table S 6). All variables were scaled to have standard deviations of 1, and centered around 0. Sites are coded as deviance contrasts.

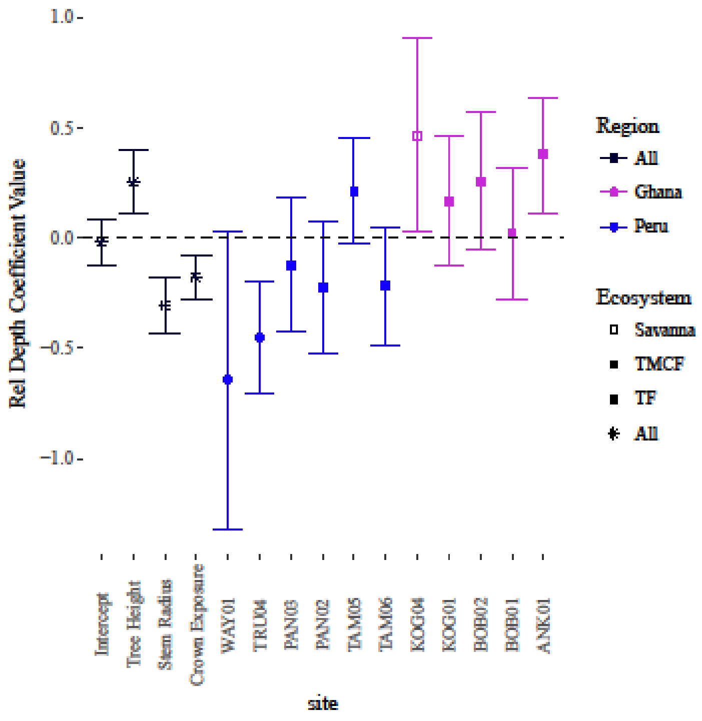


Figure S 27. Crown relative depth linear LMM predictors including tree height and crown exposure. As opposed to other similar figures here, the site effects are direct effects, and do not include any other added effects. These coefficients are derived from the following model: reldepth ~ (1|Species) + tree_height + stem_radius + crown_exposure + site (Table S 6). All variables were scaled to have standard deviations of 1, and centered around 0. Sites are coded as deviance contrasts. Crown exposure was only measured in a subset of plots.

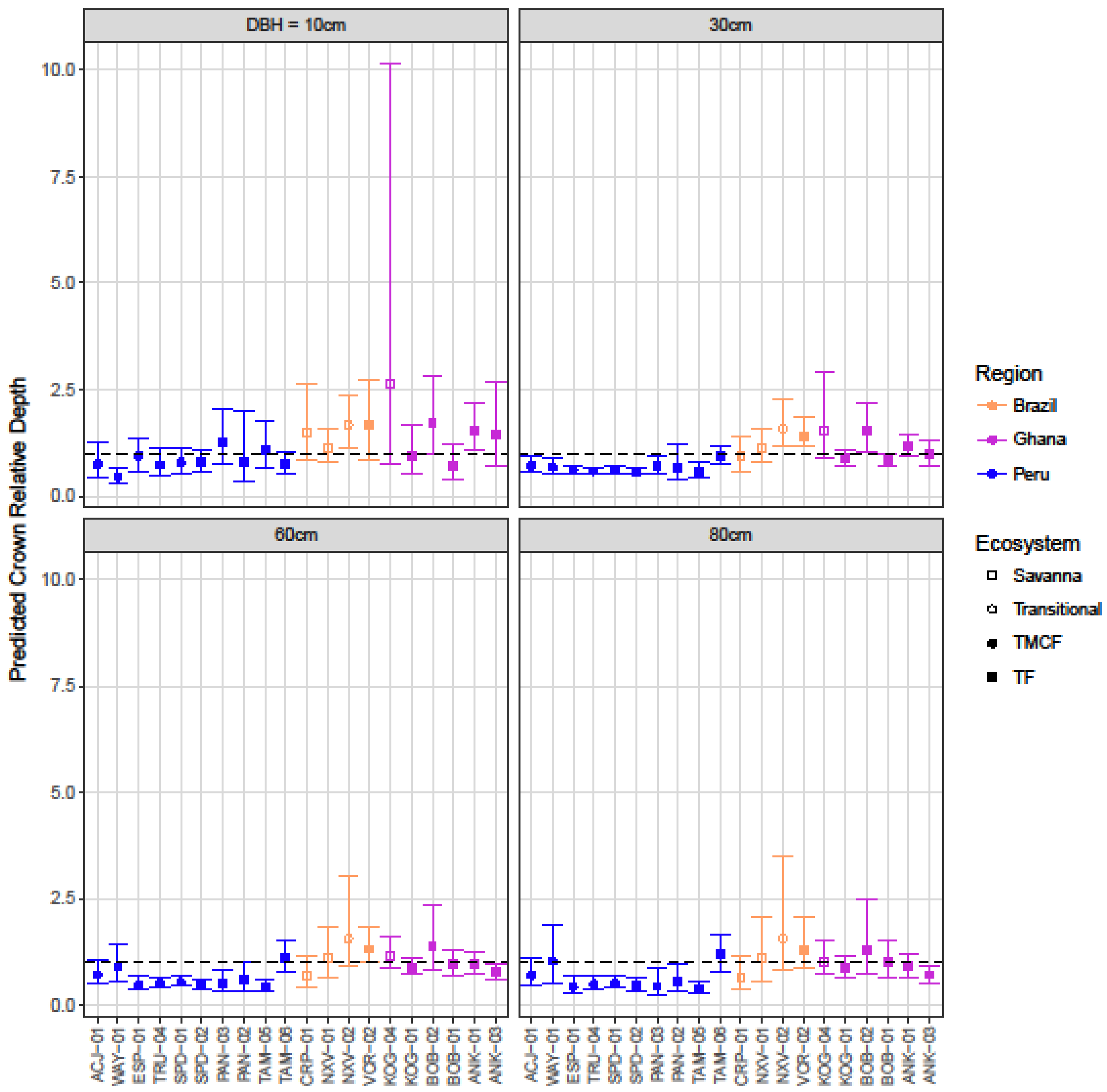


Figure S 28. Predicted relative crown depths and 95% CIs from log-log LMM for 4 tree sizes. Dashed line indicates a relative depth of one, when crown depth and width are equal. The model also included tree height, which was fixed using our height vs stem radius allometry (see SI) and averaged across sites so the same DBH and height were used for each site. CIs are estimated using a bootstrap algorithm with 100 iterations. The LMM formula used was: log10(reldepth) ~ (1+ log10(tree_height)|Species) + (1+ log10(stem_radius)|Species) + log10(tree_height)*site + log10(stem_radius)*site. Sites were coded with deviance contrasts.

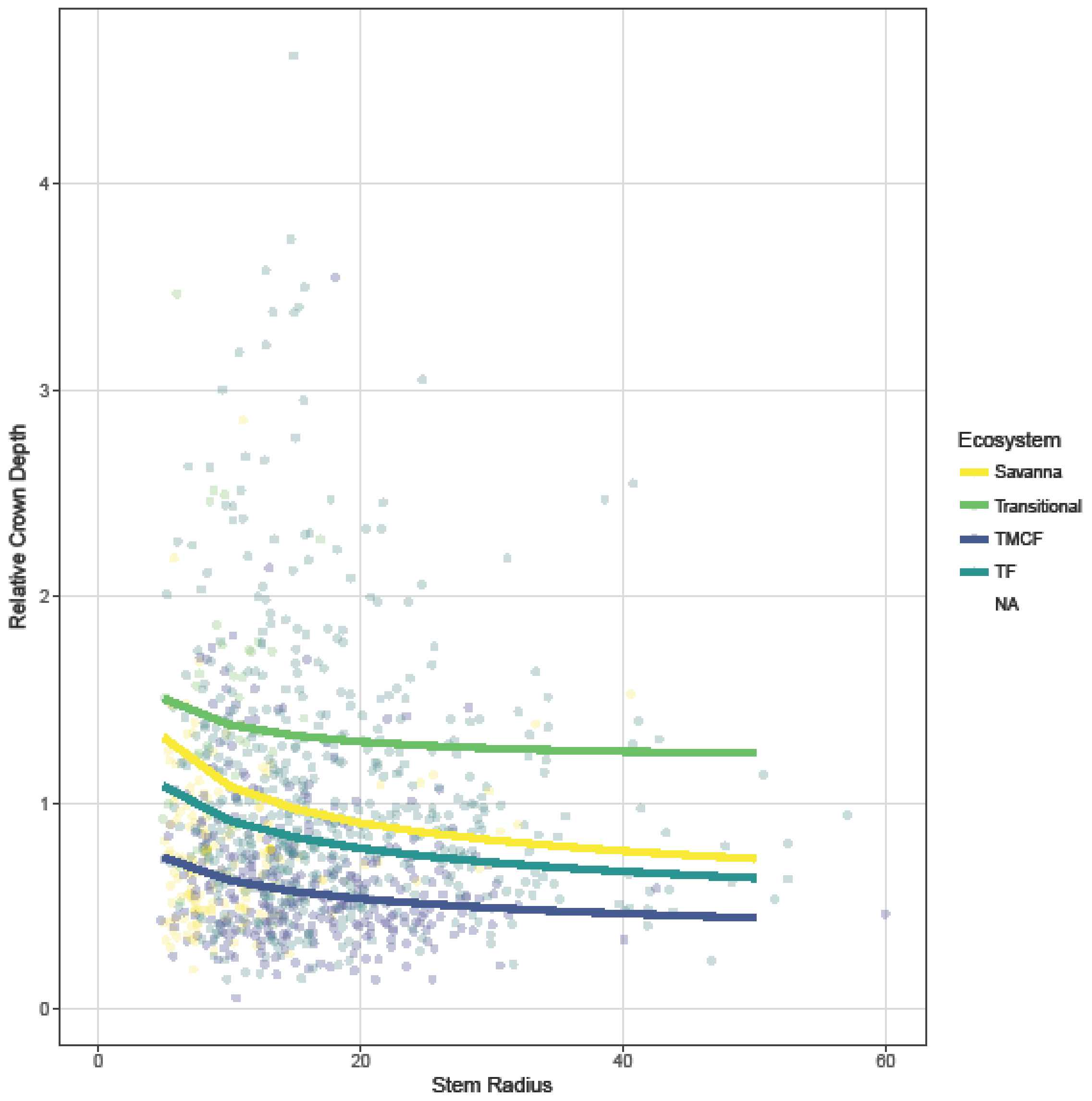


Figure S 29. Predicted crown relative depths across ecosystems based on model described in Table S 8. .

#### 5.8 Observed crown dimensions

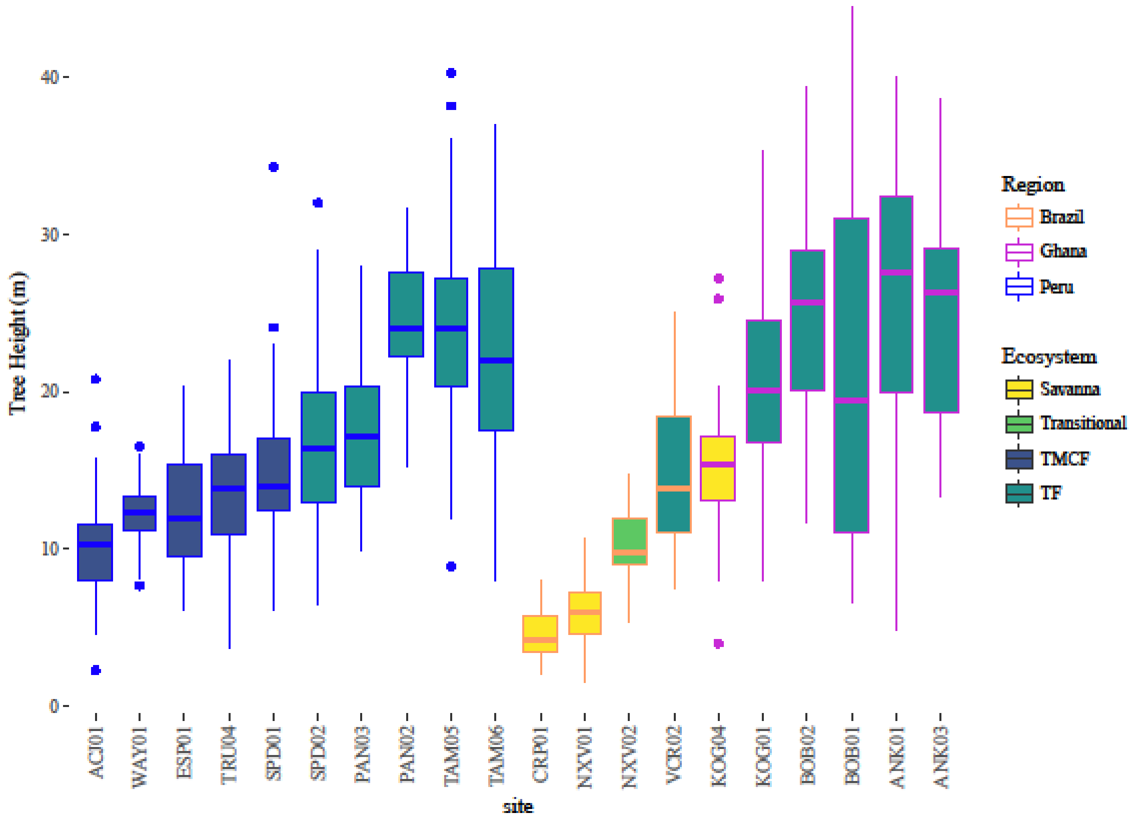


Figure S 30. Observed heights of sampled trees.

#### 5.9 PREDICTED CROWN DIMENSIONS

(a) 10cm DBH

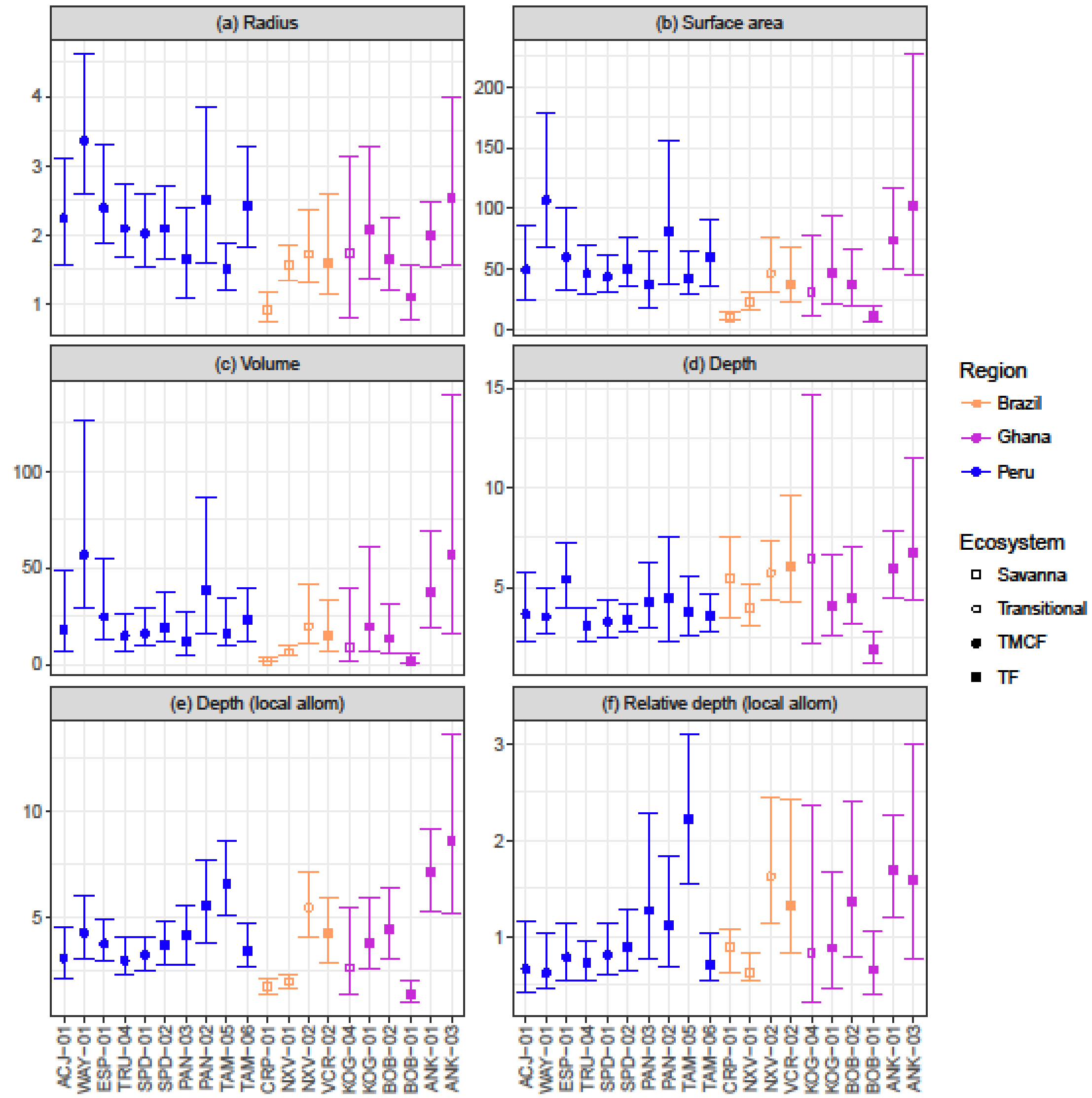

(b) 60cm DBH

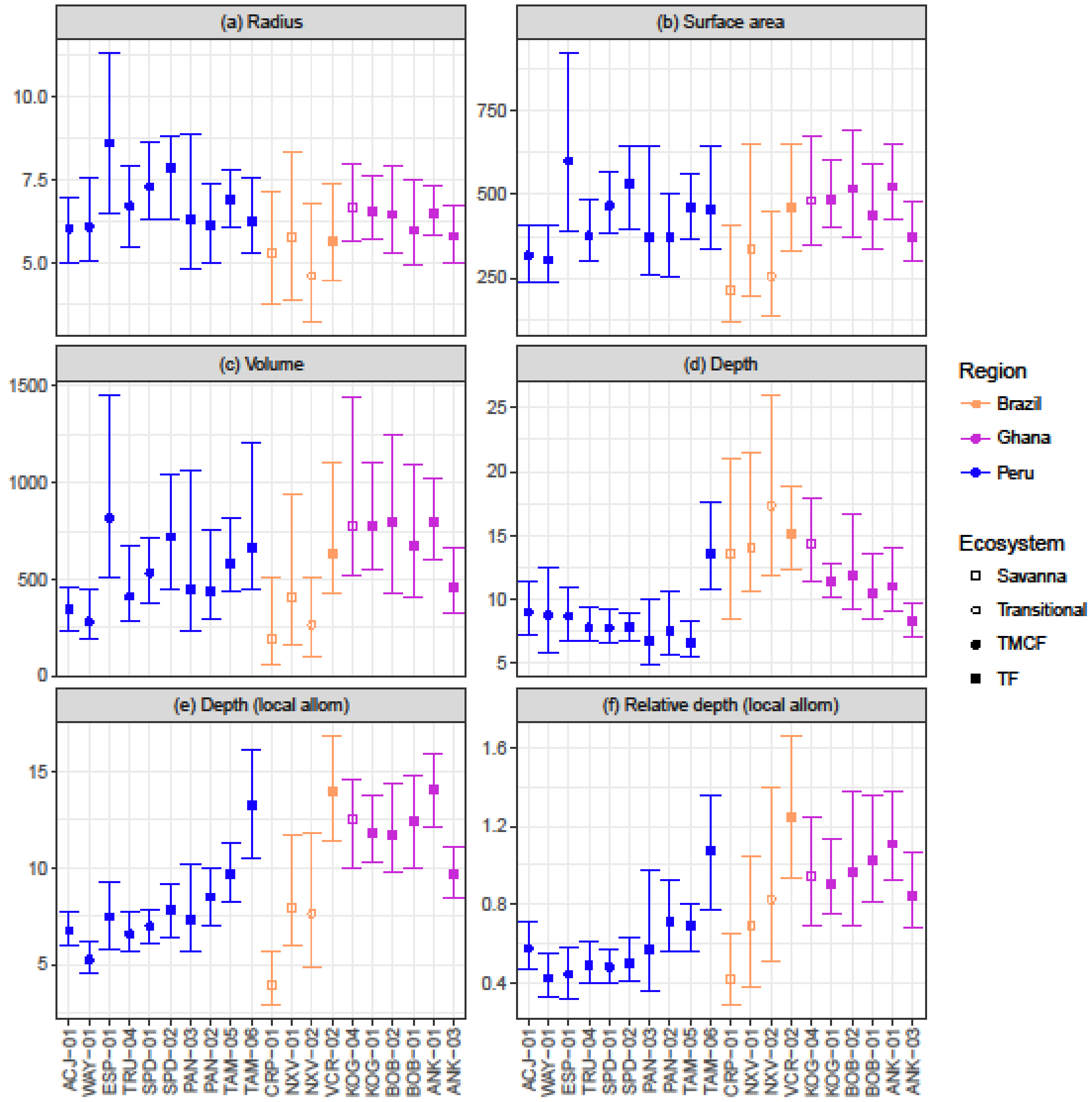
(a) Radius
(b) Surface area
(c) Volume
(d) Depth
(e) Depth (local allom)
(f) Relative depth (local allom)
Region
Brazil
Ghana
Peru
Ecosystem
Savanna
Transitional
TMCF
TF
ACJ−01
WAY−01
ESP−01
TRU−04
SPD−01
SPD−02
PAN−03
PAN−02
TAM−05
TAM−06
CRP−01
NXV−01
NXV−02
VCR−02
KOG−04
KOG−01
BOB−02
BOB−01
ANK−01
ANK−03

(c) 80cm DBH

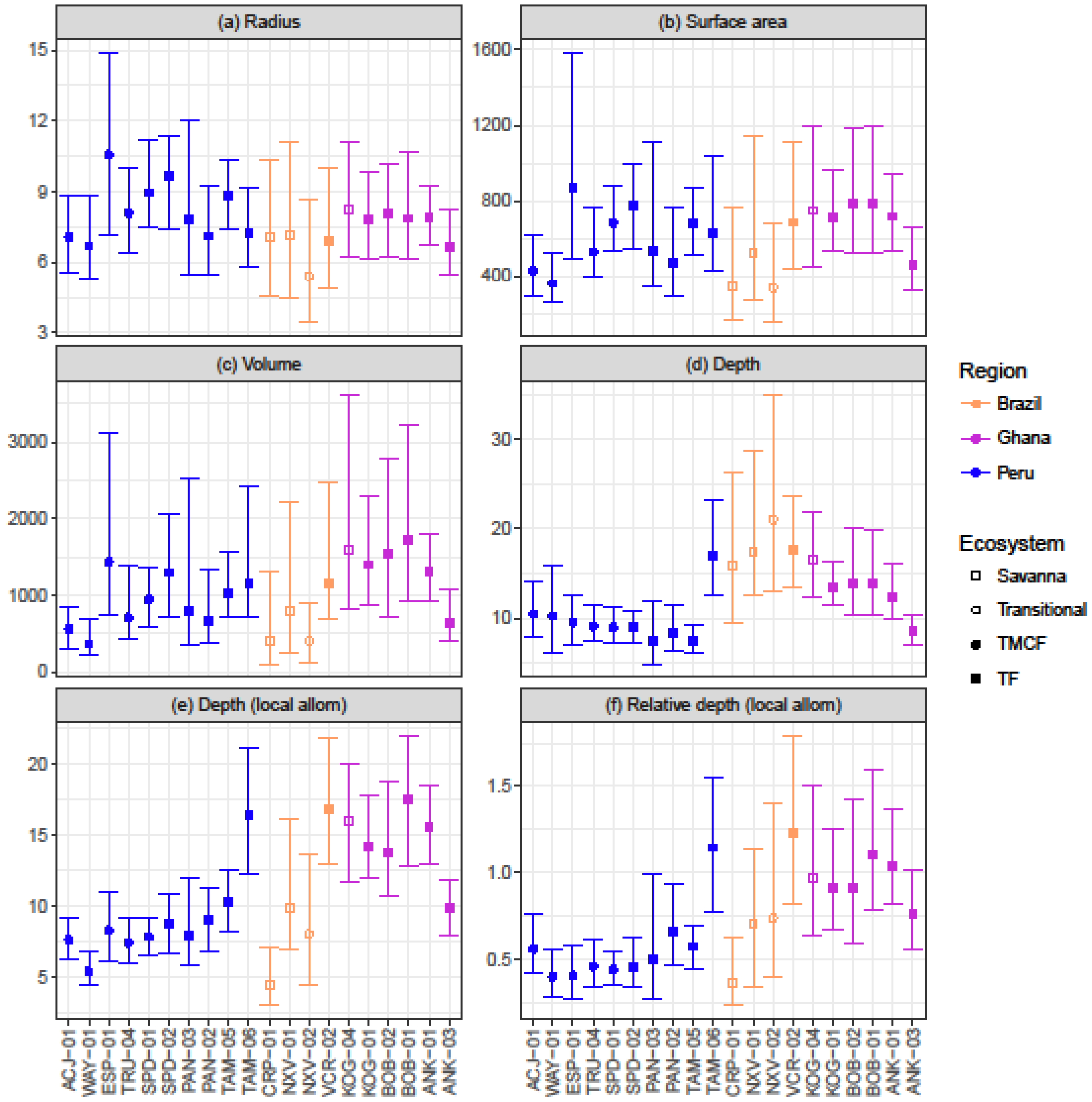


Figure S 31. Predicted crown dimensions and 95% CIs for 10cm (a), 60cm (b), and 80cm (c) DBH trees. Crown depth based on universal predicted height for 10cm, 60cm (21.4m) and 80cm (24.8m) trees. "Local allom" indicates that the tree height term in the model is generated from a height-DBH allometry fit from that particular site's data. Predictions are derived from the LMM models described in the main text, and CIs were estimated using a bootstrap algorithm with 100 iterations. Units are m, $m^2$, and $m^3$.

#### 5.10 Crown depth predictions

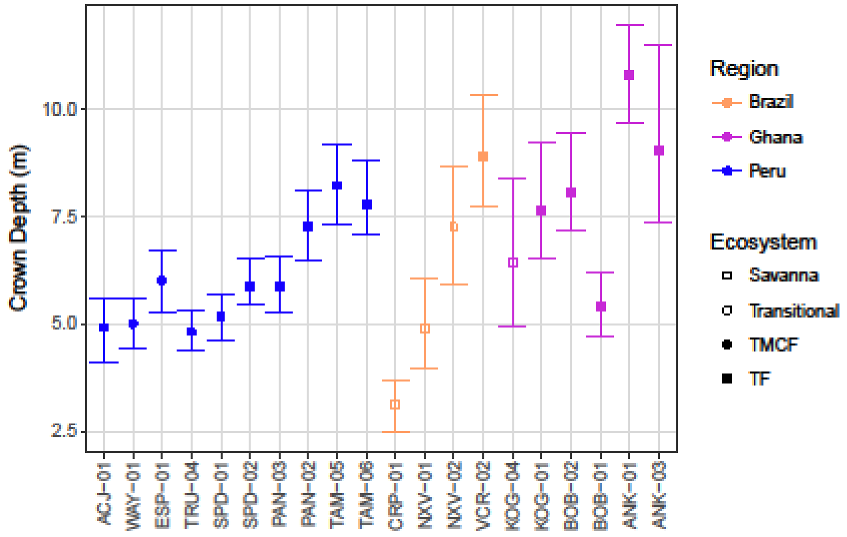


Figure S 32. Predicted crown depth and 95% CIs for a 30cm DBH tree, not including height as a predictor. Predictions are derived from the LMM models described above, and CIs estimated using a bootstrap algorithm with 100 iterations.

#### 5.11 Predicted tree height

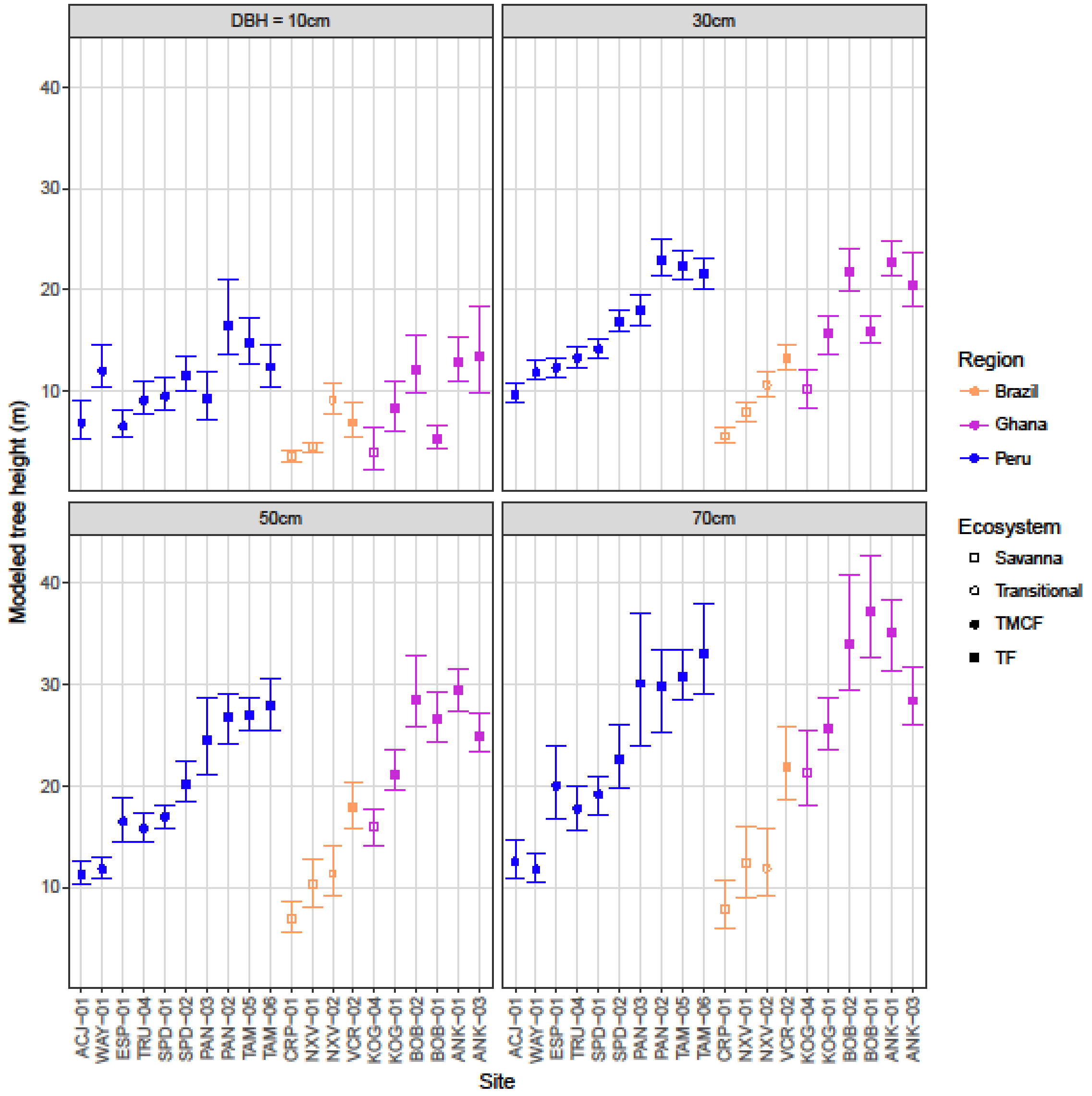


Figure S 33. Predicted tree heights and 95% CIs for for 10cm (a), 30cm (b), 50cm (c), and 70cm (d) DBH trees across sites. Predictions are derived from the tree height vs stem width LMM allometry with random per-species intercepts and slopes. The model is: log(height) ~ (1+log(DBH)|Species) + site*log(DBH). CIs were estimated using a bootstrap algorithm with 100 iterations.

### 6 SI TABLES

Table S 1. Biophysical, ecological, and environmental properties of 20 1-ha study sites.

| Region | Plot | Latitude | Longitude | Elevation* (m) | Slope* (°) | Aspect* (°) | Solar radiation (GJ $m^{-2}$ $yr^{-1}$) | Mean annual air temperature** (°C) | Precipitation (mm $yr^{-1}$) | Soil moisture (%) | Vegetation height* (m) |
|---|---|---|---|---|---|---|---|---|---|---|---|
| Peru | TAM-06 | -12.8309 | -69.2705 | 215 | 2.2 | 169 | 4.8 | 24.4 | 1900 | 35.5 | 28.2 |
| Peru | TAM-05 | -12.8385 | -69.296 | 223 | 4.5 | 186 | 4.8 | 24.4 | 1900 | 21.8 | 27.5 |
| Peru | PAN-02 | -12.6495 | -71.2626 | 595 | 11.5 | 138 | 3.82 | 23 | 2366 | | 24.4 |
| Peru | PAN-03 | -12.6383 | -71.2744 | 859 | 13.7 | 160.5 | | 21.9 | 2835 | | 18.7 |
| Peru | SPD-02 | -13.0491 | -71.5365 | 1494 | 27.1 | 125 | 4.08 | 18.8 | 5302 | 37.3 | 22.8 |
| Peru | SPD-01 | -13.0475 | -71.5423 | 1713 | 30.5 | 117 | 4.36 | 17.4 | 5302 | 37.6 | 14 |
| Peru | TRU-04 | -13.1055 | -71.5893 | 2719 | 21.2 | 118 | 3.49 | 13.5 | 2318 | 37.3 | 15.7 |
| Peru | ESP-01 | -13.1751 | -71.5948 | 2868 | 27.3 | 302 | | 13.1 | 1560 | 24.3 | 16.9 |
| Peru | WAY-01 | -13.1908 | -71.5874 | 3045 | 30.3 | 112 | 3.51 | 11.8 | 1560 | 23.1 | 14.3 |
| Peru | ACJ-01 | -13.1469 | -71.6323 | 3537 | 36.3 | 104 | 4.6 | 9 | 1980 | | 12.5 |
| Ghana | ANK-01 | 5.26728 | -2.69407 | | | | | | 2200 | | |
| Ghana | ANK-03 | 5.271492 | -2.69336 | | | | | | 2200 | | |
| Ghana | BOB-01 | 6.70442 | -1.31857 | 277 | | | | | 1600 | | |
| Ghana | BOB-02 | 6.69104 | -1.33892 | 281 | | | | | 1600 | | |
| Ghana | KOG-01 | 7.26163 | -1.15006 | 229 | | | | | 1200 | | |
| Ghana | KOG-04 | 7.30115 | -1.16493 | 221 | | | | | 1200 | | |
| Brazil | CRP-01 | -14.713 | -52.352 | 372 | | | | | 1400 | | |
| Brazil | NXV-01 | -14.708 | -52.353 | 325 | | | | | 1400 | | |
| Brazil | NXV-02 | -14.702 | -52.352 | 314 | | | | | 1400 | | |
| Brazil | VCR-02 | -14.83 | -52.13 | 294 | | | | | 1400 | | |

| Region | Plot | Number of | basal_area | Number of tree | Mean sampled | Classification | Classification_full | Experimental context |
|---|---|---|---|---|---|---|---|---|

| | | stems | | species | tree height | | | |
|---|---|---|---|---|---|---|---|---|
| Peru | TAM-06 | 660 | 34.09672 | 232 | 22.9 | TF | Tropical Forest | Elevation |
| Peru | TAM-05 | 526 | 26.18826 | 215 | 24.3 | TF | Tropical Forest | Elevation |
| Peru | PAN-02 | 567 | 28.20133 | 169 | 24.5 | TF | Tropical Forest | Elevation |
| Peru | PAN-03 | 682 | 23.96647 | 149 | 17.5 | TF | Tropical Forest | Elevation |
| Peru | SPD-02 | 794 | 31.03799 | 148 | 16.8 | TF | Tropical Forest | Elevation |
| Peru | SPD-01 | 1127 | 43.37792 | 172 | 14.9 | TMCF | Tropical Montane Cloud Forest | Elevation |
| Peru | TRU-04 | 941 | 34.93369 | 53 | 13.6 | TMCF | Tropical Montane Cloud Forest | Elevation |
| Peru | ESP-01 | 842 | 27.61236 | 53 | 12.5 | TMCF | Tropical Montane Cloud Forest | Elevation |
| Peru | WAY-01 | 1166 | 33.78726 | 55 | 12.4 | TMCF | Tropical Montane Cloud Forest | Elevation |
| Peru | ACJ-01 | 856 | 38.25007 | 26 | 10.3 | TMCF | Tropical Montane Cloud Forest | Elevation |
| Ghana | ANK-01 | 452 | 28.02478 | 81 | 26.1 | TF | semi-flooded wet rainforest | Rainfall gradient |
| Ghana | ANK-03 | 483 | 25.8465 | 79 | 24.8 | TF | wet rain forest | Rainfall gradient |
| Ghana | BOB-01 | 455 | 22.87519 | 87 | 21.1 | TF | semi-decidious seasonal forest | Rainfall gradient |
| Ghana | BOB-02 | 772 | 31.11891 | 85 | 24.5 | TF | semi-decidious seasonal forest | Rainfall gradient |
| Ghana | KOG-01 | 178 | 17.46138 | 39 | 20.6 | TF | closed savanna forest | Forest - Savanna Transition |
| Ghana | KOG-04 | 183 | 12.37718 | 28 | 15.3 | Savanna | savanna | Forest - Savanna Transition |
| Brazil | CRP-01 | 1572 | 14.00865 | 80 | 4.6 | Savanna | savanna (cerrado rupestre) | Forest - Savanna Transition |

| | | | | | | | | |
|---|---|---|---|---|---|---|---|---|
| Brazil | NXV-01 | 496 | 7.475151 | 62 | 7.3 | Savanna | woodland savanna (cerrado típico) | Forest - Savanna Transition |
| Brazil | NXV-02 | 1629 | 16.05636 | 107 | 10.3 | Transitional | tall woodland savanna (cerradão) | Forest - Savanna Transition |
| Brazil | VCR-02 | 331 | 12.78606 | 57 | 14.6 | TF | semi-deciduous seasonal forest | Forest - Savanna Transition |

* Peru data derived from high-resolution airborne Light Detection and Ranging (LiDAR) data (Asner et al., 2013;Asner et al., 2014)
** Peru data derived from observations between 6 Feb 2013 and 7 Jan 2014

Table S 2. List of species sampled across plots in this study. Color indicates region: blue = Peru, Beige = Brazil, Purple = Ghana.

| **Genus** | **Species** | **ACJ 01** | **WAY 01** | **ESP 01** | **TRU 04** | **SPD 01** | **SPD 02** | **PAN 03** | **PAN 02** | **TAM 05** | **TAM 06** | **CRP 01** | **NXV 01** | **NXV 02** | **VCR 02** | **KOG 04** | **KOG 01** | **BOB 02** | **BOB 01** | **ANK 01** | **ANK 03** |
|---|---|---|---|---|---|---|---|---|---|---|---|---|---|---|---|---|---|---|---|---|---|
| *Afzelia* | *africana* | | | | | | | | | | | | | | | | X | | | | |
| *Albizia* | *zygia* | | | | | | | | | | | | | | | | | X | X | | |
| *Alchornea* | *indet* | | | | | | X | | | | | | | | | | | | | | |
| | *latifolia* | | | | | X | X | | | | | | | | | | | | | | |
| | *pearcei* | | | | X | | | | | | | | | | | | | | | | |
| *Alzatea* | *verticillata* | | | | | X | | | | | | | | | | | | | | | |
| *Amaioua* | *guianensis* | | | | | | | | | | | | | | X | | | | | | |
| *Anacardium* | *occidentale* | | | | | | | | | | | X | | | | | | | | | |
| *Anogeissus* | *leiocarpus* | | | | | | | | | | | | | | | X | X | | | | |
| *Anthocleista* | *vogelii* | | | | | | | | | | | | | | | | | | | X | |
| *Anthodiscus* | *peruanus* | | | | | | | | | X | | | | | | | | | | | |
| *Apuleia* | *leiocarpa* | | | | | | | | | | | | | | X | | | | | | |
| *Aspidosperma* | *macrocarpon* | | | | | | | | | | | X | | | | | | | | | |
| | *multiflorum* | | | | | | | | | | | | | X | | | | | | | |
| | *tomentosum* | | | | | | | | | | | | X | | | | | | | | |
| *Astrocaryum* | *gratum* | | | | | | | | | | X | | | | | | | | | | |
| *Astronium* | *fraxinifolium* | | | | | | | | | | | | X | | | | | | | | |
| *Axinaea* | *pennellii* | | X | | | | | | | | | | | | | | | | | | |
| *Bellucia* | *pentamera* | | | | | | | | X | | | | | | | | | | | | |
| *Berlinia* | *tomentella* | | | | | | | | | | | | | | | | | | | X | X |
| *Bertholletia* | *excelsa* | | | | | | | | | X | | | | | | | | | | | |
| *Bixa* | *arborea* | | | | | | | | | X | | | | | | | | | | | |
| *Blighia* | *sapida* | | | | | | | | | | | | | | | | | | | X | |
| *Bombax* | *buonopozense* | | | | | | | | | | | | | | | | X | | | | |
| *Bowdichia* | *virgilioides* | | | | | | | | | | | X | X | | | | | | | | |
| *Bridelia* | *ferruginea* | | | | | | | | | | | | | | | X | | | | | |
| *Brosimum* | *alicastrum* | | | | | | | | | | X | | | | | | | | | | |
| | *lactescens* | | | | | | | | | X | | | | | | | | | | | |
| | *rubescens* | | | | | | | | | | | | | | X | | | | | | |
| *Brunellia* | *inermis* | | X | | | | | | | | | | | | | | | | | | |
| | *stenoptera* | | | | | | X | | | | | | | | | | | | | | |
| *Buchenavia* | *capitata* | | | | | | | | | | | | | | X | | | | | | |
| | *tomentosa* | | | | | | | | | | | | X | | | | | | | | |
| *Byrsonima* | *coccolobifolia* | | | | | | | | | | | | X | | | | | | | | |
| | *pachyphylla* | | | | | | | | | | | | X | | | | | | | | |

| Genus | Species | ACJ 01 | WAY 01 | ESP 01 | TRU 04 | SPD 01 | SPD 02 | PAN 03 | PAN 02 | TAM 05 | TAM 06 | CRP 01 | NXV 01 | NXV 02 | VCR 02 | KOG 04 | KOG 01 | BOB 02 | BOB 01 | ANK 01 | ANK 03 |
|---|---|---|---|---|---|---|---|---|---|---|---|---|---|---|---|---|---|---|---|---|---|
| *Calophyllum* | *brasiliense* | | | | | | | | | X | | | | | | | | | | | |
| *Carapa* | *procera* | | | | | | | | | | | | | | | | | | | | X |
| *Caryocar* | *amygdaliforme* | | | | | | X | X | | | | | | | | | | | | | |
| | *brasiliense* | | | | | | | | | | | | X | | | | | | | | |
| | *indet* | | | | | X | | | | | | | | | | | | | | | |
| *Cecropia* | *angustifolia* | | | | | X | X | | | | | | | | | | | | | | |
| *Cedrelinga* | *cateniformis* | | | | | | | | | X | | | | | | | | | | | |
| *Ceiba* | *pentandra* | | | | | | | | | | | | | | | | X | | X | | |
| *Celtis* | *mildbraedii* | | | | | | | | | | | | | | | | | X | X | | |
| | *schippii* | | | | | | | | | | X | | | | | | | | | | |
| | *zenkeri* | | | | | | | | | | | | | | | | | X | X | | |
| *Chaetocarpus* | *echinocarpus* | | | | | | | | | | | | | X | X | | | | | | |
| *Cheiloclinium* | *cognatum* | | | | | | | | | | | | | | X | | | | | | |
| *Chimarrhis* | *hookeri* | | | | | | | | | | X | | | | | | | | | | |
| *Chrysophyllum* | *indet* | | | | | X | | | | | | | | | | | | | | X | |
| | *subnudum* | | | | | | | | | | | | | | | | | | | X | |
| *Cinchona* | *calisaya* | | | | X | | | | | | | | | | | | | | | | |
| | *indet* | | | | | | X | | | | | | | | | | | | | | |
| *Clarisia* | *racemosa* | | | | | | | | | X | X | | | | | | | | | | |
| *Clethra* | *cuneata* | X | X | | | | | | | | | | | | | | | | | | |
| *Clusia* | *alata* | | X | X | X | | | | | | | | | | | | | | | | |
| | *elliptica* | | | X | | X | | | | | | | | | | | | | | | |
| | *flaviflora* | | X | | | | | | | | | | | | | | | | | | |
| | *indet* | | | | X | | | | | | | | | | | | | | | | |
| | *thurifera* | | | | | X | | | | | | | | | | | | | | | |
| | *trochiformis* | | | X | | | | | | | | | | | | | | | | | |
| *Cola* | *caricifolia* | | | | | | | | | | | | | | | | | | X | | |
| | *gigantea* | | | | | | | | | | | | | | | | X | | X | | X |
| | *nitida* | | | | | | | | | | | | | | | | | | | X | |
| *Coussapoa* | *cf. manuensis* | | | | | | | | X | | | | | | | | | | | | |
| | *villosa* | | | | | | X | | | | | | | | | | | | | | |
| *Cyathea* | *delgadii* | | | | X | | | | | | | | | | | | | | | | |
| | *indet* | | | | | X | X | | | | | | | | | | | | | | |
| *Cylicodiscus* | *gabunensis* | | | | | | | | | | | | | | | | | X | | | |
| *Cynometra* | *ananta* | | | | | | | | | | | | | | | | | | | X | X |
| *Dacryodes* | *klaineana* | | | | | | | | | | | | | | | | X | | | X | |
| *Dalbergia* | *miscolobium* | | | | | | | | | | | X | X | | | | | | | | |
| *Davilla* | *elliptica* | | | | | | | | | | | | X | | | | | | | | |
| *Dendropanax* | *indet* | | | | | X | | | | | | | | | | | | | | | |
| *Dialium* | *aubrevillei* | | | | | | | | | | | | | | | | | | | X | |

| Genus | Species | ACJ 01 | WAY 01 | ESP 01 | TRU 04 | SPD 01 | SPD 02 | PAN 03 | PAN 02 | TAM 05 | TAM 06 | CRP 01 | NXV 01 | NXV 02 | VCR 02 | KOG 04 | KOG 01 | BOB 02 | BOB 01 | ANK 01 | ANK 03 |
|---|---|---|---|---|---|---|---|---|---|---|---|---|---|---|---|---|---|---|---|---|---|
| *Dimorphandra* | *mollis* | | | | | | | | | | | | X | | | | | | | | |
| *Diospyros* | *mespiliformis* | | | | | | | | | | | | | | | | X | | | | |
| | *sanza-minika* | | | | | | | | | | | | | | | | | | | X | |
| *Dipteryx* | *alata* | | | | | | | | | | | X | | | | | | | | | |
| *Drypetes* | *gilgiana* | | | | | | | | | | | | | | | | | | | X | |
| *Duguetia* | *staudtii* | | | | | | | | | | | | | | | | | | | | X |
| *Elaeagia* | *indet* | | | | | X | | | | | | | | | | | | | | | |
| *Emmotum* | *nitens* | | | | | | | | | | | | X | X | | | | | | | |
| *Endlicheria* | *formosa* | | | | | | | | | | X | | | | | | | | | | |
| | *indet* | | | | | | X | | | | | | | | | | | | | | |
| *Entandrophragma* | *angolense* | | | | | | | | | | | | | | | | | X | | | |
| *Ephedranthus* | *parviflorus* | | | | | | | | | | | | | | X | | | | | | |
| *Eriotheca* | *gracilipes* | | | | | | | | | | | | X | X | | | | | | | |
| *Erythrophleum* | *suaveolens* | | | | | | | | | | | | | | | | X | | | | |
| *Erythroxylum* | *suberosum* | | | | | | | | | | | X | | | | | | | | | |
| *Eschweilera* | *coriacea* | | | | | | | | | X | | | | | | | | | | | |
| *Eugenia* | *aurata* | | | | | | | | | | | X | | | | | | | | | |
| *Euplassa* | *inaequalis* | | | | | | | | | | | | X | | | | | | | | |
| *Faramea* | *torquata* | | | | | | | X | | | | | | | | | | | | | |
| *Ficus* | *americana* | | | | | X | | | | | | | | | | | | | | | |
| | *maxima* | | | | | | X | | | | | | | | | | | | | | |
| *Freziera* | *parva* | | X | | X | | | | | | | | | | | | | | | | |
| *Funtumia* | *elastica* | | | | | | | | | | | | | | | | | X | X | | |
| *Garcinia* | *smeathmanii* | | | | | | | | | | | | | | | | | | | X | |
| *Gordonia* | *fruticosa* | | | | X | | | | | | | | | | | | | | | | |
| *Greenwayodendron* | *oliveri* | | | | | | | | | | | | | | | | | | | X | |
| *Guapira* | *graciliflora* | | | | | | | | | | | | X | | | | | | | | |
| *Guarea* | *kunthiana* | | | | | | X | | | | | | | | | | | | | | |
| *Guatteria* | *glauca* | | | | | X | X | | | | | | | | | | | | | | |
| *Hannoa* | *klaineana* | | | | | | | | | | | | | | | | | | X | | |
| *Hebepetalum* | *humiriifolium* | | | | | | | | | X | | | | | | | | | | | |
| *Hedyosmum* | *cuatrecazanum* | | | | | X | | | | | | | | | | | | | | | |
| | *goudotianum* | | | | X | | | | | | | | | | | | | | | | |
| | *maximum* | | X | | | | | | | | | | | | | | | | | | |
| | *indet* | X | | | | | | | | | | | | | | | | | | | |
| | *translucidum* | | | X | | | | | | | | | | | | | | | | | |
| *Heisteria* | *ovata* | | | | | | | | | | | | | X | | | | | | | |
| *Heliocarpus* | *americanus* | | | | | | X | | | | | | | | | | | | | | |
| *Heritiera* | *utilis* | | | | | | | | | | | | | | | | | | | X | X |
| *Hesperomeles* | *ferruginea* | | X | | | | | | | | | | | | | | | | | | |

| Genus | Species | ACJ 01 | WAY 01 | ESP 01 | TRU 04 | SPD 01 | SPD 02 | PAN 03 | PAN 02 | TAM 05 | TAM 06 | CRP 01 | NXV 01 | NXV 02 | VCR 02 | KOG 04 | KOG 01 | BOB 02 | BOB 01 | ANK 01 | ANK 03 |
|---|---|---|---|---|---|---|---|---|---|---|---|---|---|---|---|---|---|---|---|---|---|
| *Heteropterys* | *byrsonimifolia* | | | | | | | | | | | X | | | | | | | | | |
| *Hevea* | *brasiliensis* | | | | | | | | X | | | | | | | | | | | | |
| | *guianensis* | | | | | | | X | | | | | | | | | | | | | |
| *Hieronyma* | *alchorneoides* | | | | | | X | | | | | | | | | | | | | | |
| *Hirtella* | *glandulosa* | | | | | | | | | | | | | X | | | | | | | |
| *Huberodendron* | *swietenioides* | | | | | | | | | X | | | | | | | | | | | |
| *Hymenaea* | *courbaril* | | | | | | | | | | | | | | X | | | | | | |
| | *stigonocarpa* | | | | | | | | | | | X | | | | | | | | | |
| *Hymenostegia* | *afzelii* | | | | | | | | | | | | | | | | | | X | | |
| *Ilex* | *gabrielleana* | | | | | X | | | | | | | | | | | | | | | |
| *Indet* | *indet* | | X | | | | | | | | | | | | | | | | | | |
| *Inga* | *indet* | | | | | X | | | | | | | | | | | | | | | |
| | *thibaudiana* | | | | | | | | X | | | | | | | | | | | | |
| *Iriartea* | *deltoidea* | | | | | | | | | X | X | | | | | | | | | | |
| *Jacaranda* | *copaia* | | | | | | | | | X | | | | | | | | | | | |
| | *glabra* | | | | | | | | X | | | | | | | | | | | | |
| *Khaya* | *grandifoliola* | | | | | | | | | | | | | | | | X | | | | |
| | *indet* | | | | | | | | | | | | | | | | X | | | | |
| *Kielmeyera* | *rubriflora* | | | | | | | | | | | X | X | | | | | | | | |
| *Lafoensia* | *pacari* | | | | | | | | | | | X | | | | | | | | | |
| *Lannea* | *velutina* | | | | | | | | | | | | | | | X | X | | | | |
| | *welwitschii* | | | | | | | | | | | | | | | | | X | | | |
| *Leonia* | *glycycarpa* | | | | | | | | | X | | | | | | | | | | | |
| *Licania* | *heteromorpha* | | | | | | | | | X | | | | | | | | | | | |
| *Licaria* | *indet* | | | | | | X | | | | | | | | | | | | | | |
| *Lissocarpa* | *indet* | | | | | X | | | | | | | | | | | | | | | |
| *Lovoa* | *trichilioides* | | | | | | | | | | | | | | | | | | | X | |
| *Mabea* | *fistulifera* | | | | | | | | | | | | | | X | | | | | | |
| | *nitida* | | | | | | | | | | X | | | | | | | | | | |
| *Macrocnemum* | *roseum* | | | | | | | | | | X | | | | | | | | | | |
| *Magonia* | *pubescens* | | | | | | | | | | | X | X | | | | | | | | |
| *Maprounea* | *guianensis* | | | | | | | | | | | | | X | | | | | | | |
| *Maranthes* | *glabra* | | | | | | | | | | | | | | | | | | | X | |
| *Marila* | *laxiflora* | | | | | | | | X | | | | | | | | | | | | |
| *Matayba* | *guianensis* | | | | | | | | | | | | | X | | | | | | | |
| *Meliosma* | *frondosa* | | | X | | | | | | | | | | | | | | | | | |
| | *indet* | | | | | X | | | | | | | | | | | | | | | |
| *Meriania* | *indet* | | | | | X | | | | | | | | | | | | | | | |
| *Mezilaurus* | *crassiramea* | | | | | | | | | | | X | X | | | | | | | | |
| *Miconia* | *alpina* | X | | | | | | | | | | | | | | | | | | | |

| Genus | Species | ACJ 01 | WAY 01 | ESP 01 | TRU 04 | SPD 01 | SPD 02 | PAN 03 | PAN 02 | TAM 05 | TAM 06 | CRP 01 | NXV 01 | NXV 02 | VCR 02 | KOG 04 | KOG 01 | BOB 02 | BOB 01 | ANK 01 | ANK 03 |
|---|---|---|---|---|---|---|---|---|---|---|---|---|---|---|---|---|---|---|---|---|---|
| | *indet* | | | | X | | | | | | | | | | | | | | | | |
| | *micropetala* | | | X | | | | | | | | | | | | | | | | | |
| | *setulosa* | X | | | | | | | | | | | | | | | | | | | |
| *Micropholis* | *guyanensis* | | | | | | | X | | | | | | | | | | | | | |
| *Milicia* | *excelsa* | | | | | | | | | | | | | | | | X | | | | |
| *Millettia* | *rhodantha* | | | | | | | | | | | | | | | | | | | X | |
| *Mollinedia* | *lanceolata* | | | | | | X | | | | | | | | | | | | | | |
| *Musanga* | *cecropioides* | | | | | | | | | | | | | | | | | | X | | |
| *Myrcia* | *splendens* | | | | | | | | | | | | | X | | | | | | | |
| *Myroxylon* | *balsamum* | | | | | | | | | | X | | | | | | | | | | |
| *Myrsine* | *andina* | X | | | X | | | | | | | | | | | | | | | | |
| | *coriacea* | | X | X | X | | | | | | | | | | | | | | | | |
| | *indet* | | X | | | | | | | | | | | | | | | | | | |
| *Nesogordonia* | *papaverifera* | | | | | | | | | | | | | | | | | X | X | | |
| *Ocotea* | *bofo* | | | | | | | | X | X | X | | | | | | | | | | |
| | *hoehneii* | | | | | | | | | | | | | | X | | | | | | |
| | *indet* | | | | X | X | | | | | | | | | | | | | | | |
| | *insularis* | | | | | | | X | X | | | | | | | | | | | | |
| *Ongokea* | *gore* | | | | | | | | | | | | | | | | | | | | X |
| *Oreopanax* | *microflorous* | | | X | | | | | | | | | | | | | | | | | |
| *Ormosia* | *panamensis* | | | | | | | | | | X | | | | | | | | | | |
| *Otoba* | *parvifolia* | | | | | | | | | | X | | | | | | | | | | |
| *Ouratea* | *indet* | | | | | | | | | X | | | | | | | | | | | |
| *Parinari* | *parilis* | | | | | X | | | | | | | | | | | | | | | |
| *Parkia* | *bicolor* | | | | | | | | | | | | | | | | | | | | X |
| *Pentadesma* | *butyracea* | | | | | | | | | | | | | | | | | | | X | |
| *Perrottetia* | *distichophylla* | | | | | | X | | | | | | | | | | | | | | |
| *Persea* | *indet* | | | | X | | | | | | | | | | | | | | | | |
| | *ruizii* | X | | | | | | | | | | | | | | | | | | | |
| *Petersianthus* | *macrocarpus* | | | | | | | | | | | | | | | | | X | | | |
| *Piptadeniastrum* | *africanum* | | | | | | | | | | | | | | | | | X | | | |
| *Polylepis* | *pauta* | X | | | | | | | | | | | | | | | | | | | |
| *Pourouma* | *bicolor* | | | | | | X | | X | | | | | | | | | | | | |
| | *cecropiifolia* | | | | | | | | | | X | | | | | | | | | | |
| | *guianensis* | | | | | | | | | X | | | | | | | | | | | |
| | *minor* | | | | | | | | X | X | | | | | | | | | | | |
| | *mollis* | | | | | | | X | X | | | | | | | | | | | | |
| *Pouteria* | *alnifolia* | | | | | | | | | | | | | | | | X | | | | |
| | *ramiflora* | | | | | | | | | | | | X | | | | | | | | |
| | *torta* | | | | | X | | | | X | X | | | | | | | | | | |

| Genus | Species | ACJ 01 | WAY 01 | ESP 01 | TRU 04 | SPD 01 | SPD 02 | PAN 03 | PAN 02 | TAM 05 | TAM 06 | CRP 01 | NXV 01 | NXV 02 | VCR 02 | KOG 04 | KOG 01 | BOB 02 | BOB 01 | ANK 01 | ANK 03 |
|---|---|---|---|---|---|---|---|---|---|---|---|---|---|---|---|---|---|---|---|---|---|
| *Protium* | *heptaphyllum* | | | | | | | | | | | | | X | | | | | | | |
| | *montanum* | | | | | X | | | | | | | | | | | | | | | |
| *Protomegabaria* | *stapfiana* | | | | | | | | | | | | | | | | | | | | X |
| *Prunus* | *debilis* | | | | X | | | | | | | | | | | | | | | | |
| | *indet* | | | | | X | | | | | | | | | | | | | | | |
| | *integrifolia* | | X | X | | | | | | | | | | | | | | | | | |
| *Pseudobombax* | *longiflorum* | | | | | | | | | | | X | X | X | | | | | | | |
| *Pseudolmedia* | *laevigata* | | | | | | | | | X | | | | | | | | | | | |
| | *laevis* | | | | | | | | | X | | | | | | | | | | | |
| | *rigida* | | | | | X | X | | X | | | | | | | | | | | | |
| *Pterocarpus* | *erinaceus* | | | | | | | | | | | | | | | X | | | | | |
| | *indet* | | | | | | | | | X | | | | | | | | | | | |
| | *rohrii* | | | | | | | | | | X | | | | | | | | | | |
| *Pterodon* | *pubescens* | | | | | | | | | | | X | | | | | | | | | |
| *Pterygota* | *macrocarpa* | | | | | | | | | | | | | | | | | X | X | | |
| *Pycnanthus* | *angolensis* | | | | | | | | | | | | | | | | | | X | | |
| *Qualea* | *grandiflora* | | | | | | | | | | | X | X | | | | | | | | |
| | *multiflora* | | | | | | | | | | | X | X | | | | | | | | |
| | *paraensis* | | | | | | | X | | | | | | | | | | | | | |
| | *parviflora* | | | | | | | | | | | X | X | | | | | | | | |
| *Rauvolfia* | *indet* | | | | | | X | | | | | | | | | | | | | | |
| *Retrophyllum* | *rospigliosii* | | | | | X | | | | | | | | | | | | | | | |
| *Rinorea* | *viridifolia* | | | | | | | | | | X | | | | | | | | | | |
| *Roucheria* | *punctata* | | | | | | | | | X | | | | | | | | | | | |
| *Roupala* | *montana* | | | | | | | | | | | | X | | | | | | | | |
| *Salvertia* | *convallariodora* | | | | | | | | | | | | X | | | | | | | | |
| *Sapium* | *glandulosum* | | | | | | X | | | | | | | | | | | | | | |
| *Scaphopetalum* | *amoenum* | | | | | | | | | | | | | | | | | | | X | |
| *Scheelea* | *cephalotes* | | | | | | | | | | X | | | | | | | | | | |
| *Schizocalyx* | *obovatus* | | | | | | | X | | | | | | | | | | | | | |
| *Sclerolobium* | *bracteosum* | | | | | | | | | X | | | | | | | | | | | |
| *Scottellia* | *klaineana* | | | | | | | | | | | | | | | | | | | X | |
| *Scytopetalum* | *tieghemii* | | | | | | | | | | | | | | | | | | | | X |
| *Senefeldera* | *inclinata* | | | | | | | X | X | | | | | | | | | | | | |
| *Sloanea* | *guianensis* | | | | | | | X | | | | | | | | | | | | | |
| | *meianthera* | | | | | | | X | | | | | | | | | | | | | |
| | *sinemariensis* | | | | | | | X | | | | | | | | | | | | | |
| *Socratea* | *exorrhiza* | | | | | | | | | | X | | | | | | | | | | |
| *Sorocea* | *klotzschiana* | | | | | | | | | | | | | X | | | | | | | |
| *Spathodea* | *campanulata* | | | | | | | | | | | | | | | | X | | | | |

| Genus | Species | ACJ 01 | WAY 01 | ESP 01 | TRU 04 | SPD 01 | SPD 02 | PAN 03 | PAN 02 | TAM 05 | TAM 06 | CRP 01 | NXV 01 | NXV 02 | VCR 02 | KOG 04 | KOG 01 | BOB 02 | BOB 01 | ANK 01 | ANK 03 |
|---|---|---|---|---|---|---|---|---|---|---|---|---|---|---|---|---|---|---|---|---|---|
| *Staphylea* | *occidentalis* | | | | | | X | | | | | | | | | | | | | | |
| *Sterculia* | *oblonga* | | | | | | | | | | | | | | | | | X | X | | |
| | *rhinopetala* | | | | | | | | | | | | | | | | | X | X | | |
| | *tragacantha* | | | | | | | | | | | | | | | | X | | | | |
| *Strephonema* | *pseudocola* | | | | | | | | | | | | | | | | | | | X | X |
| *Strombosia* | *glaucescens* | | | | | | | | | | | | | | | | | | X | | |
| | *pustulata* | | | | | | | | | | | | | | | | | | | X | |
| *Strychnos* | *pseudoquina* | | | | | | | | | | | | X | | | | | | | | |
| *Swartzia* | *arborescens* | | | | | | | | | | X | | | | | | | | | | |
| *Symphonia* | *globulifera* | | | | | | | | | | X | | | | | | | | | | |
| *Symplocos* | *baehnii* | X | | | | | | | | | | | | | | | | | | | |
| | *quitensis* | X | | | | | | | | | | | | | | | | | | | |
| | *reflexa* | | | X | | | | | | | | | | | | | | | | | |
| *Tabebuia* | *serratifolia* | | | | | | | | | | | | | | X | | | | | | |
| *Tachigali* | *aurea* | | | | | | | | | | | X | | | | | | | | | |
| | *indet* | | | | | | X | | | | | | | | | | | | | | |
| | *polyphylla* | | | | | | | | | X | | | | | | | | | | | |
| | *setifera* | | | | | | | X | | | | | | | | | | | | | |
| | *vulgaris* | | | | | | | | | | | | X | X | | | | | | | |
| *Tapirira* | *guianensis* | | | | | | | | X | | | | | X | | | | | | | |
| | *indet* | | | | | X | X | | | | | | | | | | | | | | |
| | *obtusa* | | | | | | X | | | | | | | | | | | | | | |
| *Terminalia* | *argentea* | | | | | | | | | | | | X | | | | | | | | |
| | *indet* | | | | | | | | | | | | | | | X | | | | | |
| | *schimperiana* | | | | | | | | | | | | | | | X | | | | | |
| | *superba* | | | | | | | | | | | | | | | | | X | | | |
| *Tetragastris* | *altissima* | | | | | | | | | | | | | | X | | | | | | |
| *Trichilia* | *prieuriana* | | | | | | | | | | | | | | | | | | X | | |
| *Triplochiton* | *scleroxylon* | | | | | | | | | | | | | | | | | | X | | |
| *Turraeanthus* | *africanus* | | | | | | | | | | | | | | | | | X | | | |
| *Uapaca* | *guineensis* | | | | | | | | | | | | | | | | | | | X | X |
| *Vatairea* | *macrocarpa* | | | | | | | | | | | X | | X | | | | | | | |
| *Virola* | *sebifera* | | | | | | X | | | | | | | | | | | | | | |
| *Vismia* | *gracilis* | | | | | | X | | | | | | | | | | | | | | |
| *Vitellaria* | *paradoxa* | | | | | | | | | | | | | | | X | | | | | |
| *Vitex* | *ferruginea* | | | | | | | | | | | | | | | | | | | X | |
| *Vochysia* | *haenkeana* | | | | | | | | | | | | | X | | | | | | | |
| | *rufa* | | | | | | | | | | | | X | | | | | | | | |
| *Weinmannia* | *bangii* | | X | X | X | | | | | | | | | | | | | | | | |
| | *crassifolia* | | X | X | | | | | | | | | | | | | | | | | |

| Genus | Species | ACJ 01 | WAY 01 | ESP 01 | TRU 04 | SPD 01 | SPD 02 | PAN 03 | PAN 02 | TAM 05 | TAM 06 | CRP 01 | NXV 01 | NXV 02 | VCR 02 | KOG 04 | KOG 01 | BOB 02 | BOB 01 | ANK 01 | ANK 03 |
|---|---|---|---|---|---|---|---|---|---|---|---|---|---|---|---|---|---|---|---|---|---|
| | *fagaroides* | X | | | | | | | | | | | | | | | | | | | |
| | *pinnata* | | | | | X | | | | | | | | | | | | | | | |
| | *reticulata* | | X | | X | | | | | | | | | | | | | | | | |
| *Xylopia* | *aromatica* | | | | | | | | | | | | | X | | | | | | | |
| *Zanthoxylum* | *acreanum* | | | | | | | | | | X | | | | | | | | | | |

Table S 3. Number of individual trees observed across DBH and crown exposure classes.

| | (-0.001,0.2] | **(0.2,0.4]** | **(0.4,0.6]** | **(0.6,0.8]** | **(0.8,1]** |
|---|---|---|---|---|---|
| [0,20] | 35 | 6 | 11 | 6 | 16 |
| (20,40] | 63 | 27 | 38 | 33 | 73 |
| (40,60] | 15 | 9 | 12 | 26 | 77 |
| (60,80] | 2 | 2 | 1 | 6 | 20 |
| (80,100] | 0 | 0 | 0 | 1 | 15 |
| (100,120] | 0 | 0 | 1 | 0 | 3 |
| (120,140] | 0 | 0 | 0 | 0 | 1 |

Table S 4. Number of individual trees observed across height and crown exposure classes.

| | (-0.001,0.2] | **(0.2,0.4]** | **(0.4,0.6]** | **(0.6,0.8]** | **(0.8,1]** |
|---|---|---|---|---|---|
| [0,5] | 1 | 0 | 1 | 1 | 1 |
| (5,10] | 14 | 3 | 4 | 1 | 6 |
| (10,15] | 29 | 10 | 22 | 10 | 19 |
| (15,20] | 31 | 11 | 10 | 16 | 43 |
| (20,25] | 21 | 12 | 12 | 21 | 44 |
| (25,30] | 16 | 4 | 8 | 17 | 55 |
| (30,35] | 5 | 3 | 5 | 5 | 31 |
| (35,40] | 0 | 1 | 1 | 1 | 16 |
| (40,45] | 0 | 0 | 0 | 0 | 3 |

Table S 5. Likelihood ratio test of log(crown width) vs log(dbh) scaling models including different site effects.

| | Df | AIC | BIC | logLik | deviance | Chisq | Chi Df | Pr(>Chisq) |
|---|---|---|---|---|---|---|---|---|
| No site | 6 | -845.15 | -815.51 | 428.57 | -857.15 | | | |
| Site slope | 25 | -914.58 | -791.10 | 482.29 | -964.58 | 107.43 | 19 | 0.0000 |
| Site intercept & slope | 44 | -913.12 | -695.79 | 500.56 | -1001.12 | 36.54 | 19 | 0.0091 |

Table S 6. LMMs of crown depth and relative depth versus tree size and crown exposure (Models 10 - 15). The fourth relative depth model below includes site as a random effect.

| | ***Dependent variable:*** | | | | | | | | | |
|---|---|---|---|---|---|---|---|---|---|---|
| | reldepth | | | | log10(reldepth) | | depth | | log10(depth) | |
| | (1) | (2) | (3) | (4) | (5) | (6) | (7) | (8) | (9) | (10) |
| $h_{tree}$ | 0.298$^{***}$ | 0.253$^{***}$ | 0.194$^{***}$ | 0.284$^{***}$ | | | 0.572$^{***}$ | 0.555$^{***}$ | | |
| | (0.063) | (0.075) | (0.074) | (0.068) | | | (0.046) | (0.059) | | |
| $h_{tree}{}^2$ | | | -0.055 | -0.067$^{*}$ | | | | | | |
| | | | (0.040) | (0.038) | | | | | | |
| $r_{stem}$ | -0.336$^{***}$ | -0.304$^{***}$ | -0.316$^{***}$ | -0.271$^{***}$ | | | 0.119$^{***}$ | 0.136$^{***}$ | | |
| | (0.045) | (0.066) | (0.068) | (0.064) | | | (0.033) | (0.052) | | |
| ce | | -0.175$^{***}$ | | -0.186$^{***}$ | | -0.146$^{***}$ | | -0.222$^{***}$ | | -0.152$^{***}$ |
| | | (0.051) | | (0.049) | | (0.032) | | (0.040) | | (0.024) |
| $\log(h_{tree})$ | | | | | 0.472$^{***}$ | 0.396$^{***}$ | | | 0.697$^{***}$ | 0.624$^{***}$ |
| | | | | | (0.064) | (0.100) | | | (0.048) | (0.073) |
| $\log(r_{stem})$ | | | | | -0.422$^{***}$ | -0.306$^{***}$ | | | 0.149$^{***}$ | 0.283$^{***}$ |
| | | | | | (0.049) | (0.078) | | | (0.037) | (0.057) |
| siteACJ01 | -0.262 | | | | | | -0.018 | | | |
| | (0.166) | | | | | | (0.122) | | | |
| siteWAY01 | -0.576$^{***}$ | -0.642$^{*}$ | -0.647$^{*}$ | | -0.130$^{**}$ | | -0.229$^{**}$ | -0.323 | -0.073$^{*}$ | |
| | (0.136) | (0.352) | (0.357) | | (0.052) | | (0.100) | (0.277) | (0.038) | |
| siteESP01 | -0.443$^{***}$ | | | | -0.084 | | 0.024 | | -0.003 | |
| | (0.136) | | | | (0.053) | | (0.101) | | (0.039) | |
| siteTRU04 | -0.413$^{***}$ | -0.450$^{***}$ | -0.462$^{***}$ | | -0.100$^{**}$ | 0.051 | -0.282$^{***}$ | -0.281$^{***}$ | -0.115$^{***}$ | -0.011 |
| | (0.119) | (0.135) | (0.141) | | (0.050) | (0.096) | (0.088) | (0.105) | (0.037) | (0.070) |
| siteSPD01 | -0.333$^{***}$ | | | | -0.096$^{*}$ | | -0.278$^{***}$ | | -0.098$^{***}$ | |
| | (0.112) | | | | (0.050) | | (0.082) | | (0.037) | |
| siteSPD02 | -0.409$^{***}$ | | | | -0.106$^{**}$ | | -0.264$^{***}$ | | -0.102$^{***}$ | |
| | (0.110) | | | | (0.051) | | (0.081) | | (0.038) | |
| sitePAN03 | -0.116 | -0.123 | -0.136 | | -0.008 | 0.177$^{*}$ | -0.323$^{***}$ | -0.275$^{**}$ | -0.110$^{**}$ | 0.032 |
| | (0.159) | (0.159) | (0.163) | | (0.059) | (0.102) | (0.117) | (0.124) | (0.044) | (0.075) |
| sitePAN02 | -0.212 | -0.224 | -0.258 | | -0.047 | 0.136 | -0.387$^{***}$ | -0.317$^{***}$ | -0.118$^{***}$ | 0.032 |
| | (0.157) | (0.156) | (0.161) | | (0.060) | (0.103) | (0.116) | (0.122) | (0.045) | (0.075) |
| siteTAM05 | 0.245$^{*}$ | 0.210$^{*}$ | 0.231$^{*}$ | | 0.035 | 0.225$^{**}$ | -0.114 | -0.119 | -0.060 | 0.075 |
| | (0.127) | (0.125) | (0.129) | | (0.055) | (0.100) | (0.094) | (0.098) | (0.041) | (0.073) |
| siteTAM06 | -0.188 | -0.217 | -0.178 | | -0.013 | 0.171$^{*}$ | -0.116 | -0.155 | -0.064 | 0.069 |
| | (0.139) | (0.138) | (0.141) | | (0.057) | (0.101) | (0.102) | (0.108) | (0.042) | (0.074) |

| | | | | | | | | | | |
|---|---|---|---|---|---|---|---|---|---|---|
| siteCRP01 | -0.163 | | | | 0.117$^{**}$ | | -0.304$^{***}$ | | -0.081$^{**}$ | |
| | (0.143) | | | | (0.054) | | (0.105) | | (0.040) | |
| siteNXV01 | -0.358$^{***}$ | | | | 0.016 | | -0.218$^{**}$ | | -0.040 | |
| | (0.124) | | | | (0.050) | | (0.092) | | (0.037) | |
| siteNXV02 | 0.751$^{***}$ | | | | 0.223$^{***}$ | | 0.348$^{***}$ | | 0.104$^{**}$ | |
| | (0.150) | | | | (0.056) | | (0.111) | | (0.041) | |
| siteVCR02 | 0.863$^{***}$ | | | | 0.249$^{***}$ | | 0.832$^{***}$ | | 0.147$^{***}$ | |
| | (0.161) | | | | (0.058) | | (0.119) | | (0.043) | |
| siteKOG04 | 0.407$^{*}$ | 0.468$^{**}$ | 0.306 | | 0.156$^{**}$ | 0.355$^{***}$ | 0.787$^{***}$ | 0.953$^{***}$ | 0.133$^{***}$ | 0.284$^{***}$ |
| | (0.221) | (0.229) | (0.230) | | (0.069) | (0.108) | (0.163) | (0.179) | (0.051) | (0.079) |
| siteKOG01 | 0.153 | 0.169 | 0.095 | | 0.085 | 0.275$^{***}$ | 0.387$^{***}$ | 0.481$^{***}$ | 0.035 | 0.178$^{**}$ |
| | (0.153) | (0.155) | (0.160) | | (0.058) | (0.101) | (0.113) | (0.121) | (0.043) | (0.074) |
| siteBOB02 | 0.239 | 0.258 | 0.227 | | 0.057 | 0.265$^{**}$ | -0.094 | -0.039 | -0.053 | 0.104 |
| | (0.166) | (0.163) | (0.167) | | (0.062) | (0.104) | (0.122) | (0.128) | (0.045) | (0.076) |
| siteBOB01 | -0.025 | 0.023 | 0.007 | | 0.064 | 0.265$^{***}$ | -0.309$^{***}$ | -0.198 | -0.102$^{**}$ | 0.050 |
| | (0.155) | (0.156) | (0.163) | | (0.058) | (0.102) | (0.114) | (0.122) | (0.043) | (0.074) |
| siteANK01 | 0.555$^{***}$ | 0.380$^{***}$ | 0.555$^{***}$ | | 0.140$^{**}$ | 0.297$^{***}$ | 0.542$^{***}$ | 0.355$^{***}$ | 0.059 | 0.164$^{**}$ |
| | (0.133) | (0.138) | (0.134) | | (0.056) | (0.101) | (0.098) | (0.108) | (0.041) | (0.074) |
| siteANK03 | | | | | 0.058 | 0.209$^{**}$ | | | -0.034 | 0.065 |
| | | | | | (0.061) | (0.104) | | | (0.045) | (0.076) |
| Constant | 0.083$^{**}$ | -0.016 | 0.022 | 0.074 | 0.125$^{*}$ | -0.011 | 0.056$^{**}$ | 0.023 | -0.156$^{***}$ | -0.265$^{***}$ |
| | (0.036) | (0.055) | (0.069) | (0.092) | (0.068) | (0.124) | (0.026) | (0.043) | (0.050) | (0.091) |
| Observations | 1,032 | 496 | 496 | 496 | 1,032 | 496 | 1,032 | 496 | 1,032 | 496 |
| Log Likelihood | -1,323.549 | -665.206 | -670.359 | -667.528 | 114.147 | 18.132 | -1,014.271 | -550.800 | 415.193 | 167.750 |
| Akaike Inf. Crit. | 2,695.099 | 1,364.413 | 1,374.717 | 1,351.056 | -180.295 | -2.265 | 2,076.542 | 1,135.601 | -782.386 | -301.500 |
| Bayesian Inf. Crit. | 2,813.641 | 1,435.925 | 1,446.229 | 1,384.709 | -61.753 | 69.247 | 2,195.084 | 1,207.113 | -663.844 | -229.988 |

***Note:*** $^{*}$p$^{**}$p$^{***}$p<0.01

Table S 7. Variance decomposition of the species-mean deviance from the mean LMM scaling predictions.

| | Model from which residuals taken | eco_class | family | genus | plot_code | region | Residual | phylo_sig |
|---|---|---|---|---|---|---|---|---|
| 1 | $\log_{10}$(*crown radius*) ~ (1 + $\log_{10}(r_{stem})$ \| species) + (1 + $\log_{10}(r_{stem})$ \| site) + $\log_{10}(r_{stem})$ | 0.00 | 0.12 | 0.00 | 0.11 | 0.28 | 0.49 | 0.12 |
| 2 | $\log_{10}$(*crown depth*) ~ (1 + $\log_{10}(r_{stem})$ \| species) + (1 + $\log_{10}(r_{stem})$ \| site) + $\log_{10}(r_{stem})$ | 0.35 | 0.00 | 0.00 | 0.19 | 0.06 | 0.40 | 0.00 |
| 3 | $\log_{10}$(*crown depth*) ~ (1 + $\log_{10}$(h) \| species) + (1 + $\log_{10}$(h) \| site) + $\log_{10}$(h) | 0.00 | 0.00 | 0.00 | 0.24 | 0.17 | 0.59 | 0.00 |
| 4 | $\log_{10}$(*sa*) ~ (1 + $\log_{10}(r_{stem})$ \| species) + (1 + $\log_{10}(r_{stem})$ \| site) + $\log_{10}(r_{stem})$ | 0.14 | 0.07 | 0.00 | 0.19 | 0.13 | 0.46 | 0.07 |
| 5 | $\log_{10}$(*vol*) ~ (1 + $\log_{10}(r_{stem})$ \| species) + (1 + $\log_{10}(r_{stem})$ \| site) + $\log_{10}(r_{stem})$ | 0.15 | 0.06 | 0.00 | 0.21 | 0.10 | 0.48 | 0.06 |
| Deviance from mean scaling models, and variance composition of rel crown depth | | | | | | | | |

Table S 8. LMMs for crown dimensions including Biogeography and Ecosystem Type. 95% Wald confidence intervals in parentheses.

| | *Dependent variable:* | | | | | | | | | |
|---|---|---|---|---|---|---|---|---|---|---|
| | log(Crown Width) | | log(Depth) | | log(Relative Depth) | | log(Surface Area) | | log(Volume) | |
| | biogeog (1) | no biogeog (2) | biogeog (3) | no biogeog (4) | biogeog (5) | no biogeog (6) | biogeog (7) | no biogeog (8) | biogeog (9) | no biogeog (10) |
| eco_classTransitional | 0.272 | 0.28 | -0.223 | -0.25 | -0.267 | -0.342 | 0.741 | 0.727 | 1.197 | 1.227 |
| | (-0.383, 0.928) | (-0.342, 0.903) | (-0.762, 0.315) | (-0.751, 0.251) | (-1.068, 0.535) | (-1.106, 0.423) | (-0.522, 2.004) | (-0.456, 1.910) | (-0.716, 3.111) | (-0.537, 2.991) |
| eco_classTMCF | 0.355 | 0.496$^{**}$ | -0.055 | 0.229$^{**}$ | 0.05 | -0.089 | 0.712 | 1.058$^{***}$ | 1.239 | 1.642$^{***}$ |
| | (-0.241, 0.952) | (0.119, 0.874) | (-0.493, 0.383) | (0.028, 0.430) | (-0.596, 0.695) | (-0.469, 0.292) | (-0.430, 1.853) | (0.329, 1.787) | (-0.491, 2.968) | (0.557, 2.726) |
| eco_classTF | 0.188 | 0.304$^{*}$ | 0.107 | 0.329$^{***}$ | 0.264 | 0.174 | 0.553 | 0.797$^{**}$ | 0.983 | 1.247$^{**}$ |
| | (-0.309, 0.685) | (-0.040, 0.649) | (-0.238, 0.452) | (0.147, 0.510) | (-0.261, 0.789) | (-0.171, 0.518) | (-0.388, 1.493) | (0.135, 1.459) | (-0.443, 2.409) | (0.262, 2.232) |
| $\log(r_{stem})$ | 0.887$^{***}$ | 0.914$^{***}$ | | | -0.446$^{**}$ | -0.541$^{**}$ | 1.800$^{***}$ | 1.818$^{***}$ | 2.779$^{***}$ | 2.856$^{***}$ |
| | (0.630, | (0.684, | | | (-0.884, | (-0.967, | (1.309, | (1.386, | (2.032, | (2.208, |

| | | | | | | | | | | |
|---|---|---|---|---|---|---|---|---|---|---|
| | 1.144) | 1.143) | | | -0.008) | -0.114) | 2.291) | 2.249) | 3.526) | 3.504) |
| log($h_{tree}$) | | | 1.154*** (0.990, 1.318) | 1.090*** (0.939, 1.242) | 0.724*** (0.364, 1.085) | 0.725*** (0.363, 1.088) | | | | |
| regionGhana | 0.093 (-0.395, 0.581) | | 0.262 (-0.083, 0.606) | | 0.008 (-0.522, 0.538) | | 0.215 (-0.706, 1.137) | | 0.166 (-1.232, 1.564) | |
| regionPeru | 0.137 (-0.385, 0.659) | | 0.314 (-0.086, 0.713) | | -0.069 (-0.639, 0.501) | | 0.361 (-0.635, 1.358) | | 0.375 (-1.136, 1.885) | |
| eco_classTransitional:log($r_{stem}$) | -0.234 (-0.757, 0.288) | -0.261 (-0.757, 0.235) | | | -0.088 (-0.954, 0.778) | 0.007 (-0.850, 0.864) | -0.517 (-1.487, 0.454) | -0.535 (-1.445, 0.375) | -0.835 (-2.315, 0.644) | -0.912 (-2.284, 0.460) |
| eco_classTMCF:log($r_{stem}$) | -0.259 (-0.691, 0.172) | -0.315** (-0.594, -0.036) | | | 0.19 (-0.450, 0.830) | 0.152 (-0.354, 0.658) | -0.493 (-1.309, 0.323) | -0.725*** (-1.253, -0.197) | -0.899 (-2.142, 0.344) | -1.186*** (-1.978, -0.393) |
| eco_classTF:log($r_{stem}$) | -0.146 (-0.506, 0.214) | -0.203 (-0.458, 0.053) | | | 0.159 (-0.406, 0.723) | 0.192 (-0.283, 0.666) | -0.348 (-1.022, 0.325) | -0.506** (-0.987, -0.025) | -0.641 (-1.667, 0.384) | -0.811** (-1.534, -0.089) |

| | | | | | | | |
|---|---|---|---|---|---|---|---|
| log($r_{stem}$):regionGhana | -0.028 (-0.383, 0.328) | | | -0.085 (-0.511, 0.341) | | -0.102 (-0.764, 0.561) | -0.017 (-1.027, 0.993) |
| log($r_{stem}$):regionPeru | -0.033 (-0.414, 0.347) | | | -0.124 (-0.583, 0.335) | | -0.214 (-0.930, 0.502) | -0.212 (-1.303, 0.879) |
| eco_classTransitional:log($h_{tree}$) | | 0.224 (-0.277, 0.724) | 0.288 (-0.212, 0.787) | 0.427 (-0.483, 1.336) | 0.426 (-0.484, 1.336) | | |
| eco_classTMCF:log($h_{tree}$) | | 0.007 (-0.336, 0.349) | -0.332*** (-0.536, -0.127) | -0.298 (-0.744, 0.149) | -0.292 (-0.741, 0.156) | | |
| eco_classTF:log($h_{tree}$) | | -0.149 (-0.425, 0.127) | -0.395*** (-0.574, -0.217) | -0.380* (-0.797, 0.036) | -0.424** (-0.842, -0.005) | | |
| log($h_{tree}$):regionGhana | | -0.291** (-0.565, -0.016) | | | | | |
| log($h_{tree}$):regionPer | | -0.407** | | | | | |

| | | | | | | | | | | |
|---|---|---|---|---|---|---|---|---|---|---|
| u | | | | | | | | | | |
| | | | (-0.718, -0.097) | | | | | | | |
| Constant | $-0.534^{***}$ | $-0.542^{***}$ | $-0.363^{***}$ | $-0.336^{***}$ | 0.052 | 0.127 | -0.083 | -0.069 | $-1.428^{***}$ | $-1.458^{***}$ |
| | (-0.870, -0.197) | (-0.848, -0.235) | (-0.538, -0.187) | (-0.469, -0.203) | (-0.269, 0.373) | (-0.161, 0.415) | (-0.743, 0.576) | (-0.658, 0.521) | (-2.426, -0.430) | (-2.335, -0.581) |
| Observations | 1032 | 1032 | 1032 | 1032 | 1032 | 1032 | 1032 | 1032 | 1032 | 1032 |
| Log Likelihood | 422.306 | 426.696 | 413.816 | 413.179 | 118.826 | 118.733 | -80.959 | -77.031 | -516.167 | -514.43 |
| Akaike Inf. Crit. | -812.613 | -829.392 | -795.632 | -802.358 | -191.652 | -199.467 | 193.918 | 178.063 | 1064.335 | 1052.86 |
| Bayesian Inf. Crit. | -733.585 | -770.121 | -716.604 | -743.087 | -78.05 | -105.621 | 272.946 | 237.334 | 1143.363 | 1112.131 |
| *Note:* | | | | | | | | | | *p<0.1; **p<0.05; ***p<0.01 |

Table S 9. Crown depth models with precipitation covariates based on Model 17. Standard errors in parentheses. Significance estimated by Wald tests; our discussions are based on the significance estimates from the preferred Kenward-Roger approximations in Table S 10.

| | *Dependent variable:* | | | | |
|---|---|---|---|---|---|
| | log10(depth) | | | | |
| | Ghanaian transect | | | | all sites |
| | $r_{stem}$ + h | h | $r_{stem}$ | precip only | $r_{stem}$ + h |
| | (1) | (2) | (3) | (4) | (5) |
| precip | 0.167 | 0.100 | 0.380** | -0.020 | 0.060 |
| | (0.155) | (0.112) | (0.191) | (0.047) | (0.068) |
| log(h) | 0.516*** | 0.733*** | | | 0.670*** |
| | (0.096) | (0.070) | | | (0.053) |
| log($r_{stem}$) | 0.245** | | 0.558*** | | 0.161*** |
| | (0.109) | | (0.134) | | (0.046) |
| precip:log(h) | 0.094 | -0.097 | | | -0.116* |
| | (0.100) | (0.075) | | | (0.060) |
| precip:log($r_{stem}$) | -0.226** | | -0.264** | | 0.022 |
| | (0.111) | | (0.134) | | (0.047) |
| Constant | -0.039 | -0.005 | 0.247 | 0.904*** | -0.159** |
| | (0.153) | (0.107) | (0.190) | (0.047) | (0.062) |
| Observations | 228 | 228 | 228 | 228 | 1,032 |
| Log Likelihood | 110.422 | 104.858 | 96.010 | 98.454 | 443.602 |
| Akaike Inf. Crit. | -182.843 | -187.717 | -170.020 | -178.908 | -849.204 |
| Bayesian Inf. Crit. | -117.685 | -149.994 | -132.297 | -148.044 | -755.358 |
| *Note:* | *p<0.1; **p<0.05; ***p<0.01 | | | | |

Table S 10. ANOVA table of precipitation models (Model 17, Table S 9 Models 1 and 5). Significance level estimated via Kenward-Roger approximation.

| **Effect** | **df** | **F** | **p.value** |
|---|---|---|---|

| | | | |
|---|---|---|---|
| | **Ghana Only** | | |
| precip | 1, 4.86 | 1.00 | .37 |
| log(h) | 1, 5.41 | 20.40 ** | .005 |
| log($r_{stem}$) | 1, 5.55 | 4.30 + | .09 |
| precip:log(h) | 1, 7.62 | 0.66 | .44 |
| precip:log($r_{stem}$) | 1, 5.86 | 3.49 | .11 |
| | **All Sites** | | |
| precip | 1, 19.96 | 0.75 | .40 |
| log(h) | 1, 22.21 | 138.24 *** | <.0001 |
| log($r_{stem}$) | 1, 22.14 | 11.37 ** | .003 |
| precip:log(h) | 1, 25.73 | 3.47 + | .07 |
| precip:log($r_{stem}$) | 1, 15.26 | 0.20 | .66 |
